# RecA status determines SOS- and RecBCD-dependent outcomes in CRISPR–Cas adaptation

**DOI:** 10.64898/2026.08.20.745894

**Authors:** Harry Edwards, Christopher Cannon, Jack Braithwaite, Ronald Chalmers

## Abstract

CRISPR–Cas immunity depends on integrating DNA fragments from invading elements, yet how this process is tuned by host physiology remains poorly understood. Here, we investigate the relationship between CRISPR adaptation and the bacterial SOS DNA damage response in *Escherichia coli* using a highly sensitive colony-based adaptation assay. Blocking SOS with a *lexA3* allele suppresses adaptation, while paradoxically, deleting *recA* enhances it. These findings indicate that RecA contributes positively to adaptation indirectly, through LexA cleavage and induction of SOS-regulated functions, while complete loss of RecA is associated with a RecBCD-dependent DNA-processing state that favours successful adaptation. The RecA inhibitors RecX and PsiB did not phenocopy Δ*recA*, indicating that inhibitor-mediated perturbation of RecA in RecA-proficient cells and complete loss of RecA have different consequences for adaptation. Genome-wide maps of recovered spacers show that RecA and LexA reshape the relative distribution of chromosomal and plasmid-derived spacers, linking CRISPR adaptation to replication and DNA repair dynamics. Although Cas1–Cas2 catalyses spacer integration autonomously in vitro, our findings suggest that successful adaptation in vivo emerges from interactions between spacer acquisition, DNA repair, and bacterial stress physiology, embedding CRISPR adaptation within cellular networks that balance immune protection with the risk of autoimmunity.

**Author summary:** Bacteria can protect themselves from viruses using CRISPR systems, which store fragments of viral DNA as a genetic memory of past infections. However, this process is dangerous because the same machinery can accidentally capture fragments of the bacterium’s own DNA, potentially causing self-destruction. How bacteria balance these risks is still poorly understood.

In this study, we examined how CRISPR activity is linked to bacterial DNA damage responses using the model organism *Escherichia coli*. We used a sensitive colony-based assay that allows rare events in which bacteria acquire new CRISPR memories to be seen directly as small blue outgrowths within bacterial colonies. We found that a major stress-response pathway helps this process succeed, but that deleting one of its central proteins, RecA, unexpectedly increased adaptation.

We also found that stress-response genes influence which parts of the genome are used as sources of new CRISPR memories. Together, our results show that CRISPR immunity is closely connected to broader stress and DNA repair systems inside the cell, rather than operating as an isolated immune mechanism.

## Introduction

Bacteria face persistent threats from mobile genetic elements, including plasmids, transposons, and bacteriophages. While some elements confer selective advantages, others, particularly lytic phages, pose a lethal threat. This pressure has driven the emergence of diverse defense mechanisms, among which CRISPR–Cas provides a form of adaptive immunity [1]. Clarifying how these systems are deployed and regulated is central to understanding bacterial survival strategies.

CRISPR–Cas immunity operates by integrating short DNA fragments, called spacers, from foreign elements into the host genome. These spacers are later transcribed and used to guide sequence-specific interference against future invasions [2–4]. In contrast to the diversity of interference strategies across bacterial and archaeal lineages, the adaptation phase is universally dependent on the Cas1–Cas2 complex [5–8].

Because Cas1–Cas2 does not inherently distinguish host from foreign DNA, spacer acquisition carries a substantial risk of autoimmunity. This cost creates balancing selection on the rate of new spacer acquisition: increasing protection against mobile genetic elements comes at the expense of host viability. Mathematical modeling predicts a ‘safe zone’ where protective benefits outweigh the cost of autoimmunity [9]. CRISPR–Cas systems therefore appear tuned to a regime where acquisition is rare, producing occasional ‘jackpot’ survivors that can repopulate after phage outbreaks. These constraints suggest that the rate of adaptation should be conditioned by cellular physiology, modulated by regulatory networks that balance immune protection with the risk of autoimmunity [10].

Although Cas1–Cas2 catalyses spacer integration autonomously *in vitro*, successful adaptation *in vivo* may nevertheless depend on host DNA repair and coordination functions. Integration generates unusual DNA intermediates at the CRISPR locus, including partially integrated or single-ended products, which may require processing, maturation, or resolution by host factors before stable inheritance can occur. Consistent with this, in the absence of interference CRISPR loci themselves appear as hotspots in protospacer maps, an unexpected finding given the rarity of adaptation events [11, 12].

Importantly, rare adaptive events may not scale linearly with population-average measurements of gene expression or stress. Instead, spacer acquisition is likely governed by thresholded physiological states that arise transiently in a subset of cells, with successful outcomes reflecting the survival and expansion of these rare lineages rather than the mean behavior of the population, particularly under structured growth conditions where cellular physiology is strongly heterogeneous [11].

Several studies provide evidence for such regulation. In *E. coli*, we recently reported that growth on solid medium, more representative of structured environmental conditions, reduces the strong plasmid bias of spacer acquisition observed in liquid culture [11]. Spacer source bias also depends on Cas1–Cas2 expression level [12]. Furthermore, expression of the *cas* genes is repressed by the nucleoid-associated protein H-NS and relieved by LeuO [13, 14], and may also respond to envelope stress via the BaeSR two-component system [15, 16]. In *Streptococcus pyogenes*, *cas* gene expression is activated heterogeneously in a subset of cells, indicating that CRISPR– Cas regulation can also operate at the single-cell level [17]. These examples suggest that CRISPR–Cas immunity is embedded within broader stress-response circuits rather than operating in isolation.

Here, we examine how CRISPR–Cas adaptation interfaces with the SOS DNA damage response in *E. coli*, as illustrated schematically in Fig 1. We show that Cas1–Cas2 overexpression induces filamentation via RecA-mediated SOS activation. Adaptation is reduced in *lexA3* cells, implicating SOS-regulated functions in spacer acquisition. Paradoxically, deletion of *recA* increases adaptation, whereas expression of RecA inhibitors reduces it. These findings suggest that RecA-dependent LexA cleavage and induction of the SOS regulon support productive adaptation, while complete loss of RecA reveals a genetically separable, RecBCD-dependent DNA-processing state that can enhance adaptation. Genome-wide spacer maps further reveal that RecA influences the balance between chromosomal and plasmid acquisition, providing a potential clue to the mechanisms underlying plasmid bias in the Type I-E system. Moreover, by linking adaptation to RecA, SOS and RecBCD activities, which are targeted by phage and plasmid inhibitors, our results raise the possibility that mobile elements can influence spacer acquisition indirectly by perturbing host DNA-damage and repair physiology. Taken together, our study places CRISPR–Cas adaptation within the bacterial stress-response network, where host physiology shapes the conditions under which spacer acquisition becomes productive.

**Fig 1.**
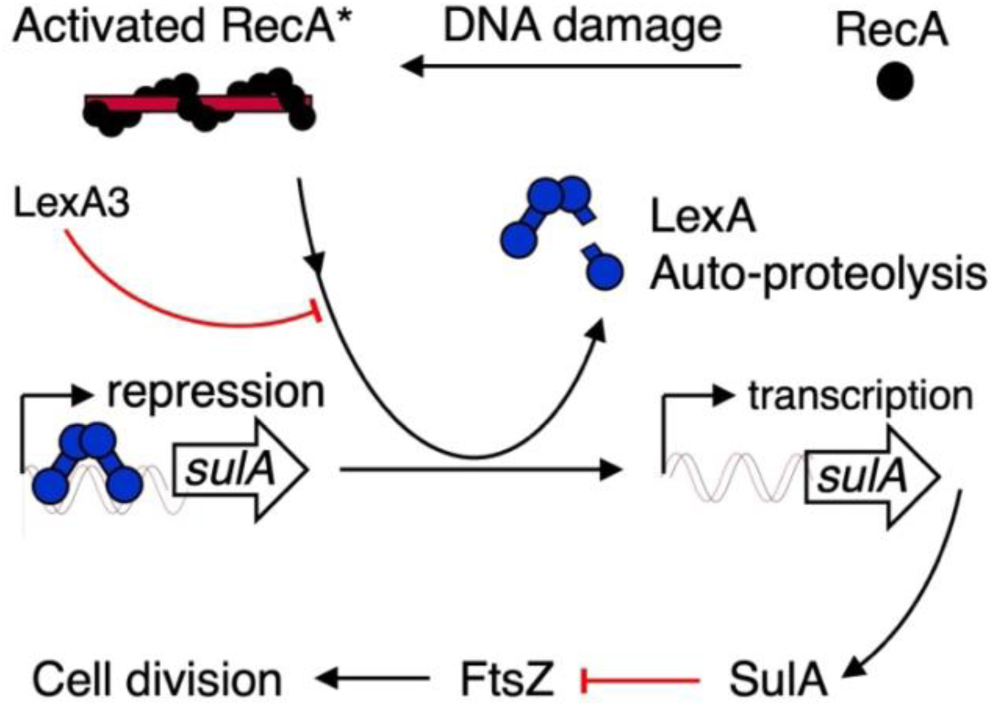
The recA-mediated response to DNA damage in *E. coli*. RecA is activated upon polymerization on ssDNA intermediates formed by DNA damage. Activated RecA promotes the autoproteolysis of LexA, a dimeric helix-turn-helix repressor. LexA depletion de-represses the LexA regulon, including the archetypal *sulA* gene. SulA inhibits cell division by preventing FtsZ polymerization at the mid-cell. Once DNA repair is complete, LexA levels are restored, repression of the regulon resumes, and cell division proceeds.

## Results

### Cas1–Cas2 Expression Triggers SOS-Dependent Filamentation

A previous study reported that expression of *cas2* or *cas1–cas2* from a T7 promoter causes filamentation in *E. coli* BL21, and that this is linked to elevated CRISPR–Cas adaptation [18]. To reproduce this result and to test whether this effect occurs under more moderate expression conditions, we expressed *cas1–cas2* from the arabinose-inducible vector pBAD and grew cells overnight in LB medium with 0.2% arabinose. Phase-contrast microscopy revealed a significant increase in the number of filamentous cells compared to uninduced controls (Fig 2A, top middle panel; S3 Table).

**Fig 2.**
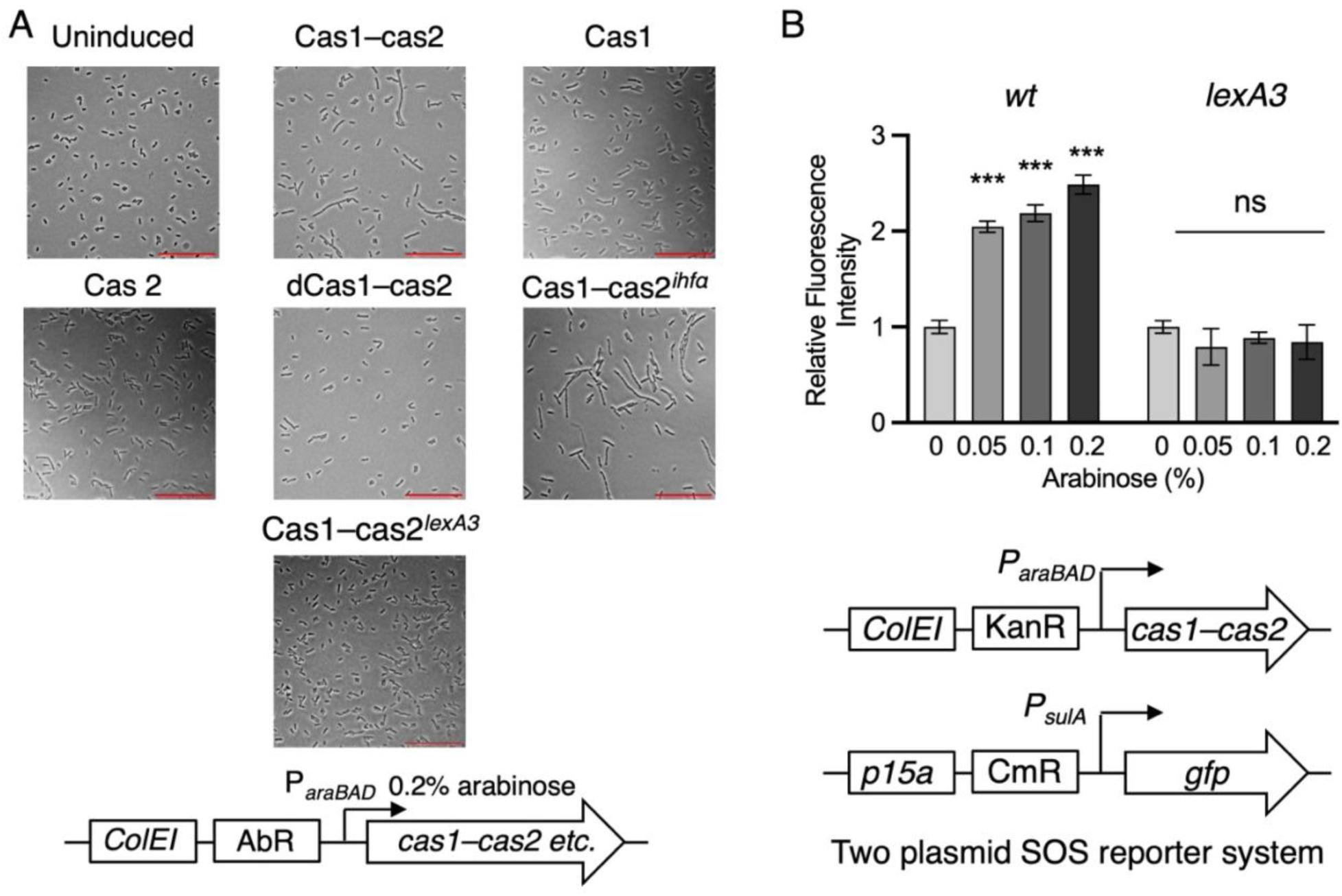
Cas1–Cas2 activity induces SOS-dependent filamentation via DNA damage. A. Cells were transformed with pBAD expression vectors encoding the indicated proteins and grown overnight in LB medium supplemented with 0.2% arabinose to induce expression. Phase-contrast images were captured at 630× magnification. Plasmids used were: Cas1–Cas2 (pRC1656 [kanamycin] or pRC1657 [chloramphenicol]), Cas1 (pRC1653), Cas2 (pRC1655), and catalytically inactive dCas1(D221A)-Cas2 (pRC1658). The host strain was *E. coli* BL21-AI. The isogenic Δ*ihfα* and *lexA3* derivatives were strains RC5362 and RC5250, respectively. Deletion of *ihfα* does not suppress Cas1–Cas2-dependent filamentation. Cells in the representative fields were categorized by morphology using consistent visual criteria across conditions, as summarized in S3 Table (normal rods, moderately elongated cells, strong filaments, extreme filaments, and ambiguous/dividing cells). Red scale bar is 20 µm. B. Cells were co-transformed with the Cas1–Cas2 expression vector (pRC1656) and an SOS reporter plasmid (pRC2790). Cultures were grown overnight with the indicated concentrations of arabinose. Wild-type and *lexA3* derivatives were as described in (A). GFP fluorescence was measured in triplicate; bars show the mean ± SEM. Conditions were compared using a two-sample *t*-test. P values: ***< 0.001; ns, not significant.

Filamentation required co-expression of *cas1* and *cas2*, as neither gene alone produced the phenotype; cells expressing the catalytically inactive complex (dCas1[D221A]– Cas2) remained morphologically normal. These findings indicate that Cas1 catalytic activity within the complex was required for filamentation.

Filamentation did not require Integration Host Factor (IHF), an essential cofactor for spacer integration into the CRISPR array [19]. Cells lacking IHF still underwent Cas1– Cas2-induced filamentation, suggesting that the phenotype does not depend only on canonical spacer integration but may arise from off-target full- or half-site integration events [20, 21].

To determine whether the filamentation reflects SOS activation, we repeated the experiment in an isogenic *lexA3* mutant background. This allele encodes a non-cleavable LexA repressor that blocks induction of the SOS regulon and inhibition of cell division [22–24]. In *lexA3* cells, Cas1–Cas2 expression failed to induce filamentation (Fig 2A, bottom panel; S3 Table), indicating that the phenotype requires SOS activation. To determine whether Cas1–Cas2–dependent filamentation reflects activation of the SOS response, we used a P*_sulA_–gfp* reporter assay. This assay is included to confirm that the filamentation phenotypes observed in Fig 2A arise from SOS induction, rather than to provide a systematic survey of SOS activity across all strains shown. Cas1– Cas2 expression led to a significant increase in fluorescence in wild-type cells, but not in the *lexA3* mutant, consistent with RecA-dependent LexA cleavage.

Together, these results demonstrate that expression of catalytically active Cas1–Cas2 generates DNA damage sufficient to activate the SOS response and induce filamentation. This occurs independently of spacer integration at the CRISPR loci and is suppressed when the SOS pathway is blocked.

### SOS Response and Cell Cycle Regulation Promote CRISPR–Cas Adaptation

To monitor CRISPR–Cas adaptation, we used a previously developed papillation assay in which newly acquired spacers restore the reading frame of a disrupted *lacZ* reporter gene [11]. Adapted cells switch from *lac⁻* to *lac⁺*, producing blue papillae on lactose/X-gal agar (S1 Fig, S1 File). This system provides a visual readout of successful adaptation, while papilla counting allows semi-quantitative comparison between conditions (S1 Fig). We typically use the P*_araBAD_* promoter (pBAD) when testing conditions expected to reduce adaptation. In contrast, the weaker P*_araMari_* promoter is preferable when the experimental variable may enhance adaptation, because its lower baseline increases sensitivity to gains in adaptation frequency (S1 Fig) [11, 25]. In all papillation experiments, Cas1–Cas2 expression was induced at levels substantially lower than those used to elicit filamentation in microscopy assays (Fig 2A), allowing adaptation to be analyzed under conditions that do not impose overt cell-cycle arrest or toxicity. Under these *P_araBAD_* conditions, SOS induction was barely detectable (S2 Fig). Cas1–Cas2 expression from the weaker *P_araMari_* promoter is approximately 40-fold lower than from *P_araBAD_* [11], and is therefore expected to produce negligible SOS induction.

The endogenous *cas* genes of *E. coli* K-12 remain strongly repressed by H-NS under standard laboratory conditions, while *E. coli* B strains such as BL21 lack the *cas* genes entirely. To test whether SOS-regulated functions are required for adaptation, we compared wild-type and *lexA3* strains (Fig 3A). Introducing the non-cleavable *lexA3* allele into the reporter strain led to a marked reduction in papillae formation relative to wild type (Fig 3A), indicating that SOS-regulated functions support successful CRISPR–Cas adaptation. This genetic requirement for SOS persists under expression conditions where Cas1–Cas2 does not measurably induce SOS on its own, indicating that SOS supports the success of adaptation events rather than simply mitigating Cas1–Cas2-induced toxicity.

**Fig 3.**
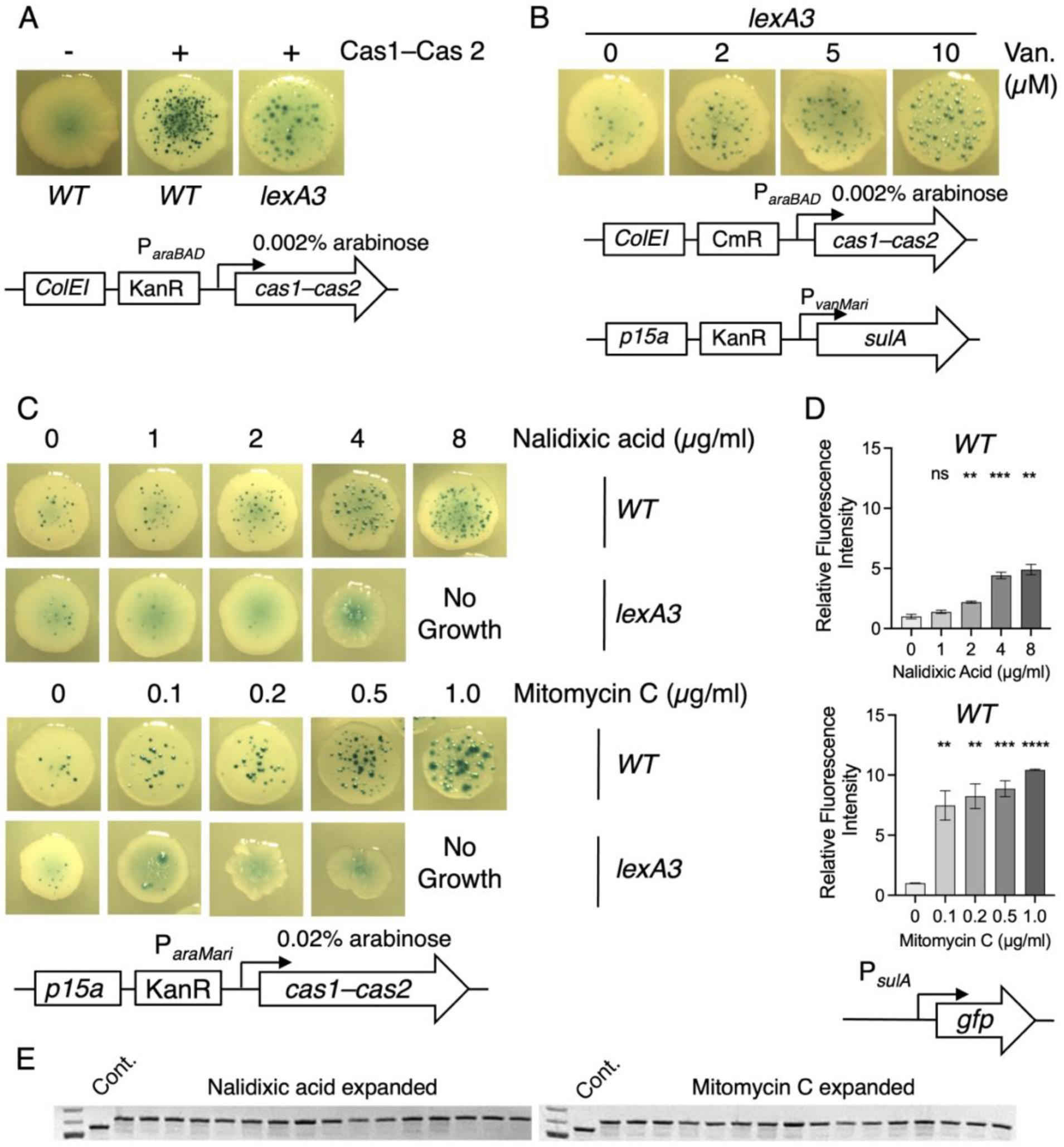
SOS-regulated physiology promotes CRISPR adaptation. Activation of the SOS response enhances spacer acquisition, while its repression reduces it. A. Papillation was assessed in the wild-type CRISPR–Cas adaptation reporter strain (RC5311) and an isogenic *lexA3* derivative (RC5353). Cas1–Cas2 was expressed from plasmid pRC1656. B. The *lexA3* strain (RC5355) was co-transformed with protein expression vectors for SulA (pRC2781) and Cas1–Cas2 (pRC1657). SulA expression was titrated using the indicated vanillate concentrations. Partial complementation of the *lexA3* phenotype was observed. Full complementation was unlikely because the exogenous promoter is not subject to the regulatory feedback of the natural promoter. C. Papillation assays were performed in the presence of the indicated concentrations of nalidixic acid and mitomycin C. Cas1–Cas2 was expressed from the P*_araMari_* promoter (pRC2747). The wild-type and isogenic *lexA3* strains were RC5311 and RC5353, respectively. D. SOS induction in response to nalidixic acid and mitomycin C was quantified using a GFP-based SOS reporter plasmid (pRC2790) in the wild-type reporter strain (RC5311). About 50 colonies were grown overnight on LB agar plates supplemented with the antibiotic concentration shown in the figure. Colonies were resuspended in LB broth and OD_600_ measurements were equalized by dilution with fresh LB. Expression of the sulA– GFP reporter is fully dependent on LexA cleavage; accordingly, no GFP induction is observed in the *lexA3* background under any of the conditions tested (data not shown). Fluorescence measurements were performed in triplicate; bars represent the mean ± SEM. Conditions were compared using a two-sample *t*-test. P values: **, <0.01; ***, <0.001; ****, <0.0001, ns, not significant. E. Individual papillae were isolated from plates treated with nalidixic acid or mitomycin C across the tested concentration range. To eliminate background lacZ⁻ cells, each papilla was re-streaked for single colonies on LB agar supplemented with X-gal. PCR amplification across the leader-proximal end of the chromosomal CRISPR array yields an upward band shift following integration of a new repeat-spacer unit.

SulA, an SOS-regulated inhibitor of cell division, blocks FtsZ polymerization and thereby causes filamentation. To determine whether SulA could rescue adaptation in the *lexA3* background, we expressed *sulA* under an inducible promoter. Papillae formation increased in a dose-dependent manner (Fig 3B), though it did not reach wild-type levels. This partial rescue suggests that cell cycle delay supports adaptation, but dynamic regulation of SulA, specifically, the transient arrest and timely release of division, may be critical. Moreover, additional SOS-regulated factors likely contribute.

This led us to test whether global activation of the SOS regulon could enhance adaptation. To determine whether the SOS response directly facilitates adaptation or simply mitigates Cas1–Cas2-induced stress, we used the P*_araMari_* expression vector (introduced above). This system lowers Cas1–Cas2 expression, thereby increasing sensitivity to factors that enhance adaptation. To test whether activating the SOS response can promote adaptation independently of Cas1–Cas2 induced stress, we treated cells with nalidixic acid, a DNA gyrase inhibitor that induces SOS via replication fork collapse. Nalidixic acid increased papillae formation in wild-type cells (Fig 3C), correlating with SOS activation as measured by a sulA–gfp reporter (Fig 3D). Similar results were obtained with mitomycin C, a DNA crosslinking agent. Crucially, neither drug increased adaptation in the *lexA3* background, despite continued exposure to DNA-damaging conditions. This finding argues that DNA damage alone is insufficient to explain enhanced adaptation and instead supports a requirement for SOS-regulated functions.

To confirm that the papillation phenotype reflected bona fide CRISPR adaptation rather than a reporter-specific event, we amplified the leader-proximal region of the endogenous CRISPR array from 15 independent papillae isolated from nalidixic acid- and mitomycin C-treated plates (Fig 3E). Expanded PCR products showed the expected size increase associated with integration of a new repeat-spacer unit, confirming bona fide spacer acquisition at the chromosomal CRISPR array. Together, these results show that the SOS response supports CRISPR–Cas adaptation as part of a permissive physiological state. This effect is amplified under genotoxic stress but requires SOS induction.

We also tested whether constitutive SOS derepression was sufficient to increase adaptation using a *lexA51 sulA* double mutant (S3 Fig). Papillation was not increased relative to wild type, indicating that chronic derepression of the SOS regulon is not sufficient to mimic the adaptation-promoting state observed after genotoxic stress. This result is not unexpected, because constitutive SOS derepression removes normal temporal regulation and requires deletion of *sulA*, a major SOS-regulated cell-division inhibitor. We note additionally that constitutive SOS derepression strongly elevates RecA levels, which may itself influence adaptation outcomes independently of other SOS functions (see below). We therefore interpret the SOS effect as dependent on a coordinated physiological state rather than on uniform elevation of the SOS regulon.

### RecA Loss Enhances CRISPR–Cas Adaptation

As adaptation is reduced in the *lexA3* background (Fig 3A), we next tested the effect of deleting *recA*. Using the papillation assay, we compared wild type and Δ*recA* under P_araMari_ and P_araBAD_ expression (Fig 4A). Surprisingly, the Δ*recA* mutant produced more papillae than wild type, despite lacking the ability to cleave LexA and activate the SOS response. This contrasts with *lexA3*, where blocking SOS reduces adaptation, indicating that the Δ*recA* phenotype cannot be explained simply by loss of SOS induction.

**Fig 4.**
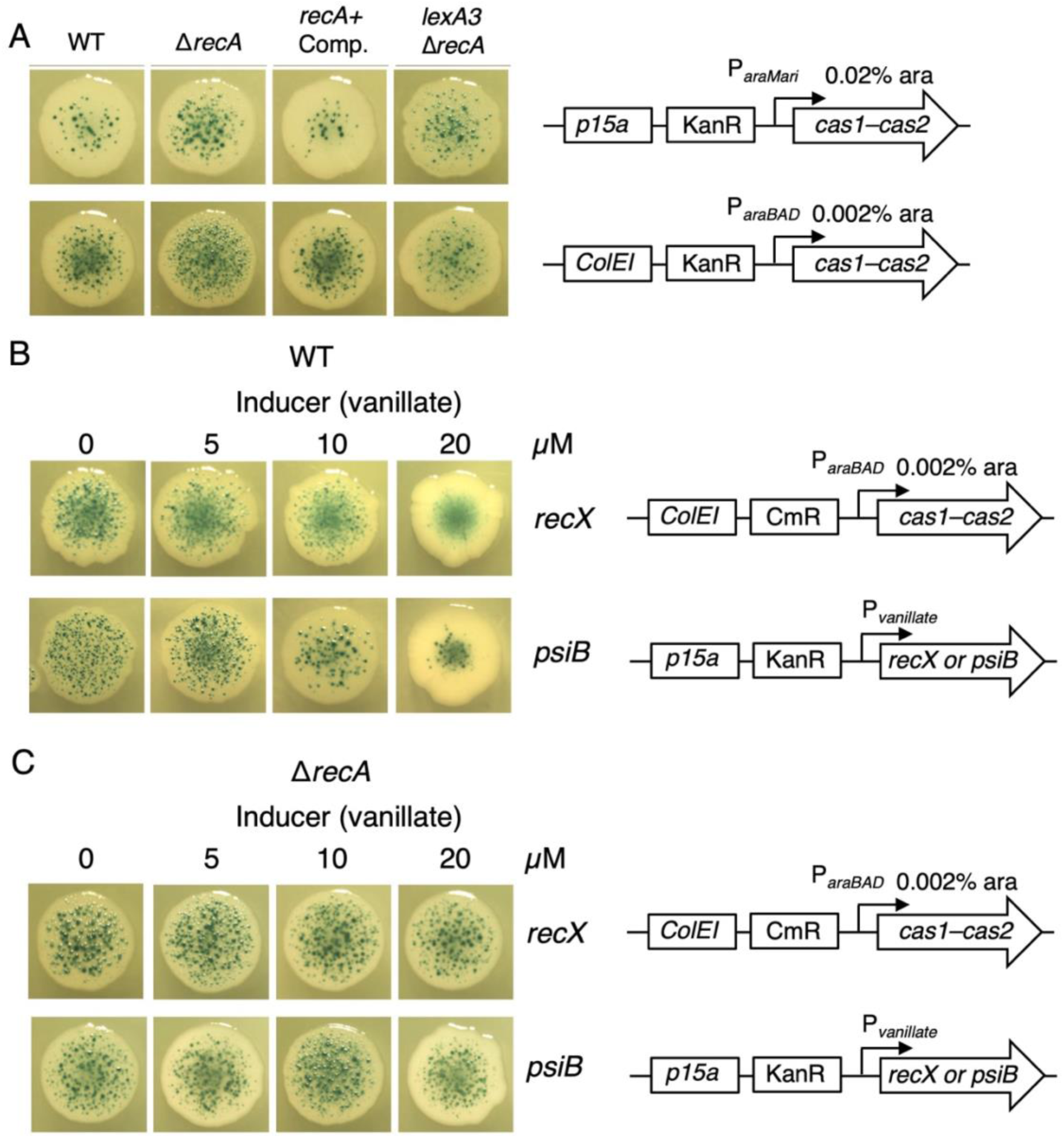
RecA loss enhances CRISPR adaptation, whereas RecA antagonism in RecA-proficient cells reduces adaptation. A. CRISPR–Cas adaptation reporter strains were transformed with plasmids expressing Cas1–Cas2 under either the weak P*_araMari_* promoter or the strong P*_araBAD_* promoter. Adaptation was unexpectedly elevated in the Δ*recA* strain, so complementation was performed by restoring the wild-type *recA* allele to its native locus using λ Red recombination (denoted Δ*recA::recA⁺*, where “::” indicates insertion of the wild-type allele at the deleted locus). Strains: wild-type CRISPR–Cas adaptation reporter (RC5311), Δ*recA* (RC5356), Δ*recA::recA⁺*, (RC5357), and *lexA3* Δ*recA* (RC5360). Plasmids: pRC2747 (P*_araMari_*), pRC1656 (P*_araBAD_*). B. The wild-type reporter strain (RC5311) was transformed with plasmids expressing Cas1–Cas2 (pRC1657), RecX (pRC3017), or PsiB (pRC3028), as indicated. RecX and PsiB expression was titrated with the specified concentrations of vanillic acid. C. As in part B, but using the Δ*recA* derivative (RC5356) to test whether RecX and PsiB effects require RecA.

Since this result was unexpected, we performed additional controls and quantitative assays to confirm the effect of *recA* deletion. First, to rule out secondary mutations introduced during P1 transduction, we restored the Δ*recA* allele to wild type using λ-Red recombination. Adaptation returned to wild-type levels, confirming that the phenotype resulted specifically from the loss of RecA (Fig 4A, third panel). Second, we quantified the papillation phenotype directly by counting papillae in 10 colonies each of wild-type and Δ*recA* strains. This analysis showed a 3.4-fold increase in papillation in the ΔrecA strain, confirming the increase observed by visual assessment (Fig 5). Third, we transformed a yfp-based adaptation reporter strain [26] and its Δ*recA* derivative with the P*araMari* Cas1–Cas2 expression plasmid and quantified adapted cells by fluorescence-activated cell sorting after growth under mock papillation conditions. The number of adapted cells increased 4.7-fold in the Δ*recA* strain (S4 Fig). Thus, loss of RecA substantially enhances adaptation under structured-colony assay conditions.

**Fig 5.**
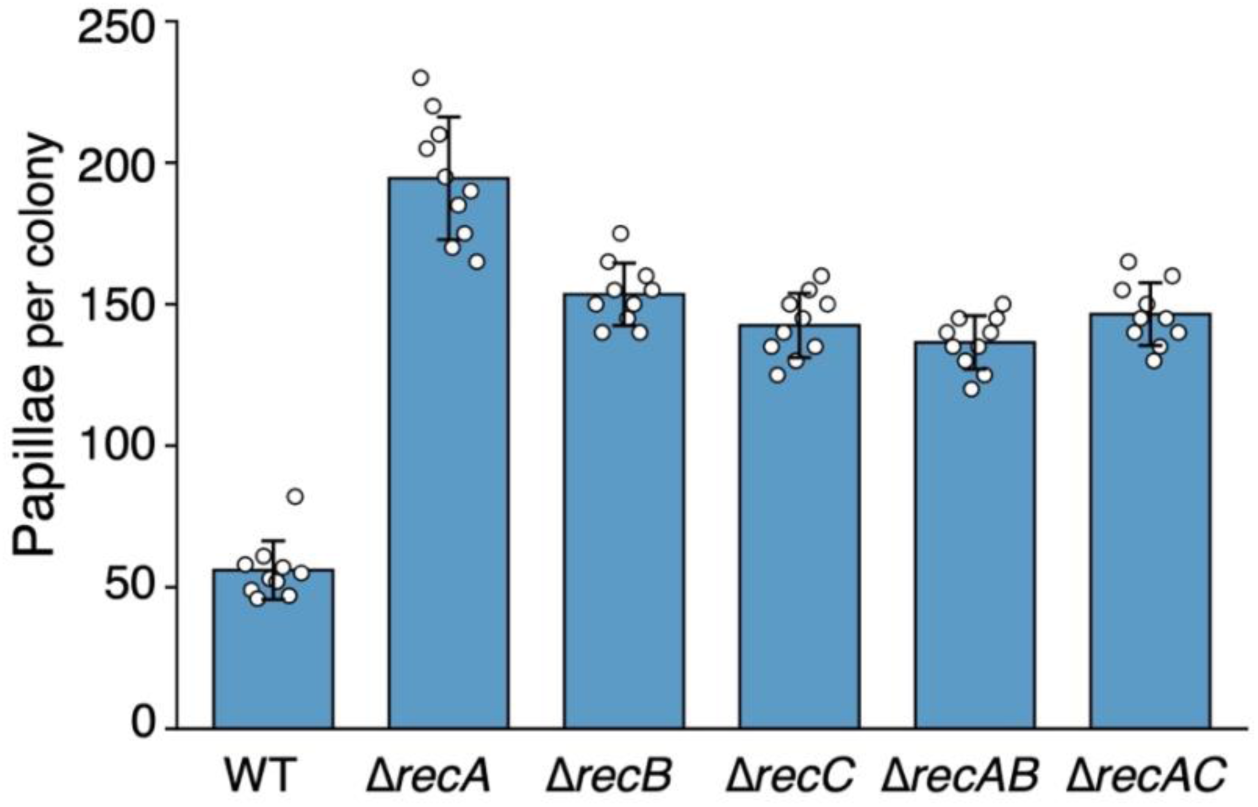
RecB and RecC contribute to the elevated adaptation phenotype of Δ*recA* cells. Papillation assays were performed using the wild-type reporter strain (RC5311) and the indicated Δ*recA* (RC5356), Δ*recB* (RC5368), Δ*recC* (RC5369), Δ*recA* Δ*recB* (RC5371), and Δ*recA* Δ*recC* (RC5372) derivatives transformed with the P_araMari_ Cas1–Cas2 expression plasmid. Cas1–Cas2 expression was induced with 0.02% arabinose. Papillae were counted from 10 colonies per genotype; bars show mean ± SD. Colonies used for quantification are shown in S5 Fig. Differences among genotypes were analysed by Welch’s one-way ANOVA (F5,25.1=123.7, P<0.0001), followed by Games– Howell multiple-comparisons tests. Papillation was increased in Δ*recA*, Δ*recB* and Δ*recC* relative to WT (all P<0.0001), and reduced relative to Δ*recA* in Δ*recA* Δ*recB* (P<0.0001) and Δ*recA* Δ*recC* (P=0.0003) relative to Δ*recA*.

To place this finding in the context of SOS regulation, we constructed a double mutant lacking both LexA cleavage and RecA (*lexA3* Δ*recA*). This strain exhibited fewer papillae than Δ*recA* alone, but markedly more than *lexA3*, indicating that the effect of RecA loss is not strictly coupled to SOS induction. Instead, the two effects overlap, indicating that RecA loss and SOS blockade make separable contributions to the adaptation phenotype under these assay conditions (Figs 3A and 4A).

We next asked whether inhibitor-mediated antagonism of RecA in otherwise RecA-proficient cells could mimic complete loss of RecA. To test this, we expressed two inhibitors: RecX, a host-encoded protein that caps RecA filaments and partially inhibits co-protease activity, and PsiB, an F plasmid protein that blocks RecA filament nucleation and strongly suppresses RecA-dependent responses. (Fig 4B) [27–30]. Both inhibitors reduced adaptation, with PsiB showing a stronger and more progressive effect. At high expression levels, both suppressed papillation more effectively than *lexA3* (compare Figs 3A and 4B). These results suggest that CRISPR adaptation is supported by RecA activity in RecA-proficient cells. Thus, RecA antagonism in otherwise RecA-proficient cells does not phenocopy complete RecA loss, consistent with the idea that the Δ*recA* phenotype reflects a distinct physiological state rather than simple loss of RecA filament function.

To confirm that RecX and PsiB act through RecA, we expressed both in a Δ*recA* background (Fig 4C). Neither inhibitor affected papillation in this context, confirming that their effects are RecA-dependent. This argues against nonspecific toxicity or direct inhibition of Cas1–Cas2 as explanations for the reduced papillation seen in RecA-proficient cells. It also indicates that inhibitor-expressing RecA-proficient cells are not equivalent to cells lacking RecA entirely.

Because RecBCD-dependent processing has been implicated in prespacer generation [12], we next tested whether the elevated papillation observed in Δ*recA* cells depends on RecB or RecC. We constructed Δ*recA* Δ*recB* and Δ*recA* Δ*recC* double mutants and quantified papillation using lower-level Cas1–Cas2 expression from the P_araMari_ promoter (Fig 5; S5 Fig). Under these conditions, Δ*recA* again produced a pronounced increase in papillation relative to wild type. Papillation was also elevated in Δ*recB* and Δ*recC* single mutants, in contrast to the reduced adaptation previously reported for RecBCD mutants in a liquid-culture assay using T7-driven Cas1–Cas2 overexpression [12]. The Δ*recA* Δ*recB* and Δ*recA* Δ*recC* double mutants remained elevated relative to wild type, but were reduced relative to Δ*recA* and resembled the corresponding Δ*recB* and Δ*recC* single mutants. Thus, loss of RecB or RecC suppresses the additional increase associated with loss of RecA, indicating that the full Δ*recA* hyperadaptation phenotype depends on RecBCD function. However, because RecBCD-deficient strains retained substantial papillation, these data do not support a model in which RecBCD activity is essential for spacer acquisition. Instead, RecA and RecBCD have interacting effects on the DNA-processing states that support successful adaptation.

### Spacer Distribution Is Rewired in Δ*recA* and *lexA3* Mutants

In *E. coli* BL21 and K12, chromosomally derived protospacers have been shown to cluster near *oriC*, *terA*, *terC*, and the CRISPR locus [11, 12]. To determine whether RecA and the SOS response affect spacer acquisition across the genome, we generated spacer maps for wild-type and mutant strains (Fig 6A, B). To do this, colonies were pooled, passaged in M9 lactose minimal medium to enrich adapted cells, and the CRISPR reporter locus was PCR-amplified and Illumina sequenced.

**Fig 6.**
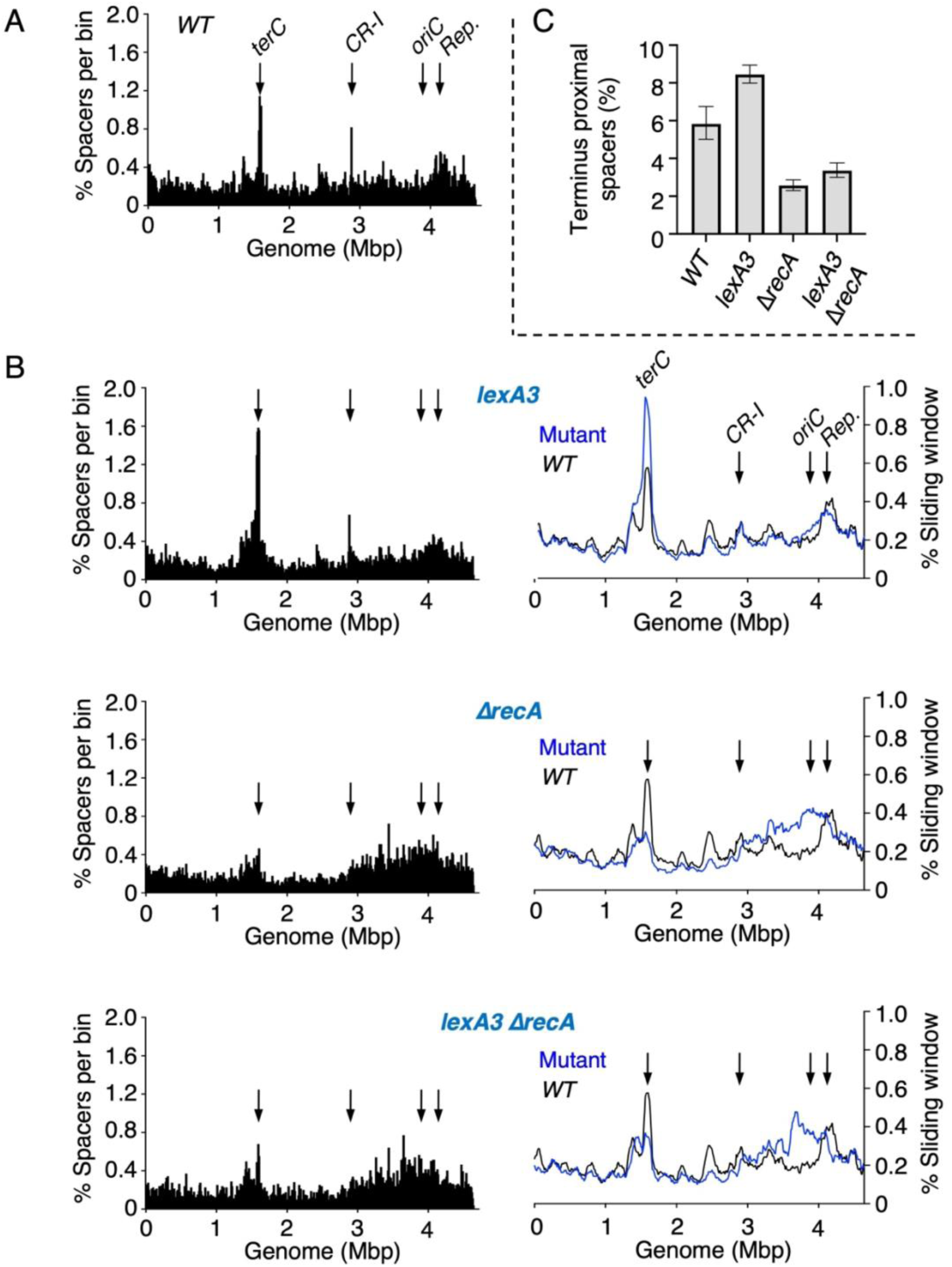
Chromosomal protospacer distribution maps for *lexA3* and Δ*recA* mutants. Colonies from papillation plates were pooled, enriched by one cycle of growth in M9 minimal lactose media, and the CRISPR reporter locus was PCR-amplified. Illumina amplicon sequencing was used to obtain acquired spacers, which were extracted and mapped onto the *E. coli* genome using custom Python scripts. Cas1–Cas2 was expressed from plasmid pRC1656 under control of the P*_araBAD_* promoter and induced with 0.002% arabinose. A. Protospacer counts were binned into 10 kb intervals and mapped across the chromosome of the wild-type CRISPR–Cas adaptation reporter strain (RC5311). B. Chromosomal protospacer distributions were also analyzed using a 100 kb sliding window with a step size of 10 kb. Profiles are shown for *lexA3* (RC5353), Δ*recA* (RC5356), and *lexA3* Δ*recA* (RC5360) and compared to wild type (RC5311). C. The percentage of total chromosomal protospacers mapping within a 100 kb region centered on the *dif* site is plotted for each strain. Error bars indicate 95% Wilson confidence intervals calculated from the underlying spacer counts. For terminus-proximal spacers, the denominator was total chromosomal spacers.

In wild-type cells, the protospacer distribution closely resembled previous maps generated via Oxford Nanopore sequencing [11], with clear hotspots near the chromosome terminus and origin, together with a sharp spike at the *CRISPR-I* locus (Fig 6A). Notably, the *CRISPR-I* spike was abolished in Δ*recA* but only partially reduced in *lexA3*, suggesting that this local signal is RecA-dependent but not simply a consequence of SOS induction (Fig 6B). A corresponding spike was not easily discernible at the reporter locus, likely because it lies within the broader origin-proximal enrichment and because the *CRISPR-II* locus was deleted in the reporter strain. More generally, in the *lexA3* mutant, terminus enrichment increased substantially, accompanied by a broad relative reduction in spacer acquisition across the genome except near the reporter.

By contrast, the Δ*recA* strain displayed a striking shift in the opposite direction: a marked increase in protospacers near *oriC* and a near-complete loss of enrichment at the terminus. The *lexA3* Δ*recA* double mutant resembled Δ*recA* alone, suggesting that RecA plays a dominant role in shaping protospacer distribution. This trend is quantified in Fig 6C, which shows the proportion of protospacers mapping to a 100 kb window centered on the *dif* site. Terminus enrichment increased by over 50% in *lexA3* and decreased by more than 50% in Δ*recA*; the double mutant showed only a slight recovery relative to Δ*recA*.

Plasmid protospacer distributions in wild-type and mutant strains mirrored the chromosomal trends (Fig 7A). In wild-type cells, we observed two pronounced peaks just downstream of the ColE1 initiation site. These peaks intensified in the *recA* deletion, consistent with the chromosome-wide shift seen in Fig 6B. By contrast, the plasmid origin-proximal region was sparsely represented, even though in a unidirectionally replicated circular plasmid this region is also where the replication fork completes a round of replication and is slightly enriched for AAG consensus PAMs. Thus, neither PAM density nor simple proximity to replication initiation or completion is sufficient to explain plasmid protospacer distribution. As with the chromosome, the *lexA3* Δ*recA* double mutant closely resembled the *recA* deletion alone, indicating that RecA exerts the dominant influence on plasmid spatial bias.

**Fig 7.**
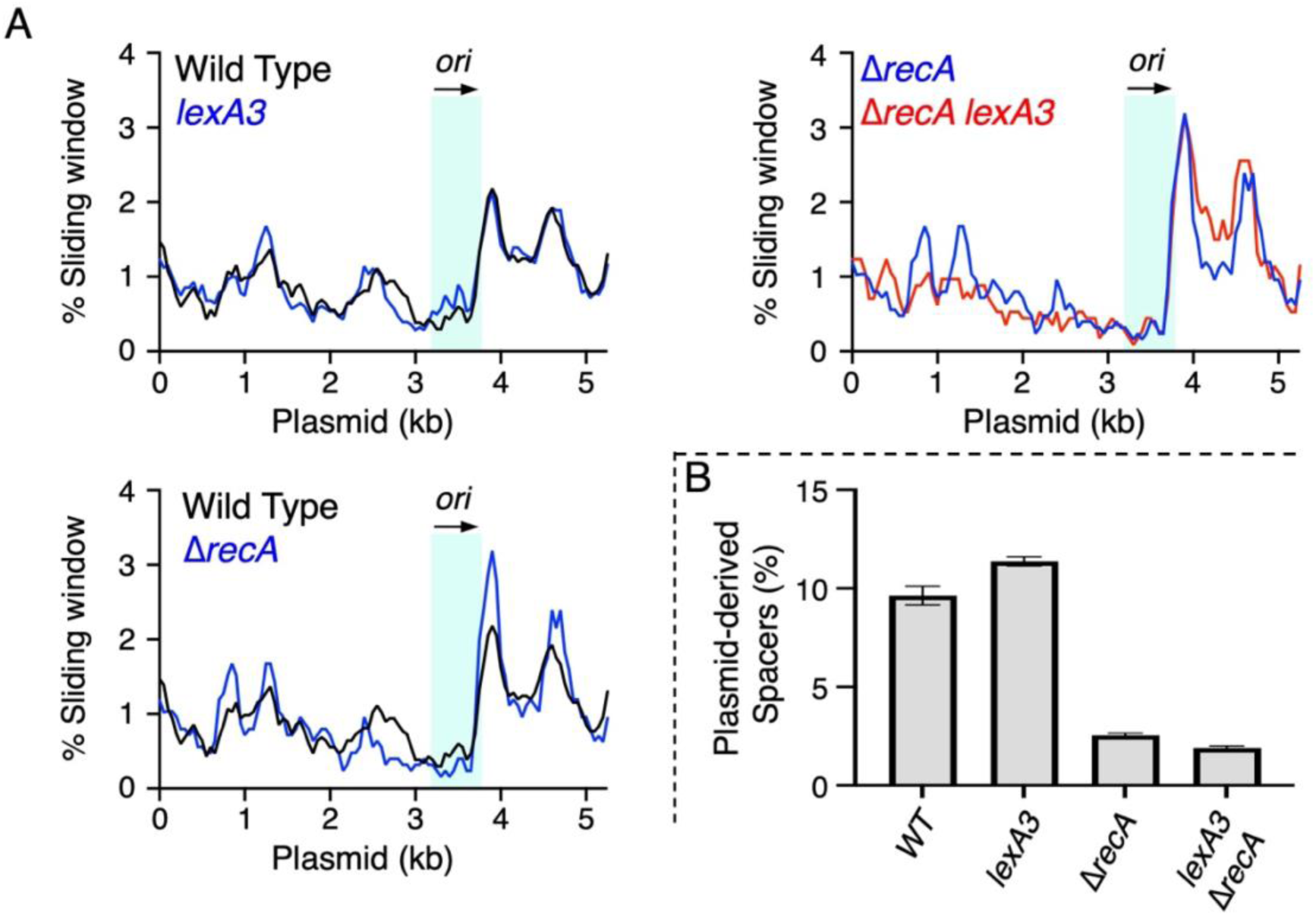
Plasmid protospacer distribution maps for *lexA3* and Δ*recA* mutants. Spacer sequences acquired in the experiment described in Fig 6 were further analyzed to assess plasmid-targeted acquisition. Spacer reads were mapped to the P*_araBAD_* expression plasmid (pRC1656). A. Protospacer density across the plasmid was calculated using a 250 bp sliding window with a step size of 50 bp. Strains were: wild-type CRISPR-Cas adaptation reporter (RC5311); *lexA3* (RC5353); Δ*recA* (RC5356) and *lexA3* Δ*recA* (RC5360). The linear plasmid maps start at the sequence 5-AAGAAACCAATTGTC, located at the start of the *araBAD* promoter. The tip of the arrow marks the exact replication initiation site, while the light blue box indicates the broader origin region. B. The percentage of total spacers that mapped to the plasmid is plotted for each strain, revealing differences in the relative recovery of plasmid-derived spacers. Error bars indicate 95% Wilson confidence intervals calculated from the underlying spacer counts. For plasmid-derived spacers, the denominator was total spacers.

These regulatory effects also altered the balance between chromosomal and plasmid-derived spacers. Compared with WT cells, Δ*recA* backgrounds showed increased sampling of chromosomal DNA relative to plasmid DNA, while *lexA3* modestly increased plasmid acquisition (Fig 7B). However, plasmid-versus-chromosome spacer usage does not directly predict successful adaptation in the papillation assay. As previously reported, growth on solid medium suppresses the strong plasmid bias characteristic of *E. coli* adaptation during liquid culture [11]. These observations therefore suggest that RecA and SOS alter not only the frequency of adaptation, but also the relative distribution of recovered spacers across chromosomal and plasmid DNA.

## DISCUSSION

CRISPR spacer acquisition in *E. coli* occurs within the highly structured microenvironment of a growing colony, where spatial gradients in nutrients, oxygen availability, growth rate, and stress generate pronounced physiological heterogeneity. Using the same colony-based reporter assay, we previously showed that this structure strongly influences both the frequency and genomic distribution of adaptation events, revealing pronounced heterogeneity that cannot be captured by population-average measurements [11].

Although successful adaptation increases with Cas1–Cas2 induction over a limited range, it saturates well below the level expected if spacer acquisition were a simple linear function of expression. This is consistent with a thresholded, lineage-restricted process with a ceiling set by cellular physiology rather than by promoter strength alone. This perspective is essential for interpreting assays that report on rare adaptive outcomes, such as papillation. Additional discussion of these interpretive considerations is provided in S1 Appendix.

### CRISPR–Cas Adaptation Interfaces with SOS and Cellular Physiology

Our study used the Type I-E system of *E. coli* to investigate how CRISPR–Cas adaptation interfaces with bacterial physiology, with a particular focus on the SOS response. This line of inquiry was prompted by a report that Cas1–Cas2 or Cas2 overexpression induces filamentation and elevates adaptation in *E. coli* BL21 [18]. We found that filamentation did not occur with Cas1, Cas2, or dCas1–Cas2 expressed individually, but wild-type Cas1–Cas2, expressed at a high level, triggered filamentation even in the absence of IHF (Fig 2A). This shows that the phenotype does not require integration at the CRISPR array and suggests that Cas1–Cas2 can perturb host physiology independently of spacer acquisition.

This physiological connection also extends to successful adaptation. Blocking LexA cleavage reduced adaptation, whereas nalidixic acid and mitomycin C enhanced adaptation only in cells capable of SOS induction, and constitutive SOS derepression did not mimic the stress-induced state. Together, these results support a model in which SOS-regulated cellular physiology, rather than DNA damage alone, contributes to successful CRISPR–Cas adaptation under genotoxic stress and during structured colony growth.

### LexA and RecA Shape the Genomic Landscape of Adaptation

The terminus region is a prominent protospacer hotspot in both *E. coli* BL21 and K-12 strains (Fig 6A; [11, 12]). As replication forks converge near the terminus, progression is impeded by Tus/ter sites, exposing lagging-strand DNA and generating substrates for repair. Supporting this, RecBCD, which functions in replication restart and double-strand break repair, has been implicated in protospacer generation in this region [12].

In *lexA3*, the enhanced enrichment of terminus-derived spacers likely reflects unresolved replication stress due to blocked induction of repair factors (Fig 6B). In contrast, the sharp loss of terminus spacers in Δ*recA* is consistent with a requirement for RecA-dependent repair, restart, or survival of lineages that acquire terminus-proximal spacers. Cells unable to restart replication at the terminus may be lost from the population, even as adaptation increases elsewhere. Together, these findings further highlight how CRISPR adaptation is intertwined with replication–repair pathways targeted by inhibitors such as PsiB, which may suppress both host repair and spacer acquisition (Fig 4).

These regulatory effects also altered the balance between chromosomal and plasmid-derived spacers. Compared with WT cells, Δ*recA* showed increased sampling of chromosomal DNA relative to plasmid DNA, whereas *lexA3* modestly increased plasmid acquisition (Fig 7B). These observations suggest that RecA and SOS influence not only the frequency of adaptation, but also the genomic context from which protospacers are acquired.

### RecA and RecBCD Have Interacting Effects on CRISPR–Cas Adaptation

The genetic data point to a more complex relationship between RecA, SOS induction, and DNA-end processing than a simple positive or negative role for RecA. Blocking LexA cleavage with *lexA3* reduced adaptation, whereas deleting *recA* increased adaptation (Figs 3A and 4A). This contrast shows that the Δ*recA* phenotype cannot be explained simply by loss of SOS induction. However, the RecA–RecBCD epistasis experiments (Fig 5) argue against interpreting the phenotype as definitive evidence for a discrete, standalone RecA inhibitory activity. Instead, complete loss of RecA appears to create a RecBCD-dependent DNA-processing state that favors successful adaptation.

The behaviour of RecX and PsiB creates a second genetic separation (Fig 4). Inhibitor-mediated perturbation of RecA in otherwise RecA-proficient cells reduced adaptation, and this effect was abolished in Δ*recA* cells, showing that the inhibitors act through RecA rather than through nonspecific toxicity or direct inhibition of Cas1–Cas2. The key distinction is therefore RecA absence versus RecA presence under inhibition. RecA-antagonized cells retain RecA protein and may therefore occupy a DNA-processing state distinct from that of Δ*recA* cells. This provides a framework for understanding why RecX and PsiB reduce adaptation in wild-type cells but have no further effect in Δ*recA*, while Δ*recA* itself increases adaptation through a RecBCD-dependent mechanism.

The use of PsiB also places this genetic separation in a natural mobile-element context. The F plasmid encodes PsiB, which inhibits RecA filament formation, while some phages encode proteins that inhibit SOS induction by stabilizing LexA–DNA complexes or blocking LexA autocleavage [27–29]. The recurrent targeting of RecA and SOS by mobile elements underscores that these repair pathways are important interfaces between hosts and invaders. Our results suggest that such antagonism may also influence CRISPR–Cas adaptation indirectly, by altering the physiological and repair states in which spacer acquisition becomes productive.

RecA and SOS-regulated functions might also act downstream of prespacer capture, for example by helping to resolve half-site intermediates at the CRISPR array. Since Cas1– Cas2 integration can generate single-ended or partially resolved structures, these could resemble DNA damage and require repair proteins for maturation into stable repeats. This idea is consistent with the observation that the *CRISPR-I* locus itself appears as a protospacer hotspot, an effect that is reduced in a *lexA3* background and eliminated in Δ*recA* and the *lexA3* Δ*recA* double mutant (Fig 6B). The differential behavior of the CRISPR-I hotspot in *lexA3* and Δr*ecA* further suggests that this local signal is not simply a passive consequence of SOS induction, but instead reflects a more specific dependence on RecA-linked physiological or repair functions.

The Δ*recA* phenotype is unlikely to reflect a direct role for RecA-mediated homologous recombination in spacer integration. Instead, increased adaptation in Δ*recA* cells more likely reflects indirect effects of RecA loss on DNA repair dynamics, replication-associated substrate availability, or the physiological states permissive for successful adaptation.

Genotoxic stress is expected to increase the availability of damaged DNA substrates accessible to Cas1–Cas2. However, adaptation in *lexA3* backgrounds was reduced still further following treatment with nalidixic acid or mitomycin C (Fig 3C), indicating that substrate availability alone is insufficient to explain the papillation phenotypes. Together with the RecX/PsiB results, this argues against a simple linear model in which DNA damage generates substrate and substrate abundance alone determines adaptation.

The RecA–RecBCD epistasis experiments (Fig 5) further refine the interpretation of the Δ*recA* phenotype. Loss of RecB or RecC suppressed the additional increase associated with deletion of *recA*, indicating that the full Δ*recA* hyperadaptation phenotype depends on RecBCD function. Thus, the Δ*recA* phenotype is unlikely to reflect simple removal of an inhibitory RecA activity. Instead, it appears to arise from a RecBCD-dependent repair/substrate-processing state that becomes prominent when RecA is absent. This is consistent with the possibility that, when homologous recombination is blocked, RecBCD-dependent processing of unrepaired DNA generates or exposes additional substrates for adaptation. However, this genetic interaction does not identify the relevant DNA intermediate, and therefore does not show that extensive RecBCD-dependent degradation of chromosomal DNA is responsible. More generally, DNA degradation is not equivalent to productive prespacer generation: unrepaired DNA ends may be degraded to non-productive products, processed by alternative nucleases, or lost through death of the damaged cell. Loss of RecA could instead prolong the availability of RecBCD-unwound DNA, prevent RecA from coating or channeling DNA into recombination, alter repair-pathway choice, or modify subsequent processing by other nucleases.

These interactions also need to be interpreted in the context of SOS feedback. RecA is required for LexA cleavage and SOS induction, and *recA* itself is SOS-regulated, creating positive feedback between DNA damage sensing and RecA availability. RecBCD contributes to SOS signal generation by processing double-strand ends and loading RecA onto ssDNA, whereas *recB* and *recC* are not themselves canonical SOS genes. Thus, *recB* or *recC* mutants may decouple the accumulation of unrepaired DNA lesions from RecBCD-dependent RecA loading, SOS signalling, and substrate processing. In this context, the phenotypes of *lexA3*, Δ*recA*, Δ*recA lexA3*, Δ*recB*, Δ*recC*, and the Δ*recA ΔrecB* and Δ*recA ΔrecC* double mutants are best viewed as different perturbations of a coupled DNA-processing and stress-response network, rather than as independent tests of substrate abundance.

The behaviour of the Δ*recB* and Δ*recC* single mutants also argues against a model in which RecBCD nuclease products are obligatory precursors of adaptation. Both mutants retained substantial papillation and, under the lower-expression P*_araMari_* conditions used here, showed increased papillation relative to wild type. This differs from a previous report which reported reduced acquisition in *recB*, *recC*, and *recD* mutants in a liquid-culture assay using T7-driven Cas1–Cas2 overexpression [12]. The different outcomes may reflect differences in Cas1–Cas2 expression, growth format, and assay readout. One possibility is that RecBCD-deficient cells are especially sensitive to Cas1–Cas2-associated DNA damage or stress under high-expression conditions, reducing the recovery or expansion of adapted lineages. By contrast, lower P*_araMari_* expression may allow RecBCD-deficient cells to remain viable enough for adaptation events to be detected during prolonged colony growth. Thus, liquid-culture array expansion and colony papillation may emphasize different balances between spacer acquisition, DNA damage, repair capacity, and lineage survival.

Taken together, these results argue for a network model rather than a single RecA inhibitory function. RecA promotes adaptation through SOS induction and through RecA-dependent functions that support productive repair or maturation of adaptation intermediates. At the same time, loss of RecA redirects DNA-end processing in a RecBCD-dependent manner that can increase successful adaptation under structured-colony conditions. RecB and RecC are required for the full increase produced by loss of RecA, but RecBCD function is not essential for substantial adaptation under the conditions used here. Thus, the Δ*recA* phenotype is best interpreted as arising from altered DNA repair and substrate-processing physiology, rather than as evidence for a discrete biochemical inhibitory activity of RecA.

More broadly, successful adaptation appears to reflect the balance among RecBCD-mediated DNA processing, RecA-dependent repair or DNA occupancy, SOS-regulated physiology, alternative processing pathways, and the survival and expansion of adapted lineages. This framework explains why SOS blockade, RecA deletion, RecA antagonism, and RecBCD loss produce distinct but interacting phenotypes. It also places CRISPR–Cas adaptation within the broader DNA-damage and repair network of the cell, where the availability of potential prespacers is only one component of the pathway leading to heritable spacer acquisition.

## Material & Methods

### Reagents and Enzymes

Chemicals were obtained from Sigma, BDH Laboratory Supplies, Fisher Scientific, and Thermo Scientific. Enzymes were purchased from New England Biolabs and used according to the manufacturer’s instructions except where stated.

### Plasmids and Molecular Biology

Complete nucleotide sequences of all plasmids used are provided in S1 Table. Plasmids were constructed by Gibson assembly [30], and junction sequences were verified by Sanger sequencing. Genes under the P*_araBAD_* promoter in pBAD [31] were cloned in frame with the first start codon downstream of the ribosome binding site. The P*_araMari_* and P*_vanMari_* promoters were obtained from plasmids pAJM.677 and pAJM.773, respectively [25], and our genes of interest were cloned to replace the YFP coding sequence.

The following plasmids were generated during this work: Cas1–Cas2 (pRC1656, pRC1657, pRC2747, pRC3058), Cas1 (pRC1653), Cas2 (pRC1655), dCas1–Cas2 (pRC1658), SulA (pRC2781), RecX (pRC3017), PsiB (pRC3028). The SOS reporter plasmid, in which GFP is under control of P*_sulA_*, was pRC2790 (pZA31-sulA-GFP, Addgene plasmid number #78493). Plasmid origins and promoters are indicated in figure legends. PCR reactions were performed using Q5 polymerase (New England Biolabs). Products were visualized on 1.0% TBE-buffered agarose gels run at 2.7 V/cm, stained with ethidium bromide (0.3 µg/mL), and imaged under UV. If needed, DNA was gel purified using Qiagen QIAquick columns. CRISPR array expansion was detected by PCR across the leader–repeat junction of the lacZ reporter in strain RC5311 using primers: 5′-ggctagcaggaggaattcacc and 5′-cgcatcgtaaccgtgcatctg.

### Bacterial Growth Conditions and Papillation Assays

LB-Lennox consisted of 10 g/L tryptone, 5 g/L yeast extract, and 5 g/L NaCl. Bacteria were grown at 37 °C in LB-Lennox broth or on LB-Lennox agar (1.5%). For papillation assays, plates were supplemented with 0.1% L-lactose and 40 µg/mL X-gal. Each plate was seeded with 25–50 colonies. Where appropriate, antibiotics were added as follows: ampicillin (100 µg/mL), kanamycin (50 µg/mL), chloramphenicol (30 µg/mL). Inducers and sub-inhibitory concentrations of antibiotics were used as described in figure legends. Papillating colonies were imaged using a Cole Parmer National Optical Professional Stereozoom Microscope with Motic Images Plus 3.1 software. Papillae were counted from high-resolution colony images, with a multimodal AI tool used to assist initial scoring. All counts used for statistical analysis were manually checked against the source images. All counts used for analysis were checked manually by the authors against the original images. Overnight cultures were preserved in 50% glycerol at –80 °C. Chromosomal markers were maintained with tetracycline (12.5 µg/mL), kanamycin (30 µg/mL) or chloramphenicol (5 µg/mL) but selection was relaxed during experiments.

### Genome Engineering and Bacterial Strains

*E. coli* BL21-AI (*λ-, F^−^, ompT, hsdS_B_ (r ^−^, m_B_ ^−^), gal, dcm, araB::T7RNAP-tetA*) (ThermoFisher) was used in Fig 2. *E. coli* ER1793 (*λ-, F-, fhuA2,* Δ*[lacZ]r1, glnV44, e14-[McrA-], trp-31, his-1, rpsL104, xyl-7, mtl-2, metB1,* Δ*(mcrC-mrr)114::IS10*) supports a high level of lambda red recombineering and was used as an intermediate host as described below. The papillation reporter strain was RC5311 (JB028) (*F-, λ^-^, rph-1,* Δ*lacZ,* Δ*CRISPR-II, araBAD::T7 RNAP tetA, argE::[J23100 CRISPR-II (T) lacZ– reporter]*) [11]. *E. coli* MLS989 is a derivative of MG1655 with a Type I-E *CRISPR-II* (T) YFP adaptation-reporter inserted at the *galK* locus (*λ-F-rph-1 araB::T7pol–tetA* Δ*araA* Δ*cas3–CRISPR-I* Δ*CRISPR-II galK::[CRISPR-II (T) yfp–reporter]*) [26]. MLS990 is MLS989 with a single base added to the reporter to restore the yfp open reading frame to act as a positive control [26]. All other strains were *E. coli* K12 MG1655 (*λ-, F-, rph-1*) or its derivatives as indicated below.

Genome modifications were performed using P1 transduction and lambda Red recombination [32, 33]. Antibiotic markers were removed by FLP recombination between flanking *frt* sites where possible.

The *lexA3* allele was transduced into the reporter strains RC5311 and MLS989, from DE407 [34] using a nearby *malB::Tn9* insertion as a selectable marker to yield RC5353 (*F-, λ^-^, rph-1,* Δ*lacZ,* Δ*CRISPR-II, lexA3, araBAD::T7 RNAP tetA, malB::Tn9 argE::[J23100 CRISPR-II (T) lacZ-Reporter]*) and RC5373 (*λ-F-rph-1 araB::T7pol–tetA*Δ*araA* Δ*cas3–CRISPR-I* Δ*CRISPR-II galK::[CRISPR-II (T) yfp–reporter] lexA3 malB::Tn9*). This marker was then removed by P1 transduction of a nearby gene knockout (*yjaB::kan*) from the Keio Collection [35]. The kanamycin marker was deleted by FLP recombination between flanking frt sites to yield RC5355 (*F-, λ^-^, rph-1,* Δ*lacZ,* Δ*CRISPR-II,* Δ*yjaB, lexA3, araBAD::T7 RNAP tetA, argE::[J23100 CRISPR-II (T) lacZ– reporter]*). To make the *lexA51* strain, Δ*sulA* was introduced into the reporter strain by P1 transduction from the Keio collection. The kanamycin marker was deleted by FLP recombination between flanking *frt* sites. The *lexA51* allele was then transduced from DE406 [36] using a nearby *malB::Tn9* insertion as a selectable marker to yield RC5370 (*F-, λ^-^, rph-1,* Δ*lacZ,* Δ*CRISPR-II,* Δ*sulA, lexA51, araBAD::T7 RNAP tetA, malB::Tn9 argE::[J23100 CRISPR-II (T) lacZ-Reporter]*). Further P1 transductions were used to introduce Δ*recA* from the Keio Collection into RC5311, RC5355 and MLS989 and to yield RC5356 (*F-, λ^-^, rph-1,* Δ*lacZ,* Δ*CRISPR-II, araBAD::T7 RNAP tetA, argE::[J23100 CRISPR-II (T) lacZ–reporter]* Δ*recA*), RC5360 (*F-, λ^-^, rph-1,* Δ*lacZ,* Δ*CRISPR-II,* Δ*yjaB, lexA3, araBAD::T7 RNAP tetA, argE::[J23100 CRISPR-II (T) lacZ–reporter] lexA3* Δ*recA*), and RC5288 (*λ-F-rph-1 araB::T7pol–tetA* Δ*araA* Δ*cas3–CRISPR-I* Δ*CRISPR-II galK::[CRISPR-II (T) yfp–reporter]* Δ*recA*). The *recA* allele was restored by lambda Red recombination in RC5356 to yield RC5357 (*F-, λ^-^, rph-1,* Δ*lacZ,* Δ*CRISPR-II, araBAD::T7 RNAP tetA, argE::[J23100 CRISPR-II (T) lacZ–reporter]* Δ*recA::recA⁺*). Further P1 transductions were used to introduce Δ*recB and* Δ*recC* from the Keio Collection into RC5311 and RC5356, to yield RC5368 (*F-, λ^-^, rph-1,* Δ*lacZ,* Δ*CRISPR-II, araBAD::T7 RNAP tetA, argE::[J23100 CRISPR-II (T) lacZ–reporter]* Δ*recB*), RC5369 (*F-, λ^-^, rph-1,* Δ*lacZ,* Δ*CRISPR-II, araBAD::T7 RNAP tetA, argE::[J23100 CRISPR-II (T) lacZ–reporter]* Δ*recC*), RC5371 (*F-, λ^-^, rph-1,* Δ*lacZ,* Δ*CRISPR-II, araBAD::T7 RNAP tetA, argE::[J23100 CRISPR-II (T) lacZ–reporter]* Δ*recA* Δ*recB*) and RC5372 (*F-, λ^-^, rph-1,* Δ*lacZ,* Δ*CRISPR-II, araBAD::T7 RNAP tetA, argE::[J23100 CRISPR-II (T) lacZ– reporter]* Δ*recA* Δ*recC*). The *lexA3* and Δ*ihfα* alleles were also transduced into BL21-AI, yielding RC5250 (*λ-, F^−^, ompT, hsdS_B_ (r ^−^, m_B_ ^−^), gal, dcm, araB::T7RNAP-tetA lexA3, malB::Tn9*) and RC5362 (*λ-, F^−^, ompT, hsdS_B_ (r ^−^, m_B_ ^−^), gal, dcm, araB::T7RNAP-tetA* Δ*ihfα*).

### Microscopy, Fluorescence Imaging and Cell Sorting

To assess filamentation, overnight cultures were washed in M9 salts and dried onto a 1.2% agarose pad, placed on a 35 mm glass bottom culture dish, and viewed under a Zeiss Axio Observer 7 microscope with Ph 3 annulus and 63×, 1.25 N.A objective using a T-PMT.2 detector. Representative fields are presented.

For quantification of the GFP–SOS reporter in liquid media (Fig 2B), overnight cultures were adjusted to OD_600_ = 2, serially diluted, and 175 µL was loaded per well in a 96-well plate. GFP fluorescence was measured using the Cy2 setting on an Amersham Typhoon Imager. For quantification of the GFP–SOS reporter on solid media (Fig 3), ∼50 colonies were grown overnight on plates. Cells were pooled, resuspended in LB, normalized to OD600 = 2, and fluorescence was quantified as above. Cells were sorted using a Beckman Coulter CytoFLEX S flow cytometer in the University of Nottingham Flow Cytometry Facility. Gating conditions were established to distinguish between YFP^-^ and YFP^+^ cells, using *E. coli* strains MLS989 and MLS990 as negative and positive controls, respectively. Samples were run at 60 μL/min until 20,000 cells had been sorted. Data were analyzed using the CytExpert software package.

For analysis of spacer acquisition by flow cytometry after growth on solid media, MLS989 was transformed with pRC2747 (P_araMari_–Cas1–Cas2). Transformed cells were plated on LB agar supplemented with kanamycin and 0.02% L-arabinose at a density of approximately 50 colonies per plate and incubated at 37 °C for 5 days. Colonies were then pooled, the OD₆₀₀ was measured, and cells were diluted to approximately 10⁶ per mL for flow cytometry analysis as described above.

### Illumina Sequencing of Acquired Spacers

Papillation assays were performed using Cas1–Cas2 under P*_araBAD_* control (pRC1656), induced with 0.002% arabinose. The strains used were: wild-type CRISPR–Cas adaptation reporter (RC5311), *lexA3* (RC5355), Δ*recA* (RC5356), and *lexA3* Δ*recA* (RC5360). Colonies from 25 (or 50 for *lexA3*) plates were pooled in 50 mL ice-cold LB. 500 µL (or 1 mL for *lexA3*) was washed in PBS and inoculated into 200 mL M9 minimal lactose medium (6 g/L Na2HPO4, 3 g/L KH2PO4, 0.5 g/L NaCl, 1 g/L NH4Cl, 1 mM MgSO4, 0.1 mM CaCl_2_, 5 µg/mL thiamine, 0.1% lactose). Cultures were grown for 48 h at 37 °C with aeration. DNA was amplified by PCR (150 µL Q5 High GC reaction, primers: 5′-GGTCTTAATGAATGGCCGGG and 5′-CATGGATCCGAAGTCGAGC), gel purified, and sequenced by Illumina (Azenta/Genewiz).

Spacer sequences were extracted from raw sequence reads and mapped to the genome and plasmid using Python 3.13.2 with the packages Biopython 1.85, Matplotlib 3.10.1, NumPy 2.2.4, and Pandas 2.2.3. Custom Python scripts are provided in S2 Table. Chromosomal spacer distributions were plotted in 10 kb bins or using 100 kb sliding windows. Plasmid distributions were plotted using 250 bp sliding windows. Unique spacers were plotted for all conditions, except for plasmid derived spacers percentage where duplicates were retained.

## Data Availability

Data are available under BioProject PRJNA1327804. All code to replicate the analysis can be found in S2 Table.

## Acknowledgements

We would like to thank Roger Woodgate for insightful advice and providing *recA* and *lexA* mutants.

## Funding

This work was funded by The Leverhulme Trust grant RPG-2020-079 to RC.

## Author Contributions

HE conceived the study. HE and RC designed the experiments. HE conducted the experiments. JB built the reporter system. CC wrote and executed the Python scripts. RC and HE wrote the manuscript. All authors read and approved the manuscript.

---

**S1 Fig. A quantitative papillation assay for CRISPR–Cas adaptation**

A. A promoter and ribosome binding site drive transcription and translation through the reporter CRISPR array toward *lacZ*. However, *lacZ* is not expressed due to an in-frame stop codon. Acquisition of a 61 bp spacer-repeat unit restores the reading frame, enabling *lacZ* expression. Cells are transformed with a Cas1–Cas2 expression vector and plated on LB agar supplemented with 0.1% lactose and X-gal. Cells that acquire a spacer-repeat unit during colony development form blue-stained papillae. Papillation assay for CRISPR Type I-E adaptation in *E. coli*, adapted from [11].

B. To demonstrate the reproducibility of the papillation assay, representative whole-plate photographs are shown with Cas1–Cas2 induced using 0.002% or 0.02% arabinose, as indicated. Plates were seeded at a density of approximately 25–50 colonies per plate and photographed after incubation at 37 °C for five days. The papillation patterns were reproducible across replicate plates. High-resolution images of six additional examples of each plate type are provided in S1 File.

C. Representative colony arrays from an arabinose titration of Cas1–Cas2 expression from P*_araBAD_*. Ten colonies were photographed at each indicated concentration and are shown in rows from left to right. Discrete, convincingly blue papillae were counted; diffuse pale-blue central staining and light-reflecting structures were excluded, and confluent regions were not subdivided into an arbitrary number of events. The graph shows the mean number of papillae per colony plotted against arabinose concentration on a logarithmic scale. Points show mean ± SD (n=10 colonies). The line is an ordinary least-squares regression fitted across all countable induced concentrations from 0.0005 to 0.002% arabinose (R^2^=0.976). Colonies at 0.005% arabinose were confluent and could not be counted. See S1 Appendix for further discussion and a detailed description of the counting method.

D. As in C, but using the P*_araMari_* Cas1–Cas2 expression system. The regression was fitted across all countable induced concentrations from 0.001 to 0.05% arabinose (R^2^=0.932). The approximately log-linear relationship is consistent with the expected transfer function of the evolved P_araMari_ sensor.

**S2 Fig. Modest SOS induction at standard Cas1–Cas2 expression levels.**

Cas1–Cas2 expression was induced in *E. coli* RC5311 (JB028) from pRC1656 (*P_araBAD_*) using 0.002 and 0.2% arabinose. SOS was measured by GFP expression from the *sulA* promoter on the GFP SOS reporter plasmid (pRC2790). GFP fluorescence was measured in triplicate as described in the Materials and Methods section; bars show the mean ± SEM. Conditions were compared using a two-sample *t*-test. P values: ***P < 0.001; **P < 0.01.

**S3 Fig. Constitutive SOS derepression does not enhance CRISPR–Cas adaptation** Papillation assays were performed using the wild-type CRISPR–Cas adaptation reporter strain (RC5311) and a *lexA51 sulA* double mutant (RC5370) with constitutive SOS derepression. Cas1–Cas2 was expressed from pRC2747 under P*_araMari_* control. Despite constitutive derepression of the SOS regulon, papillation was not enhanced relative to wild type and was modestly reduced.

**S4 Fig. CRISPR–Cas adaptation is increased in Δ*recA* cells**

The YFP adaptation reporter strain (MLS989) and its Δ*recA* derivative (RC5288) were transformed with the P*_araMari_* Cas1–Cas2 expression plasmid pRC2747. Cas1–Cas2 expression was induced with 0.02% arabinose. After five days of incubation on mock papillation plates lacking lactose and X-gal, cells were pooled and analyzed by fluorescence-activated cell sorting. Fluorescence measurements were performed in triplicate; bars show the mean ± SEM. Conditions were compared using a two-sample t-test. P value: *, <0.05.

**S5 Fig. Representative colonies used for quantification of RecA–RecBCD epistasis.**

Sets of 10 colonies per genotype are shown for the papillation assay quantified in Fig 5. Papillae in Δ*recB* and Δ*recC* backgrounds were numerous and sometimes diffuse, so small differences among RecBCD-deficient genotypes should be interpreted cautiously.

**S1 File. High-resolution images of papillation plates to demonstrate reproducibility.**

**S1 Table. Plasmid sequences S2 Table. Python scripts**

**S3 Table. Distribution of cell-length phenotypes in representative micrographs** Representative .czi micrographs were scored using a custom Python-based image-analysis workflow. Cell lengths were calibrated from embedded image metadata and the 50 µm scale bar, and segmented objects were assigned to predefined length-based categories: normal rods (<4 µm), mildly elongated (4 to <8 µm), filaments (8 to <15 µm), or extreme filaments (≥15 µm). Septating, overlapping, clustered, or poorly resolved cells were assigned to an ambiguous category. Values represent approximate percentages of scored objects within the fields shown.

## Notes

### Competing Interest Statement

The authors have declared no competing interest.

