## Supplementary Data for "RecA status determines SOS- and RecBCD-dependent outcomes in CRISPR–Cas adaptation"

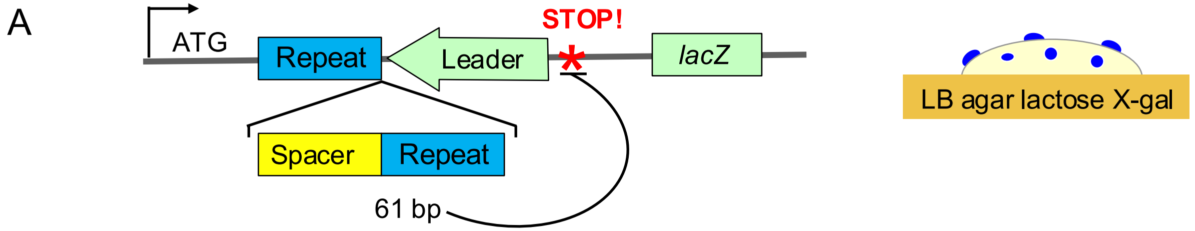


B

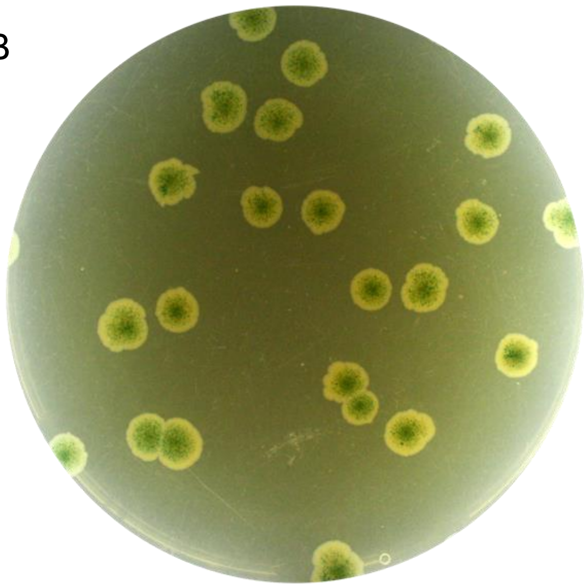

0.002% arabinose $P_{araBAD}$

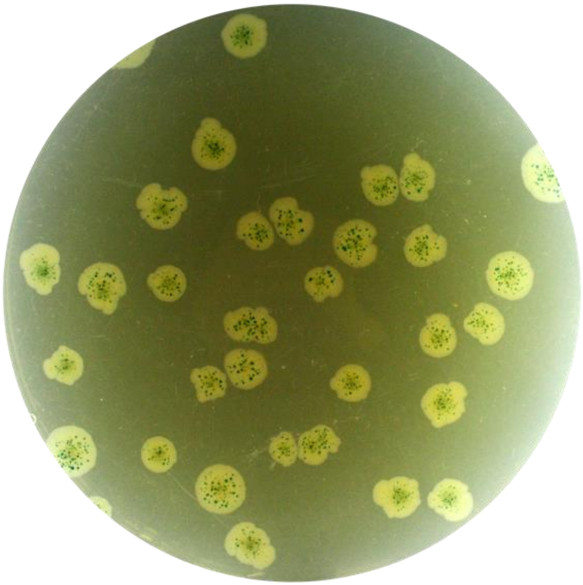

0.02% arabinose $P_{araMari}$

S1 Fig.

C

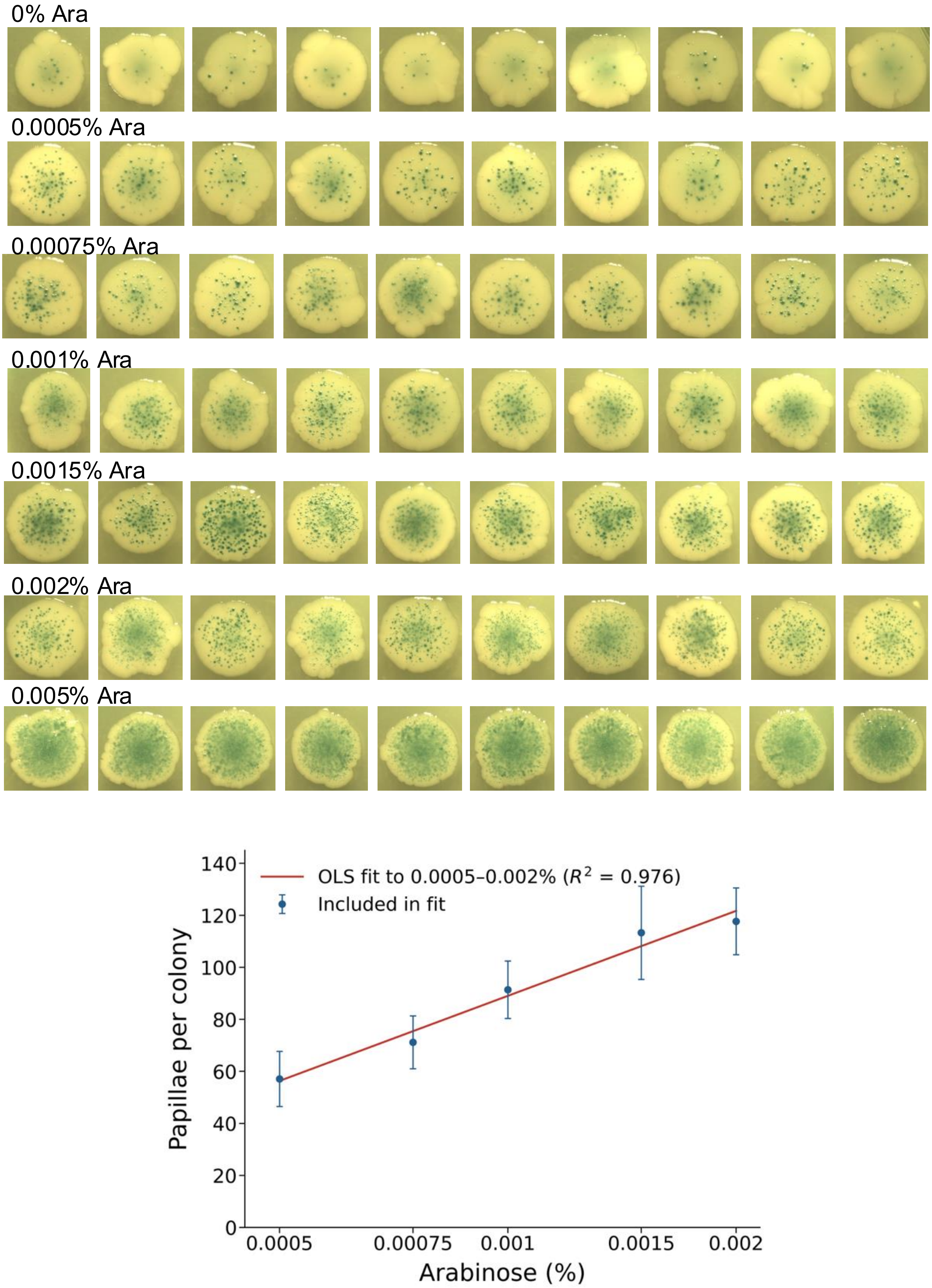

0% Ara
0.0005% Ara
0.00075% Ara
0.001% Ara
0.0015% Ara
0.002% Ara
0.005% Ara
OLS fit to 0.0005–0.002% ($R^2$ = 0.976)
Included in fit
Papillae per colony
140
120
100
80
60
40
20
0
0.0005
0.00075
0.001
0.0015
0.002
Arabinose (%)


S1 Fig.

D

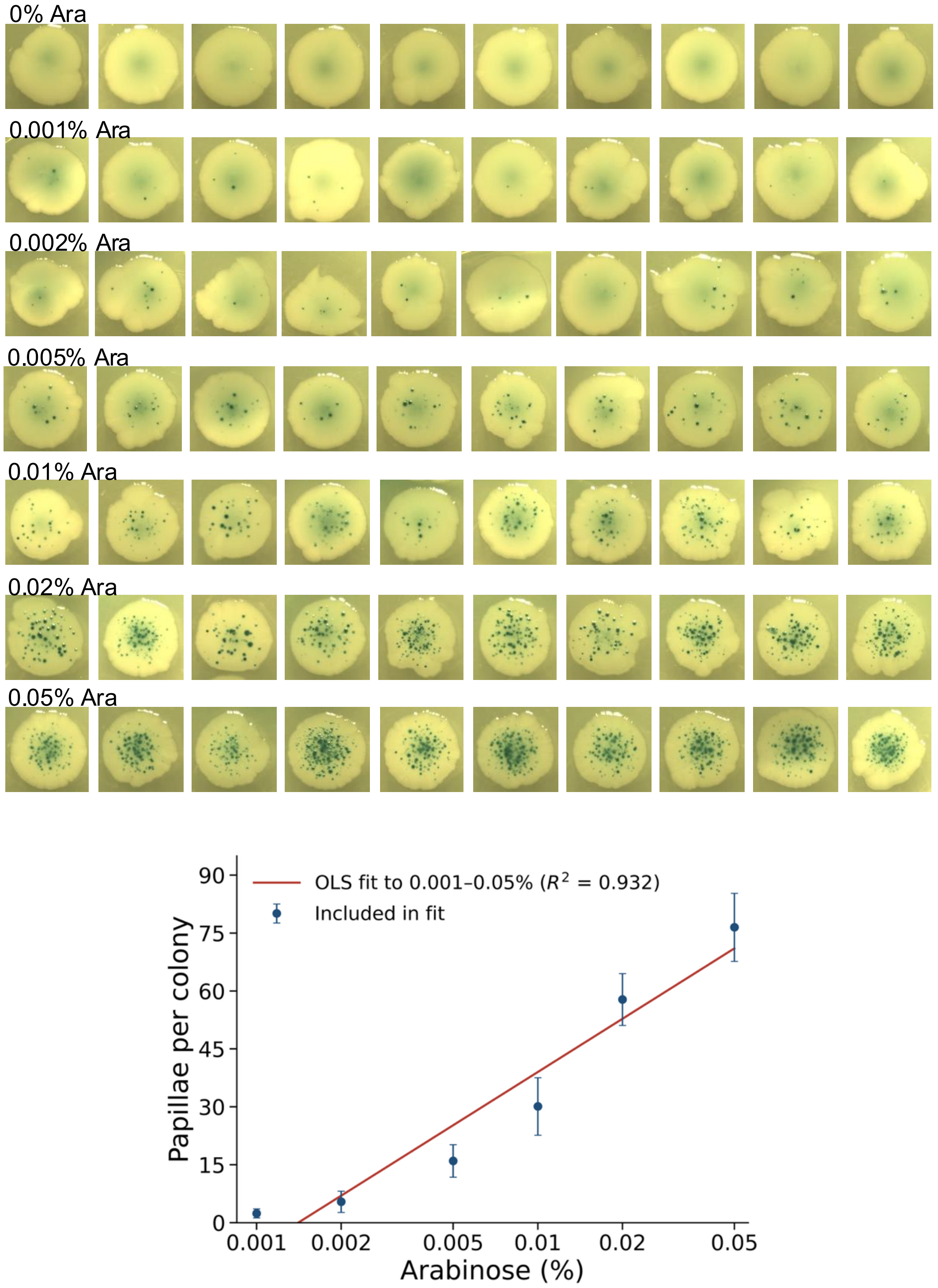

0% Ara
0.001% Ara
0.002% Ara
0.005% Ara
0.01% Ara
0.02% Ara
0.05% Ara
OLS fit to 0.001–0.05% ($R^2$ = 0.932)
Included in fit
Papillae per colony
Arabinose (%)
0
15
30
45
60
75
90
0.001
0.002
0.005
0.01
0.02
0.05


S1 Fig.

**S1 Fig. A quantitative papillation assay for CRISPR–Cas adaptation**

**A**. A promoter and ribosome binding site drive transcription and translation through the reporter CRISPR array toward *lacZ*. However, *lacZ* is not expressed due to an in-frame stop codon. Acquisition of a 61 bp spacer-repeat unit restores the reading frame, enabling *lacZ* expression. Cells are transformed with a Cas1–Cas2 expression vector and plated on LB agar supplemented with 0.1% lactose and X-gal. Cells that acquire a spacer-repeat unit during colony development form blue-stained papillae. Papillation assay for CRISPR Type I-E adaptation in *E. coli*, adapted from doi.org/10.1093/nar/gkaf1044.

**B.** To demonstrate the reproducibility of the papillation assay, representative whole-plate photographs are shown with Cas1–Cas2 induced using 0.002% or 0.02% arabinose, as indicated. Plates were seeded at a density of approximately 25–50 colonies per plate and photographed after incubation at 37 °C for five days. The papillation patterns were reproducible across replicate plates. High-resolution images of six additional examples of each plate type are provided in S1 File.

**C.** Representative colony arrays from an arabinose titration of Cas1–Cas2 expression from ParaBAD. Ten colonies were photographed at each indicated concentration and are shown in rows from left to right. Discrete, convincingly blue papillae were counted; diffuse pale-blue central staining and light-reflecting structures were excluded, and confluent regions were not subdivided into an arbitrary number of events. The graph shows the mean number of papillae per colony plotted against arabinose concentration on a logarithmic scale. Points show mean ± SD (n=10 colonies). The line is an ordinary least-squares regression fitted across all countable induced concentrations from 0.0005 to 0.002% arabinose ($R^2$=0.976). Colonies at 0.005% arabinose were confluent and could not be counted. See S1 Appendix for further discussion and a detailed description of the counting method.

**D.** As in C, but using the ParaMari Cas1–Cas2 expression system. The regression was fitted across all countable induced concentrations from 0.001 to 0.05% arabinose ($R^2$=0.932). The approximately log-linear relationship is consistent with the expected transfer function of the evolved $P_{araMari}$ sensor.

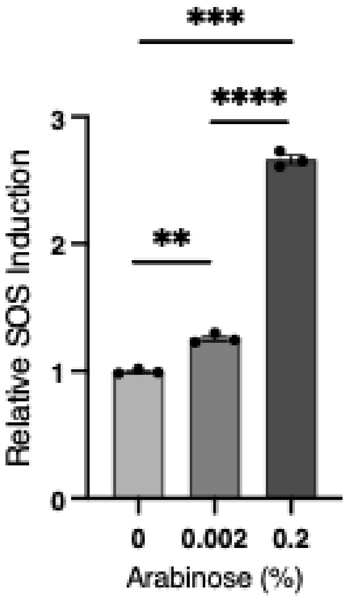


| Ara (%) | 1 | 2 | 3 | Average |
|---|---|---|---|---|
| 0 | 0.98 | 0.99 | 1.02 | 1.00 |
| 0.002 | 1.23 | 1.25 | 1.30 | 1.26 |
| 0.2 | 2.61 | 2.73 | 2.65 | 2.66 |

**S2 Fig. Modest SOS induction at standard Cas1–Cas2 expression levels.** Cas1–Cas2 expression was induced in *E. coli* RC5311 (JB028) from pRC1656 ($P_{araBAD}$) using 0.002 and 0.2% arabinose. SOS was measured by GFP expression from the *sulA* promoter on the GFP SOS reporter plasmid (pRC2790). GFP fluorescence was measured in triplicate as described in the Materials and Methods section; bars show the mean ± SEM. Conditions were compared using a two-sample *t*-test. P values: ***P < 0.001; **P < 0.01.

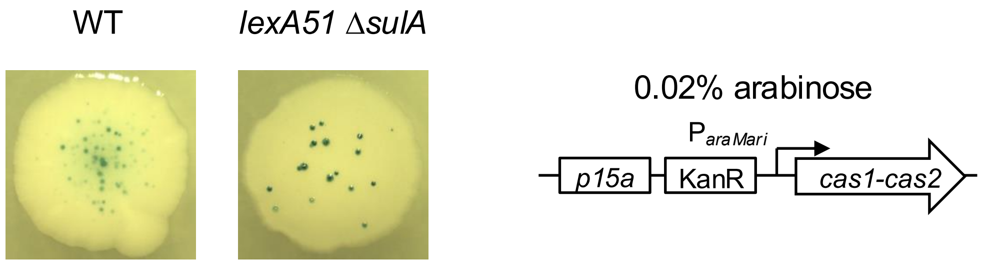


**S3 Fig. Constitutive SOS derepression does not enhance CRISPR–Cas adaptation**

Papillation assays were performed using the wild-type CRISPR–Cas adaptation reporter strain (RC5311) and a *lexA51 sulA* double mutant (RC5370) with constitutive SOS derepression. Cas1–Cas2 was expressed from pRC2747 under $P_{araMari}$ control. Despite constitutive derepression of the SOS regulon, papillation was not enhanced relative to wild type and was modestly reduced.

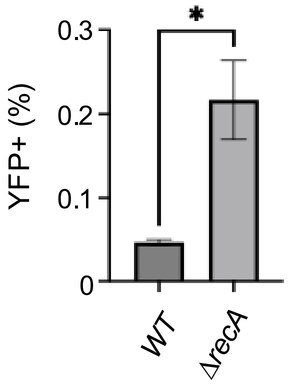


**S4 Fig. CRISPR–Cas adaptation is increased in Δ*recA* cells**

The YFP adaptation reporter strain (MLS989) and its Δ*recA* derivative (RC5288) were transformed with the $P_{araMari}$ Cas1–Cas2 expression plasmid pRC2747. Cas1–Cas2 expression was induced with 0.02% arabinose. After five days of incubation on mock papillation plates lacking lactose and X-gal, cells were pooled and analyzed by fluorescence-activated cell sorting. Fluorescence measurements were performed in triplicate; bars show the mean ± SEM. Conditions were compared using a two-sample t-test. P value: *, <0.05.

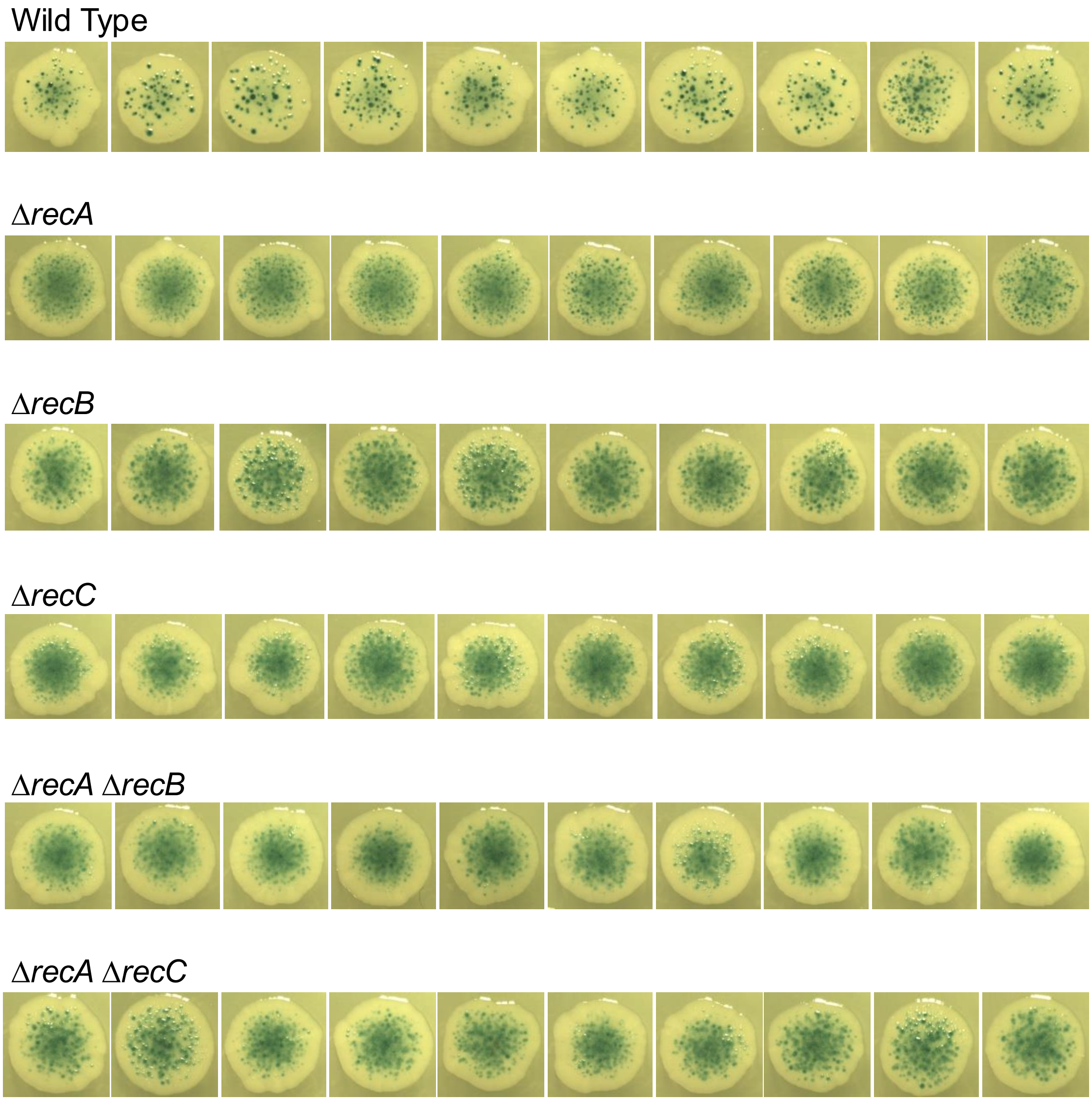


**S5 Fig. Representative colonies used for quantification of RecA–RecBCD epistasis.**

Sets of 10 colonies per genotype are shown for the papillation assay quantified in Fig 6. Papillae in Δ*recB* and Δ*recC* backgrounds were numerous and sometimes diffuse, so small differences among RecBCD-deficient genotypes should be interpreted cautiously.

### S1 Appendix. Quantification and interpretation of rare adaptation events in structured colony assays

#### Quantification and dynamic range of the papillation assay

To determine how papillation responds to Cas1–Cas2 expression, arabinose was titrated across the expression ranges of the stronger $P_{araBAD}$ system and the low-expression $P_{araMari}$ system. For each arabinose concentration, ten representative colonies were photographed from the corresponding plate. The resulting arrays therefore contain 70 colonies for each promoter (S1C and S1D Figs.). Papillae were counted by machine-assisted image analysis using rules established by comparison with manual inspection. Only discrete, convincingly blue papillae were counted. Clear or light-reflecting structures associated with colony deformation were excluded, diffuse pale-blue staining was not counted as papillae, and merged regions were treated as unresolved rather than subdivided into an arbitrary number of events. A random sample of countable colonies spanning the experimental range was also counted manually by the experimenter; these counts agreed with the machine-assisted values to within two papillae for colonies within the countable, non-confluent range of the assay (see below). Means and standard deviations were calculated from the ten colonies at each concentration.

The $P_{araBAD}$ titration first defined the practical counting range of the assay. Papillation increased from a mean of 10.6 without arabinose to 117.7 at 0.002% arabinose. Across the nonzero, countable concentrations from 0.0005 to 0.002%, the response was well described by a straight line on the semilogarithmic plot ($R^2$=0.976; S1C Fig). This interval spans fourfold in inducer concentration, and the plot reports the integrated response of the complete papillation assay rather than a direct measurement of promoter output. Nevertheless, it provides a strong empirical description of the assay response across the countable induced range.

At 0.002% arabinose, the $P_{araBAD}$ colonies were visibly crowded and some central papillae were difficult to resolve. However, the observed mean remained close to the semilogarithmic fit, and the numerical data therefore do not by themselves demonstrate counting saturation at this concentration. At 0.005%, by contrast, individual papillae were extensively confluent and could not be counted. Nevertheless, visual inspection showed substantially greater papillation than at 0.002%, indicating that the biological response had continued to increase after quantitative enumeration became impossible. Thus, 0.002% approaches the upper practical counting range. Beyond this limit, the assay retains a qualitative indication of further increases in papillation but does not provide a reliable numerical estimate.

The $P_{araMari}$ titration extended this analysis across a substantially wider inducer range. The mean number of papillae increased from zero without arabinose to 57.8 at 0.02% arabinose and 76.5 at 0.05% arabinose. Across all countable induced concentrations from 0.001 to 0.05%, the data were well described by a straight line on the semilogarithmic plot ($R^2$=0.932; S1D Fig.). This log-linear response is consistent with the central portion of the expected sigmoidal transfer function of the $P_{araMari}$ sensor, which was extensively evolved to reduce the non-ideal behavior of the native $P_{araBAD}$ system [1]. Notably, the relationship extends across a 50-fold range of nonzero inducer concentrations, compared with the fourfold range sampled by the countable induced $P_{araBAD}$ series.

Visual inspection of the colonies at 0.05% arabinose showed that individual papillae remained readily resolvable, consistent with the observed mean of 76.5. The standard 0.02% $P_{araMari}$ condition used in the paper therefore lies below the upper end of the tested countable response and leaves quantitative capacity to detect experimental variables that increase papillation, including the mutant effects.

As an internal check on the reproducibility of the counting procedure, $P_{araBAD}$ colonies at 0.0005% arabinose, with a mean of 57.1 papillae, closely resembled $P_{araMari}$ colonies at 0.02% arabinose, with a mean of 57.8. Comparable papillation morphologies therefore received almost identical scores in independent experiments performed on different days using reporter strains carrying different Cas1–Cas2 expression plasmids. It is not an independent biological measurement, because visual similarity and similar counts are necessarily related, but it shows that the scoring calibration was transferred consistently between experiments.

The calibration experiments also show that the point at which counting becomes incomplete cannot be defined by a single numerical threshold. $P_{araBAD}$ colonies remained numerically consistent with the semilogarithmic response at 0.002% arabinose, and individual colonies containing approximately 130–140 visible papillae could still be scored. At 0.005%, however, all colonies were confluent and uncountable. Counts from colonies showing substantial overlap or diffuse central staining should be regarded as minimum estimates. The same qualification applies when an experimental perturbation increases papillation while the inducer concentration remains fixed.

$P_{araBAD}$ was generally induced with 0.002% arabinose in experiments designed to detect conditions expected to reduce adaptation. This concentration lies at the upper end of the fitted countable range; reductions in papillation move the assay towards lower densities at which individual papillae are more readily resolved.

Conversely, $P_{araMari}$ was generally induced with 0.02% arabinose when testing conditions expected to increase adaptation, including the mutant comparisons. This concentration produced a mean of 57.8 papillae, whereas the same titration remained countable at 0.05% arabinose, where the mean increased to 76.5. The $P_{araBAD}$ calibration further shows that colonies containing approximately 130–140 papillae can still be counted when the individual papillae remain spatially resolved. The standard $P_{araMari}$ condition therefore retains substantial quantitative capacity to resolve experimentally induced increases in papillation. Only responses that produce substantial merging or diffuse background staining will be progressively underestimated.

The precise density at which counting fails is not expected to be a fixed number. It will depend on papillus size and intensity, the timing and spatial distribution of adaptation events, colony morphology and the extent of diffuse background staining. In particular, heavily papillated or growth-impaired colonies sometimes developed a pale-blue central “bullseye.” This staining is morphologically distinct from the intensely blue, sharply bounded papillae. Similar pale-blue or blue-centred colonies are familiar from prolonged blue/white cloning screens. This background staining has been attributed to the cryptic *Escherichia coli* β-galactosidase locus, *ebg*, and was reported to be reduced by limiting incubation to less than 18 hours and eliminated in an *ebg*-defective host [2].

The pale bullseye was therefore treated as background rather than as a signal. Nevertheless, it further reduces the effective dynamic range because dark papillae lying within a diffuse blue region become difficult to distinguish. At high adaptation rates, this background staining and the physical confluence of genuine papillae act together to cause progressive undercounting.

Taken together, both titrations show an approximately log-linear empirical response over their nonzero, countable concentration ranges. The $P_{araBAD}$ response follows the semilogarithmic fit through 0.002% arabinose, although the colonies are crowded at this concentration; unequivocal failure of counting occurs at 0.005%, where the papillae are confluent. The $P_{araMari}$ response follows the expected semilogarithmic transfer function of the evolved sensor through 0.05% arabinose, leaving its standard 0.02% condition within a quantitatively useful region with capacity to detect increases in papillation. Where individual papillae remain resolved, the assay provides a reproducible quantitative measure of papillation. Counts should be regarded as lower-bound estimates when substantial confluence or diffuse background staining prevents individual papillae from being distinguished.

**Rare adaptive events and the limits of population averages**

CRISPR spacer acquisition is an intrinsically rare and discontinuous event. In any given bacterial population, only a small fraction of cells ever acquire a spacer, and those events are typically separated by many generations from the time at which they are detected. As with other rare adaptive or stress-associated phenotypes in bacteria—such as persistence, competence, or phase-variable gene expression—spacer acquisition is best understood as arising from transient physiological or genetic states present in a minority of cells, rather than from the average state of the population.

In such systems, population-averaged measurements can obscure the causative states that give rise to rare outcomes. This principle has been increasingly recognised in CRISPR–Cas biology, where regulation of *cas* gene expression and activity can operate heterogeneously at the single-cell level in some organisms [3]. Although the present study does not attempt to resolve single-cell mechanisms, this perspective is important for interpreting assays that report on rare adaptive events.

The papillation assay used here reflects this logic [4]. The visible blue papillae scored on day five represent the selective amplification of rare lineages that entered a permissive state earlier during colony development and subsequently expanded locally under strong selection. They do not report directly on the dominant physiological state of the colony as a whole.

**Colony microenvironmental heterogeneity**

An *E. coli* colony growing on LB agar is a strongly heterogeneous microenvironment. Most of the biomass is near stationary phase, whereas growth and transcriptional activity are concentrated in an outer annulus experiencing distinct nutrient, oxygen, and redox conditions. Cells in the colony interior are frequently stressed or nutrient limited. As a result, gene expression and physiological state may vary substantially across the spatial structure of a single colony.

This heterogeneity does not invalidate quantitative approaches, but it does complicate interpretation of bulk population measurements. In the context of the papillation assay, the biologically relevant entities are the rare lineages that ultimately give rise to papillae. Any bulk harvest from the colony surface is therefore dominated by the surrounding non-adapted population and may obscure the physiological state associated with successful adaptation events.

**Experimental reproducibility and interpretation of papillation assays**
Because papillation assays integrate rare adaptive events over several days of colony growth, absolute papillation frequencies can vary between experiments performed at different times, even when the underlying biological trends are reproducible. Such variation likely reflects sensitivity to local microenvironmental factors including agar composition, humidity, incubation conditions, and subtle differences in colony architecture.

For this reason, papillation assays are most informative when interpreted comparatively within individual experimental panels, where strains are analysed side-by-side on the same batch of plates under identical conditions. Throughout this study, conclusions are therefore drawn primarily from relative differences observed within experiments rather than from direct comparison of absolute papillation frequencies across panels generated in separate experiments.

### S1 Table. Plasmid sequences

pRC1653

aagaaaccaattgtccatattgcatcagacattgccgtcactgcgtcttttactggctcttctcgctaaccaaaccggtaaccccgcttattaaaagcatt
ctgtaacaaagcgggaccaaagccatgacaaaaacgcgtaacaaaagtgtctataatcacggcagaaaagtccacattgattatttgcacggcgt
cacactttgctatgccatagcatttttatccataagattagcggatcctacctgacgctttttatcgcaactctctactgtttctccatacccgtttttgggctaa
caggaggaattaaccatggcctggcttccccttaatcccattccactcaaagatcgcgtctccatgatctttctgcaatatgggcagatcgatgtaatag
atggcgcgtttgtacttatcgacaagacagggatccgcactcatattcctgttggctcggttgcctgcatcatgctggaacctggtacacgggtttcgcat
gcagctgtacgcctggctgcgcaagttggaacattgttggtatgggtgggggaagcgggcgttcgtgtttatgcttctggtcagcctggaggtgcgcgtt
cagataagctgctctatcaggcaaaacttgctctggatgaagatttgcgtctgaaggtcgtacgtaaaatgtttgaacttcggtttggagaacctgcgcc
tgcccggcgctccgtagagcaactcagaggtatagaaggcagtcgcgtgcgggcaacctacgcacttctggcgaagcaatacggcgtgacatgg
aatggacgtcgctacgatccgaaagactgggaaaagggcgatacgatcaaccaatgcattagcgctgcaacttcctgtttatacggcgtaactgaa
gcggcgatacttgcagctggttatgcaccagctattgggtttgtgcatacaggaaagcctctttcctttgtttacgatattgcagacatcattaaatttgaca
ctgttgtaccgaaagcttttgagatagcgcgtcgtaaccctggtgagccggaccgggaagtccgtttggcgtgcagggatatttttcgcagtagtaaaa
cattagccaaattgattccgcttatagaggacgtgcttgccgctggagaaatacaaccgccggccccacctgaagatgcacagcctgttgccattcc
gcttcctgtttcactgggagatgcaggccatcggagtagcacccatttgctgtccaccagtcatcatcaccatcaccacggatgagagaagattttca
gcctgatacagattaaatcagaacgcagaagcggtctgataaaacagaatttgcctggcggcagtagcgcggtggtcccacctgaccccatgccg
aactcagaagtgaaacgccgtagcgccgatggtagtgtggggtctccccatgcgagagtagggaactgccaggcatcaaataaaacgaaaggc
tcagtcgaaagactgggcctttcgttttatctgttgtttgtcggtgaacgctctcctgagtaggacaaatccgccgggagcggatttgaacgttgcgaag
caacggcccggagggtggcgggcaggacgcccgccataaactgccaggcatcaaattaagcagaaggccatcctgacggatggcctttttgcgt
ttctacaaactcttttgtttatttttctaaatacattcaaatatgtatccgctcatgagacaataaccctgataaatgcttcaataatattgaaaaaggaaga
gtatgagtattcaacatttccgtgtcgcccttattcccttttttgcggcattttgccttcctgtttttgctcacccagaaacgctggtgaaagtaaaagatgctg
aagatcagttgggtgcacgagtgggttacatcgaactggatctcaacagcggtaagatccttgagagttttcgccccgaagaacgttttccaatgatg
agcacttttaaagttctgctatgtggcgcggtattatcccgtgttgacgccgggcaagagcaactcggtcgccgcatacactattctcagaatgacttgg
ttgagtactcaccagtcacagaaaagcatcttacggatggcatgacagtaagagaattatgcagtgctgccataaccatgagtgataacactgcgg
ccaacttacttctgacaacgatcggaggaccgaaggagctaaccgcttttttgcacaacatgggggatcatgtaactcgccttgatcgttgggaaccg
gagctgaatgaagccataccaaacgacgagcgtgacaccacgatgcctgtagcaatggcaacaacgttgcgcaaactattaactggcgaactac
ttactctagcttcccggcaacaattaatagactggatggaggcggataaagttgcaggaccacttctgcgctcggcccttccggctggctggtttattgc
tgataaatctggagccggtgagcgtgggtctcgcggtatcattgcagcactggggccagatggtaagccctcccgtatcgtagttatctacacgacgg
ggagtcaggcaactatggatgaacgaaatagacagatcgctgagataggtgcctcactgattaagcattggtaactgtcagaccaagtttactcata
tatactttagattgatttaaaacttcatttttaatttaaaaggatctaggtgaagatcctttttgataatctcatgaccaaaatcccttaacgtgagttttcgttcc
actgagcgtcagaccccgtagaaaagatcaaaggatcttcttgagatcctttttttctgcgcgtaatctgctgcttgcaaacaaaaaaaccaccgctac
cagcggtggtttgtttgccggatcaagagctaccaactctttttccgaaggtaactggcttcagcagagcgcagataccaaatactgtccttctagtgta
gccgtagttaggccaccacttcaagaactctgtagcaccgcctacatacctcgctctgctaatcctgttaccagtggctgctgccagtggcgataagtc
gtgtcttaccgggttggactcaagacgatagttaccggataaggcgcagcggtcgggctgaacggggggttcgtgcacacagcccagcttggagc
gaacgacctacaccgaactgagatacctacagcgtgagctatgagaaagcgccacgcttcccgaagggagaaaggcggacaggtatccggta
agcggcagggtcggaacaggagagcgcacgagggagcttccagggggaaacgcctggtatctttatagtcctgtcgggtttcgccacctctgactt
gagcgtcgatttttgtgatgctcgtcaggggggcggagcctatggaaaaacgccagcaacgcggcctttttacggttcctggccttttgctggccttttgc
tcacatgttctttcctgcgttatcccctgattctgtggataaccgtattaccgcctttgagtgagctgataccgctcgccgcagccgaacgaccgagcgc
agcgagtcagtgagcgaggaagcggaagagcgcctgatgcggtattttctccttacgcatctgtgcggtatttcacaccgcatatggtgcactctcagt
acaatctgctctgatgccgcatagttaagccagtatacactccgctatcgctacgtgactgggtcatggctgcgccccgacacccgccaacacccgc
tgacgcgccctgacgggcttgtctgctcccggcatccgcttacagacaagctgtgaccgtctccgggagctgcatgtgtcagaggttttcaccgtcatc
accgaaacgcgcgaggcagcagatcaattcgcgcgcgaaggcgaagcggcatgcataatgtgcctgtcaaatggacgaagcagggattctgc
aaaccctatgctactccgtcaagccgtcaattgtctgattcgttaccaattatgacaacttgacggctacatcattcactttttcttcacaaccggcacgga
actcgctcgggctggccccggtgcatttttaaatacccgcgagaaatagagttgatcgtcaaaaccaacattgcgaccgacggtggcgataggcat
ccgggtggtgctcaaaagcagcttcgcctggctgatacgttggtcctcgcgccagcttaagacgctaatccctaactgctggcggaaaagatgtgac
agacgcgacggcgacaagcaaacatgctgtgcgacgctggcgatatcaaaattgctgtctgccaggtgatcgctgatgtactgacaagcctcgcgt
acccgattatccatcggtggatggagcgactcgttaatcgcttccatgcgccgcagtaacaattgctcaagcagatttatcgccagcagctccgaata

gcgcccttccccttgcccggcgttaatgatttgcccaaacaggtcgctgaaatgcggctggtgcgcttcatccgggcgaaagaaccccgtattggca
aatattgacggccagttaagccattcatgccagtaggcgcgcggacgaaagtaaacccactggtgataccattcgcgagcctccggatgacgacc
gtagtgatgaatctctcctggcgggaacagcaaaatatcacccggtcggcaaacaaattctcgtccctgatttttcaccaccccctgaccgcgaatgg
tgagattgagaatataacctttcattcccagcggtcggtcgataaaaaaatcgagataaccgttggcctcaatcggcgttaaacccgccaccagatg
ggcattaaacgagtatcccggcagcaggggatcattttgcgcttcagccatacttttcatactcccgccattcagag

pRC1655

aagaaaccaattgtccatattgcatcagacattgccgtcactgcgtcttttactggctcttctcgctaaccaaaccggtaaccccgcttattaaaagcatt
ctgtaacaaagcgggaccaaagccatgacaaaaacgcgtaacaaaagtgtctataatcacggcagaaaagtccacattgattatttgcacggcgt
cacactttgctatgccatagcatttttatccataagattagcggatcctacctgacgctttttatcgcaactctctactgtttctccatacccgtttttgggctaa
caggaggaattaaccatgagtatgttggtcgtggtcactgaaaatgtacctccgcgcttacgaggcagattagccatctggttgttggaggtacgtgca
ggggtatatgtaggtgatgtatccgcaaaaattcgtgaaatgatctgggaacaaatagctggactggcggaagaaggcaatgtagtgatggcatgg
gcaacgaatacggaaacgggatttgagttccagacatttgggttaaacaggcgtaccccggtagatttggatggtttaaggttggtgtctttttacctgtt
tgagcggccgcccatgggggggttctcatcatcatcatcatcatggtatggctagcatgactggtggacagcaaatgggtcgggatctgtacgacgat
gacgataaggatcgatggggatccgagctcgagatctgcagctggtaccatatgggaattcgaagcttggctgttttggcggatgagagaagattttc
agcctgatacagattaaatcagaacgcagaagcggtctgataaaacagaatttgcctggcggcagtagcgcggtggtcccacctgaccccatgcc
gaactcagaagtgaaacgccgtagcgccgatggtagtgtggggtctccccatgcgagagtagggaactgccaggcatcaaataaaacgaaagg
ctcagtcgaaagactgggcctttcgttttatctgttgtttgtcggtgaacgctctcctgagtaggacaaatccgccgggagcggatttgaacgttgcgaa
gcaacggcccggagggtggcgggcaggacgcccgccataaactgccaggcatcaaattaagcagaaggccatcctgacggatggcctttttgc
gtttctacaaactctttttgtttatttttctaaatacattcaaatatgtatccgctcatgagacaataaccctgataaatgcttcaataatattgaaaaaggaag
agtatgagtattcaacatttccgtgtcgcccttattcccttttttgcggcattttgccttcctgtttttgctcacccagaaacgctggtgaaagtaaaagatgct
gaagatcagttgggtgcacgagtgggttacatcgaactggatctcaacagcggtaagatccttgagagttttcgccccgaagaacgttttccaatgat
gagcacttttaaagttctgctatgtggcgcggtattatcccgtgttgacgccgggcaagagcaactcggtcgccgcatacactattctcagaatgacttg
gttgagtactcaccagtcacagaaaagcatcttacggatggcatgacagtaagagaattatgcagtgctgccataaccatgagtgataacactgcg
gccaacttacttctgacaacgatcggaggaccgaaggagctaaccgcttttttgcacaacatgggggatcatgtaactcgccttgatcgttgggaacc
ggagctgaatgaagccataccaaacgacgagcgtgacaccacgatgcctgtagcaatggcaacaacgttgcgcaaactattaactggcgaacta
cttactctagcttcccggcaacaattaatagactggatggaggcggataaagttgcaggaccacttctgcgctcggcccttccggctggctggtttattg
ctgataaatctggagccggtgagcgtgggtctcgcggtatcattgcagcactggggccagatggtaagccctcccgtatcgtagttatctacacgacg
gggagtcaggcaactatggatgaacgaaatagacagatcgctgagataggtgcctcactgattaagcattggtaactgtcagaccaagtttactcat
atatactttagattgatttaaaacttcatttttaatttaaaaggatctaggtgaagatcctttttgataatctcatgaccaaaatcccttaacgtgagttttcgttc
cactgagcgtcagaccccgtagaaaagatcaaaggatcttcttgagatcctttttttctgcgcgtaatctgctgcttgcaaacaaaaaaaccaccgcta
ccagcggtggtttgtttgccggatcaagagctaccaactctttttccgaaggtaactggcttcagcagagcgcagataccaaatactgtccttctagtgt
agccgtagttaggccaccacttcaagaactctgtagcaccgcctacatacctcgctctgctaatcctgttaccagtggctgctgccagtggcgataagt
cgtgtcttaccgggttggactcaagacgatagttaccggataaggcgcagcggtcgggctgaacggggggttcgtgcacacagcccagcttggag
cgaacgacctacaccgaactgagatacctacagcgtgagctatgagaaagcgccacgcttcccgaagggagaaaggcggacaggtatccggt
aagcggcagggtcggaacaggagagcgcacgagggagcttccagggggaaacgcctggtatctttatagtcctgtcgggtttcgccacctctgact
tgagcgtcgatttttgtgatgctcgtcaggggggcggagcctatggaaaaacgccagcaacgcggcctttttacggttcctggccttttgctggccttttg
ctcacatgttctttcctgcgttatcccctgattctgtggataaccgtattaccgcctttgagtgagctgataccgctcgccgcagccgaacgaccgagcg
cagcgagtcagtgagcgaggaagcggaagagcgcctgatgcggtattttctccttacgcatctgtgcggtatttcacaccgcatatggtgcactctca
gtacaatctgctctgatgccgcatagttaagccagtatacactccgctatcgctacgtgactgggtcatggctgcgccccgacacccgccaacaccc
gctgacgcgccctgacgggcttgtctgctcccggcatccgcttacagacaagctgtgaccgtctccgggagctgcatgtgtcagaggttttcaccgtc
atcaccgaaacgcgcgaggcagcagatcaattcgcgcgcgaaggcgaagcggcatgcataatgtgcctgtcaaatggacgaagcagggattct
gcaaaccctatgctactccgtcaagccgtcaattgtctgattcgttaccaattatgacaacttgacggctacatcattcactttttcttcacaaccggcacg
gaactcgctcgggctggccccggtgcatttttaaatacccgcgagaaatagagttgatcgtcaaaaccaacattgcgaccgacggtggcgatagg
catccgggtggtgctcaaaagcagcttcgcctggctgatacgttggtcctcgcgccagcttaagacgctaatccctaactgctggcggaaaagatgt
gacagacgcgacggcgacaagcaaacatgctgtgcgacgctggcgatatcaaaattgctgtctgccaggtgatcgctgatgtactgacaagcctc

gcgtacccgattatccatcggtggatggagcgactcgttaatcgcttccatgcgccgcagtaacaattgctcaagcagatttatcgccagcagctccg
aatagcgcccttccccttgcccggcgttaatgatttgcccaaacaggtcgctgaaatgcggctggtgcgcttcatccgggcgaaagaaccccgtattg
gcaaatattgacggccagttaagccattcatgccagtaggcgcgcggacgaaagtaaacccactggtgataccattcgcgagcctccggatgacg
accgtagtgatgaatctctcctggcgggaacagcaaaatatcacccggtcggcaaacaaattctcgtccctgatttttcaccacccctgaccgcga
atggtgagattgagaatataacctttcattcccagcggtcggtcgataaaaaaatcgagataaccgttggcctcaatcggcgttaaacccgccacca
gatgggcattaaacgagtatcccggcagcaggggatcattttgcgcttcagccatacttttcatactcccgccattcagag

pRC1656

aagaaaccaattgtccatattgcatcagacattgccgtcactgcgtcttttactggctcttctcgctaaccaaaccggtaaccccgcttattaaaagcatt
ctgtaacaaagcgggaccaaagccatgacaaaaacgcgtaacaaaagtgtctataatcacggcagaaaagtccacattgattatttgcacggcgt
cacactttgctatgccatagcatttttatccataagattagcggatcctacctgacgctttttatcgcaactctctactgtttctccatacccgtttttgggctaa
caggaggaattaaccatggcctggcttccccttaatcccattccactcaaagatcgcgtctccatgatctttctgcaatatgggcagatcgatgtaatag
atggcgcgtttgtacttatcgacaagacagggatccgcactcatattcctgttggctcggttgcctgcatcatgctggaacctggtacacgggtttcgcat
gcagctgtacgcctggctgcgcaagttggaacattgttggtatgggtgggggaagcgggcgttcgtgtttatgcttctggtcagcctggaggtgcgcgtt
cagataagctgctctatcaggcaaaacttgctctggatgaagatttgcgtctgaaggtcgtacgtaaaatgtttgaacttcggtttggagaacctgcgcc
tgcccggcgctccgtagagcaactcagaggtatagaaggcagtcgcgtgcgggcaacctacgcacttctggcgaagcaatacggcgtgacatgg
aatggacgtcgctacgatccgaaagactgggaaaagggcgatacgatcaaccaatgcattagcgctgcaacttcctgtttatacggcgtaactgaa
gcggcgatacttgcagctggttatgcaccagctattgggtttgtgcatacaggaaagcctctttcctttgtttacgatattgcagacatcattaaatttgaca
ctgttgtaccgaaagcttttgagatagcgcgtcgtaaccctggtgagccggaccgggaagtccgtttggcgtgcagggatatttttcgcagtagtaaaa
cattagccaaattgattccgcttatagaggacgtgcttgccgctggagaaatacaaccgccggccccacctgaagatgcacagcctgttgccattcc
gcttcctgtttcactgggagatgcaggccatcggagtagctgaaatgagtatgttggtcgtggtcactgaaaatgtacctccgcgcttacgaggcagat
tagccatctggttgttggaggtacgtgcaggggtatatgtaggtgatgtatccgcaaaaattcgtgaaatgatctgggaacaaatagctggactggcg
gaagaaggcaatgtagtgatggcatgggcaacgaatacggaaacgggatttgagttccagacatttgggttaaacaggcgtaccccggtagatttg
gatggtttaaggttggtgtctttttacctgtttgagcggccgcccatgggggttctcatcatcatcatcatcatggtatggctagcatgactggtggacag
caaatgggtcgggatctgtacgacgatgacgataaggatcgatggggatccgagctcgagatctgcagctggtaccatatgggaattcgaagcttg
gctgttttggcggatgagagaagattttcagcctgatacagattaaatcagaacgcagaagcggtctgataaaacagaatttgcctggcggcagtag
cgcggtggtcccacctgaccccatgccgaactcagaagtgaaacgccgtagcgccgatggtagtgtggggtctccccatgcgagagtagggaac
tgccaggcatcaaataaaacgaaaggctcagtcgaaagactgggcctttcgttttatctgttgtttgtcggtgaacgctctcctgagtaggacaaatcc
gccgggagcggatttgaacgttgcgaagcaacggcccggagggtggcgggcaggacgcccgccataaactgccaggcatcaaattaagcaga
aggccatcctgacggatggcctttttgcgtttctacaaactcttgtctcaaaatctctgatgttacattgcacaagataaaaatatatcatcatgaacaata
aaactgtctgcttacataaacagtaatacaaggggtgttatgagccatattcaacgggaaacgtcttgctcgaggccgcgattaaattccaacatgga
tgctgatttatatgggtataaatgggctcgcgataatgtcgggcaatcaggtgcgacaatctatcgattgtatgggaagcccgatgcgccagagttgttt
ctgaaacatggcaaaggtagcgttgccaatgatgttacagatgagatggtcagactaaactggctgacggaatttatgcctcttccgaccatcaagc
attttatccgtactcctgatgatgcatggttactcaccactgcgatccccgggaaaacagcattccaggtattagaagaatatcctgattcaggtgaaa
atattgttgatgcgctggcagtgttcctgcgccggttgcattcgattcctgtttgtaattgtccttttaacagcgatcgcgtatttcgtctcgctcaggcgcaat
cacgaatgaataacggtttggttgatgcgagtgattttgatgacgagcgtaatggctggcctgttgaacaagtctggaaagaaatgcataagcttttgc
cattctcaccggattcagtcgtcactcatggtgatttctcacttgataaccttatttttgacgaggggaaattaataggttgtattgatgttggacgagtcgg
aatcgcagaccgataccaggatcttgccatcctatggaactgcctcggtgagttttctccttcattacagaaacggctttttcaaaaatatggtattgata
atcctgatatgaataaattgcagtttcatttgatgctcgatgagtttttctaactgtcagaccaagtttactcatatatactttagattgatttaaaacttcatttt
aatttaaaaggatctaggtgaagatcctttttgataatctcatgaccaaaatcccttaacgtgagttttcgttccactgagcgtcagaccccgtagaaaa
gatcaaaggatcttcttgagatcctttttttctgcgcgtaatctgctgcttgcaaacaaaaaaaccaccgctaccagcggtggtttgtttgccggatcaag
agctaccaactctttttccgaaggtaactggcttcagcagagcgcagataccaaatactgtccttctagtgtagccgtagttaggccaccacttcaaga
actctgtagcaccgcctacatacctcgctctgctaatcctgttaccagtggctgctgccagtggcgataagtcgtgtcttaccgggttggactcaagacg
atagttaccggataaggcgcagcggtcgggctgaacggggggttcgtgcacacagcccagcttggagcgaacgacctacaccgaactgagata
cctacagcgtgagctatgagaaagcgccacgcttcccgaagggagaaaggcggacaggtatccggtaagcggcagggtcggaacaggagag
cgcacgagggagcttccagggggaaacgcctggtatctttatagtcctgtcgggtttcgccacctctgacttgagcgtcgatttttgtgatgctcgtcagg

ggggcggagcctatggaaaaacgccagcaacgcggcctttttacggttcctggccttttgctggccttttgctcacatgttctttcctgcgttatcccctgat
tctgtggataaccgtattaccgcctttgagtgagctgataccgctcgccgcagccgaacgaccgagcgcagcgagtcagtgagcgaggaagcgg
aagagcgcctgatgcggtattttctccttacgcatctgtgcggtatttcacaccgcatatggtgcactctcagtacaatctgctctgatgccgcatagttaa
gccagtatacactccgctatcgctacgtgactgggtcatggctgcgccccgacacccgccaacacccgctgacgcgccctgacgggcttgtctgctc
ccggcatccgcttacagacaagctgtgaccgtctccgggagctgcatgtgtcagaggttttcaccgtcatcaccgaaacgcgcgaggcagcagatc
aattcgcgcgcgaaggcgaagcggcatgcataatgtgcctgtcaaatggacgaagcagggattctgcaaaccctatgctactccgtcaagccgtc
aattgtctgattcgttaccaattatgacaacttgacggctacatcattcactttttcttcacaaccggcacggaactcgctcgggctggccccggtgcatttt
ttaaatacccgcgagaaatagagttgatcgtcaaaaccaacattgcgaccgacggtggcgataggcatccgggtggtgctcaaaagcagcttcgc
ctggctgatacgttggtcctcgcgccagcttaagacgctaatccctaactgctggcggaaaagatgtgacagacgcgacggcgacaagcaaacat
gctgtgcgacgctggcgatatcaaaattgctgtctgccaggtgatcgctgatgtactgacaagcctcgcgtacccgattatccatcggtggatggagc
gactcgttaatcgcttccatgcgccgcagtaacaattgctcaagcagatttatcgccagcagctccgaatagcgcccttccccttgcccggcgttaatg
atttgcccaaacaggtcgctgaaatgcggctggtgcgcttcatccgggcgaaagaaccccgtattggcaaatattgacggccagttaagccattcat
gccagtaggcgcgcggacgaaagtaaacccactggtgataccattcgcgagcctccggatgacgaccgtagtgatgaatctctcctggcgggaa
cagcaaaatatcacccggtcggcaaacaaattctcgtccctgatttttcaccaccccctgaccgcgaatggtgagattgagaatataacctttcattcc
cagcggtcggtcgataaaaaaatcgagataaccgttggcctcaatcggcgttaaacccgccaccagatgggcattaaacgagtatcccggcagc
aggggatcattttgcgcttcagccatacttttcatactcccgccattcagag

pRC1657

aagaaaccaattgtccatattgcatcagacattgccgtcactgcgtcttttactggctcttctcgctaaccaaaccggtaaccccgcttattaaaagcatt
ctgtaacaaagcgggaccaaagccatgacaaaaacgcgtaacaaaagtgtctataatcacggcagaaaagtccacattgattatttgcacggcgt
cacactttgctatgccatagcatttttatccataagattagcggatcctacctgacgctttttatcgcaactctctactgtttctccatacccgtttttgggctaa
caggaggaattaaccatggcctggcttccccttaatcccattccactcaaagatcgcgtctccatgatctttctgcaatatgggcagatcgatgtaatag
atggcgcgtttgtacttatcgacaagacagggatccgcactcatattcctgttggctcggttgcctgcatcatgctggaacctggtacacgggtttcgcat
gcagctgtacgcctggctgcgcaagttggaacattgttggtatgggtgggggaagcgggcgttcgtgtttatgcttctggtcagcctggaggtgcgcgtt
cagataagctgctctatcaggcaaaacttgctctggatgaagatttgcgtctgaaggtcgtacgtaaaatgtttgaacttcggtttggagaacctgcgcc
tgcccggcgctccgtagagcaactcagaggtatagaaggcagtcgcgtgcgggcaacctacgcacttctggcgaagcaatacggcgtgacatgg
aatggacgtcgctacgatccgaaagactgggaaaagggcgatacgatcaaccaatgcattagcgctgcaacttcctgtttatacggcgtaactgaa
gcggcgatacttgcagctggttatgcaccagctattgggtttgtgcatacaggaaagcctctttcctttgtttacgatattgcagacatcattaaatttgaca
ctgttgtaccgaaagcttttgagatagcgcgtcgtaaccctggtgagccggaccgggaagtccgtttggcgtgcagggatatttttcgcagtagtaaaa
cattagccaaattgattccgcttatagaggacgtgcttgccgctggagaaatacaaccgccggccccacctgaagatgcacagcctgttgccattcc
gcttcctgtttcactgggagatgcaggccatcggagtagctgaaatgagtatgttggtcgtggtcactgaaaatgtacctccgcgcttacgaggcagat
tagccatctggttgttggaggtacgtgcaggggtatatgtaggtgatgtatccgcaaaaattcgtgaaatgatctgggaacaaatagctggactggcg
gaagaaggcaatgtagtgatggcatgggcaacgaatacggaaacgggatttgagttccagacatttgggttaaacaggcgtaccccggtagatttg
gatggtttaaggttggtgtctttttttacctgtttgagcggccgcccatgggggggttctcatcatcatcatcatcatggtatggctagcatgactggtggacag
caaatgggtcgggatctgtacgacgatgacgataaggatcgatggggatccgagctcgagatctgcagctggtaccatatgggaattcgaagcttg
gctgttttggcggatgagagaagattttcagcctgatacagattaaatcagaacgcagaagcggtctgataaaacagaatttgcctggcggcagtag
cgcggtggtcccacctgaccccatgccgaactcagaagtgaaacgccgtagcgccgatggtagtgtggggtctccccatgcgagagtagggaac
tgccaggcatcaaataaaacgaaaggctcagtcgaaagactgggcctttcgttttatctgttgtttgtcggtgaacgctctcctgagtaggacaaatcc
gccgggagcggatttgaacgttgcgaagcaacggcccggagggtggcgggcaggacgcccgccataaactgccaggcatcaaattaagcaga
aggccatcctgacggatggcctttttgcgtttctacaaactctttacgccccgccctgccactcatcgcagtactgttgtattcattaagcatctgccgaca
tggaagccatcacaaacggcatgatgaacctgaatcgccagcggcatcagcaccttgtcgccttgcgtataatatttgcccatggtgaaaacgggg
gcgaagaagttgtccatattggccacgtttaaatcaaaactggtgaaactcacccagggattggctgagacgaaaaacatattctcaataaaccctttt
agggaaataggccaggttttcaccgtaacacgccacatcttgcgaatatatgtgtagaaactgccggaaatcgtcgtggtattcactccagagcgat
gaaaacgtttcagtttgctcatggaaaacggtgtaacaagggtgaacactatcccatatcaccagctcaccgtctttcattgccatacgtaattccggat
gagcattcatcaggcgggcaagaatgtgaataaaggccggataaaacttgtgcttatttttctttacggtctttaaaaaggccgtaatatccagctgaa
cggtctggttataggtacattgagcaactgactgaaatgcctcaaaatgttctttacgatgccattgggatatatcaacggtggtatatccagtgattttttt

ctccattttagcttccttagctcctgaaaatctcgacaactcaaaaaatacgcccggtagtgatcttatttcattatggtgaaagttggaacctcttacgtgc
cgatcactgtcagaccaagtttactcatatatactttagattgatttaaaacttcatttttaatttaaaaggatctaggtgaagatcctttttgataatctcatga
ccaaaatcccttaacgtgagttttcgttccactgagcgtcagaccccgtagaaaagatcaaaggatcttcttgagatcctttttttctgcgcgtaatctgct
gcttgcaaacaaaaaaaccaccgctaccagcggtggtttgtttgccggatcaagagctaccaactctttttccgaaggtaactggcttcagcagagc
gcagataccaaatactgtccttctagtgtagccgtagttaggccaccacttcaagaactctgtagcaccgcctacatacctcgctctgctaatcctgtta
ccagtggctgctgccagtggcgataagtcgtgtcttaccgggttggactcaagacgatagttaccggataaggcgcagcggtcgggctgaacggg
gggttcgtgcacacagcccagcttggagcgaacgacctacaccgaactgagatacctacagcgtgagctatgagaaagcgccacgcttcccgaa
gggagaaaggcggacaggtatccggtaagcggcagggtcggaacaggagagcgcacgagggagcttccagggggaaacgcctggtatcttt
atagtcctgtcgggtttcgccacctctgacttgagcgtcgatttttgtgatgctcgtcaggggggcggagcctatggaaaaacgccagcaacgcggcc
tttttacggttcctggccttttgctggccttttgctcacatgttctttcctgcgttatcccctgattctgtggataaccgtattaccgcctttgagtgagctgatacc
gctcgccgcagccgaacgaccgagcgcagcgagtcagtgagcgaggaagcggaagagcgcctgatgcggtattttctccttacgcatctgtgcg
gtatttcacaccgcatatggtgcactctcagtacaatctgctctgatgccgcatagttaagccagtatacactccgctatcgctacgtgactgggtcatg
gctgcgccccgacacccgccaacacccgctgacgcgccctgacgggcttgtctgctcccggcatccgcttacagacaagctgtgaccgtctccgg
gagctgcatgtgtcagaggttttcaccgtcatcaccgaaacgcgcgaggcagcagatcaattcgcgcgcgaaggcgaagcggcatgcataatgtg
cctgtcaaatggacgaagcagggattctgcaaaccctatgctactccgtcaagccgtcaattgtctgattcgttaccaattatgacaacttgacggcta
catcattcactttttcttcacaaccggcacggaactcgctcgggctggccccggtgcattttttaaatacccgcgagaaatagagttgatcgtcaaaacc
aacattgcgaccgacggtggcgataggcatccgggtggtgctcaaaagcagcttcgcctggctgatacgttggtcctcgcgccagcttaagacgct
aatccctaactgctggcggaaaagatgtgacagacgcgacggcgacaagcaaacatgctgtgcgacgctggcgatatcaaaattgctgtctgcca
ggtgatcgctgatgtactgacaagcctcgcgtacccgattatccatcggtggatggagcgactcgttaatcgcttccatgcgccgcagtaacaattgct
caagcagatttatcgccagcagctccgaatagcgcccttccccttgcccggcgttaatgatttgcccaaacaggtcgctgaaatgcggctggtgcgct
tcatccgggcgaaagaaccccgtattggcaaatattgacggccagttaagccattcatgccagtaggcgcgcggacgaaagtaaacccactggtg
ataccattcgcgagcctccggatgacgaccgtagtgatgaatctctcctggcgggaacagcaaaatatcacccggtcggcaaacaaattctcgtcc
ctgatttttcaccaccccctgaccgcgaatggtgagattgagaatataacctttcattcccagcggtcggtcgataaaaaaatcgagataaccgttggc
ctcaatcggcgttaaacccgccaccagatgggcattaaacgagtatcccggcagcaggggatcattttgcgcttcagccatacttttcatactcccgc
cattcagag

pRC1658

aagaaaccaattgtccatattgcatcagacattgccgtcactgcgtcttttactggctcttctcgctaaccaaaccggtaaccccgcttattaaaagcatt
ctgtaacaaagcgggaccaaagccatgacaaaaacgcgtaacaaaagtgtctataatcacggcagaaaagtccacattgattatttgcacggcgt
cacactttgctatgccatagcatttttatccataagattagcggatcctacctgacgctttttatcgcaactctctactgtttctccatacccgtttttttgggctaa
caggaggaattaaccatggcctggcttccccttaatcccattccactcaaagatcgcgtctccatgatctttctgcaatatgggcagatcgatgtaatag
atggcgcgtttgtacttatcgacaagacagggatccgcactcatattcctgttggctcggttgcctgcatcatgctggaacctggtacacgggtttcgcat
gcagctgtacgcctggctgcgcaagttggaacattgttggtatgggtgggggaagcgggcgttcgtgtttatgcttctggtcagcctggaggtgcgcgtt
cagataagctgctctatcaggcaaaacttgctctggatgaagatttgcgtctgaaggtcgtacgtaaaatgtttgaacttcggtttggagaacctgcgcc
tgcccggcgctccgtagagcaactcagaggtatagaaggcagtcgcgtgcgggcaacctacgcacttctggcgaagcaatacggcgtgacatgg
aatggacgtcgctacgatccgaaagactgggaaaagggcgatacgatcaaccaatgcattagcgctgcaacttcctgtttatacggcgtaactgaa
gcggcgatacttgcagctggttatgcaccagctattgggtttgtgcatacaggaaagcctctttcctttgtttacgatattgcagccatcattaaatttgaca
ctgttgtaccgaaagcttttgagatagcgcgtcgtaaccctggtgagccggaccgggaagtccgtttggcgtgcagggatatttttcgcagtagtaaaa
cattagccaaattgattccgcttatagaggacgtgcttgccgctggagaaatacaaccgccggccccacctgaagatgcacagcctgttgccattcc
gcttcctgtttcactgggagatgcaggccatcggagtagctgaaatgagtatgttggtcgtggtcactgaaaatgtacctccgcgcttacgaggcagat
tagccatctggttgttggaggtacgtgcaggggtatatgtaggtgatgtatccgcaaaaattcgtgaaatgatctgggaacaaatagctggactggcg
gaagaaggcaatgtagtgatggcatgggcaacgaatacggaaacgggatttgagttccagacatttgggttaaacaggcgtaccccggtagatttg
gatggtttaaggttggtgtctttttttacctgtttgagcggccgcccatgggggttctcatcatcatcatcatcatggtatggctagcatgactggtggacag
caaatgggtcgggatctgtacgacgatgacgataaggatcgatggggatccgagctcgagatctgcagctggtaccatatgggaattcgaagcttg
gctgttttggcggatgagagaagattttcagcctgatacagattaaatcagaacgcagaagcggtctgataaaacagaatttgcctggcggcagtag
cgcggtggtcccacctgaccccatgccgaactcagaagtgaaacgccgtagcgccgatggtagtgtggggtctccccatgcgagagtagggaac

tgccaggcatcaaataaaacgaaaggctcagtcgaaagactgggcctttcgttttatctgttgtttgtcggtgaacgctctcctgagtaggacaaatcc
gccgggagcggatttgaacgttgcgaagcaacggcccggagggtggcgggcaggacgcccgccataaactgccaggcatcaaattaagcaga
aggccatcctgacggatggcctttttgcgtttctacaaactcttgtctcaaaatctctgatgttacattgcacaagataaaaatatatcatcatgaacaata
aaactgtctgcttacataaacagtaatacaaggggtgttatgagccatattcaacgggaaacgtcttgctcgaggccgcgattaaattccaacatgga
tgctgatttatatgggtataaatgggctcgcgataatgtcgggcaatcaggtgcgacaatctatcgattgtatgggaagcccgatgcgccagagttgttt
ctgaaacatggcaaaggtagcgttgccaatgatgttacagatgagatggtcagactaaactggctgacggaatttatgcctcttccgaccatcaagc
attttatccgtactcctgatgatgcatggttactcaccactgcgatccccgggaaaacagcattccaggtattagaagaatatcctgattcaggtgaaa
atattgttgatgcgctggcagtgttcctgcgccggttgcattcgattcctgtttgtaattgtccttttaacagcgatcgcgtatttcgtctcgctcaggcgcaat
cacgaatgaataacggtttggttgatgcgagtgattttgatgacgagcgtaatggctggcctgttgaacaagtctggaaagaaatgcataagcttttgc
cattctcaccggattcagtcgtcactcatggtgatttctcacttgataaccttatttttgacgaggggaaattaataggttgtattgatgttggacgagtcgg
aatcgcagaccgataccaggatcttgccatcctatggaactgcctcggtgagttttctccttcattacagaaacggctttttcaaaaatatggtattgata
atcctgatatgaataaattgcagtttcatttgatgctcgatgagtttttctaactgtcagaccaagtttactcatatatactttagattgatttaaaacttcatttt
aatttaaaaggatctaggtgaagatcctttttgataatctcatgaccaaaatcccttaacgtgagttttcgttccactgagcgtcagaccccgtagaaaa
gatcaaaggatcttcttgagatcctttttttctgcgcgtaatctgctgcttgcaaacaaaaaaaccaccgctaccagcggtggtttgtttgccggatcaag
agctaccaactctttttccgaaggtaactggcttcagcagagcgcagataccaaatactgtccttctagtgtagccgtagttaggccaccacttcaaga
actctgtagcaccgcctacatacctcgctctgctaatcctgttaccagtggctgctgccagtggcgataagtcgtgtcttaccgggttggactcaagacg
atagttaccggataaggcgcagcggtcgggctgaacggggggttcgtgcacacagcccagcttggagcgaacgacctacaccgaactgagata
cctacagcgtgagctatgagaaagcgccacgcttcccgaagggagaaaggcggacaggtatccggtaagcggcagggtcggaacaggagag
cgcacgagggagcttccagggggaaacgcctggtatctttatagtcctgtcgggtttcgccacctctgacttgagcgtcgatttttgtgatgctcgtcagg
ggggcggagcctatggaaaaacgccagcaacgcggcctttttacggttcctggccttttgctggccttttgctcacatgttctttcctgcgttatcccctgat
tctgtggataaccgtattaccgcctttgagtgagctgataccgctcgccgcagccgaacgaccgagcgcagcgagtcagtgagcgaggaagcgg
aagagcgcctgatgcggtattttctccttacgcatctgtgcggtatttcacaccgcatatggtgcactctcagtacaatctgctctgatgccgcatagttaa
gccagtatacactccgctatcgctacgtgactgggtcatggctgcgccccgacacccgccaacacccgctgacgcgccctgacgggcttgtctgctc
ccggcatccgcttacagacaagctgtgaccgtctccgggagctgcatgtgtcagaggttttcaccgtcatcaccgaaacgcgcgaggcagcagatc
aattcgcgcgcgaaggcgaagcggcatgcataatgtgcctgtcaaatggacgaagcagggattctgcaaaccctatgctactccgtcaagccgtc
aattgtctgattcgttaccaattatgacaacttgacggctacatcattcactttttcttcacaaccggcacggaactcgctcgggctggccccggtgcatttt
ttaaatacccgcgagaaatagagttgatcgtcaaaaccaacattgcgaccgacggtggcgataggcatccgggtggtgctcaaaagcagcttcgc
ctggctgatacgttggtcctcgcgccagcttaagacgctaatccctaactgctggcggaaaagatgtgacagacgcgacggcgacaagcaaacat
gctgtgcgacgctggcgatatcaaaattgctgtctgccaggtgatcgctgatgtactgacaagcctcgcgtacccgattatccatcggtggatggagc
gactcgttaatcgcttccatgcgccgcagtaacaattgctcaagcagatttatcgccagcagctccgaatagcgcccttccccttgcccggcgttaatg
atttgcccaaacaggtcgctgaaatgcggctggtgcgcttcatccgggcgaaagaaccccgtattggcaaatattgacggccagttaagccattcat
gccagtaggcgcgcggacgaaagtaaacccactggtgataccattcgcgagcctccggatgacgaccgtagtgatgaatctctcctggcgggaa
cagcaaaatatcacccggtcggcaaacaaattctcgtccctgatttttcaccaccccctgaccgcgaatggtgagattgagaatataacctttcattcc
cagcggtcggtcgataaaaaaatcgagataaccgttggcctcaatcggcgttaaacccgccaccagatgggcattaaacgagtatcccggcagc
aggggatcattttgcgcttcagccatacttttcatactcccgccattcagag

pRC2747

ctcggtaccaaattccagaaaagaggcctcccgaaagggggcctttttcgttttggtccaatggcggcgcgccatcgaatggtgcaaaacctttcg
cggtatggcatgatagcgcccggaagagagtcaattcagggtggtgaatatggctgaagcgcaaaatgatcccctgctgccgggatactcgtttaat
gcccatctggtggcgggtttaacgccgattgaggccaacggttatctcgatttttttatcgaccgaccgctgggaatgaaaggttatattctcaatctcac
cattcgcggtcagggggtggtgaaaaatcagggacgagaatttgtttgccgaccgggtgatattttgctgttcccgccaggagagattcatcactacg
gtcgtcatccggaggctcgcgaatggtatcaccagtgggtttactttcgtccgcgcgcctactggcatgaatggcttaactggccgtcaatatttgccaa
tacggggttctttcgcccggatgaagcgcaccagccgcatttcagcgactttttgggcaaatcattaacgccggcaaggggaagggcgctattcg
gagctgctggcgataaatctgcttgagcaattgttactgcggcgcatgctagcgattaacggatcgctccatccaccgatggataatcgggtacgcga
ggcttgtcagtacatcagcgatcacctggcagacagcaattttgatatcgccagcgtcgcacagcatgtttgcttgtcgccgtcgcgtctgtcacatctttt
ccgccagcagttagggattagcgtcttaagctggcgcgaggaccaacgtatcagccaggcgaagctgcttttgagcaccacccggatgcctatcgc

caccgtcggtcgcaatgttggttttgacgatcaactctatttctcgcgggtatttaaaaaatgcaccggggccagcccgagcgagttccgtgccggtttg
gaagaaaaagtgaatgatgtagccgtcaagttgtcatgataagatcctattccagcgggattaaagaggagcgattaagcatggttactatcaatac
ggaatctgctttaacgccacgttctttgcgggatacgcggcgtatgaatatgtttgtttcggtagctgctgcggtcgcaggattgttatttggtcttgatatcg
gcgtaatcgccggagcgttgccgttcattaccgatcactttgtgctgaccagtcgtttgcaggaatgggtggttagtagcatgatgctcggtgcagcaat
tggtgcgctgtttaatggttggctgtcgttccgcctggggcgtaaatacagcctgatggcgggggccatcctgtttgtactcggttctatagggtccgctttt
gcgaccagcgtagagatgttaatcgccgctcgtgtggtgctgggcattgctgtcgggatcgcgtcttacaccgctcctctgtatctttctgaaatggcaa
gtgaaaacgttcgcggtaagatgatcagtatgtaccagttgatggtcacactcggcatcgtgctggcgtttttatccgatacagcgttcagttatagcggt
aactggcgcgcaatgttgggggttcttgctttaccagcagttctgctgattattctggtagtatttctgccaaatagcccgcgctggctggcggaaaagg
ggcgtcatattgaggcggaagaagtattgcgtatgctgcgcgatacgtcggaaaaagcgcgagaagaactcaacgaaattcgtgaaagcctgaa
gttaaaacagggcggttgggcactgtttaagatcaaccgtaacgtccgtcgtgctgtgtttctcggtatgttgttgcaggcgatgcagcagtttaccggta
tgaacatcatcatgtactacgcgccgcgtatcttcaaaatggcgggctttacgaccacagaacaacagatgattgcgactctggtcgtagggctgac
ctttatgttcgccacctttattgcggtgtttacggtagataaagcagggcgtaaaccggctctgaaaattggtttcagcgtgatggcgttaggcactctggt
gctgggctattgcctgatgcagtttgataacggtacggcttccagtggcttgtcctggctctctgttggcatgacgatgatgtgtattgccggttatgcgatg
agcgccgcgccagtggtgtggatcctgtgctctgaaattcagccgctgaaatgccgcgatttcggtattacctgttcgaccaccacgaactgggtgtc
gaatatgattatcggcgcgaccttcctgacactgcttgatagcattggcgctgccggtacgttctggctctacactgcgctgaacattgcgtttgtgggca
ttactttctggctcattccggaaaccaaaaatgtcacgctggaacatatcgaacgcaaactgatggcaggcgagaagttgagaaatatcggcgtctg
ataaggatcctaattggtaacgaatcagacaattgacggctcgagggagtagcatagggtttgcagaatccctgcttcgtccatttgacaggcacatt
atgcatcgatgataagctgtcaaacatgagcagatcctctacgccggacgcatcgtggccggcatcaccggcgccacaggtgcggttgctggcgc
ctatatcgccgacatcaccgatggggaagatcgggctcgccacttcgggctcatgagcaaatattttatctgaggtgcttcctcgctcactgactcgctg
cacgaggcagacctcagcgctagcggagtgtatactggcttactatgttggcactgatgagggtgtcagtgaagtgcttcatgtggcaggagaaaa
aaggctgcaccggtgcgtcagcagaatatgtgatacaggatatattccgcttcctcgctcactgactcgctacgctcggtcgttcgactgcggcgagc
ggaaatggcttacgaacggggcggagatttcctggaagatgccaggaagatacttaacagggaagtgagagggccgcggcaaagccgtttttcc
ataggctccgcccccctgacaagcatcacgaaatctgacgctcaaatcagtggtggcgaaacccgacaggactataaagataccaggcgtttccc
cctggcggctccctcgtgcgctctcctgttcctgcctttcggtttaccggtgtcattccgctgttatggccgcgtttgtctcattccacgcctgacactcagttc
cgggtaggcagttcgctccaagctggactgtatgcacgaaccccccgttcagtccgaccgctgcgccttatccggtaactatcgtcttgagtccaacc
cggaaagacatgcaaaagcaccactggcagcagccactggtaattgatttagaggagttagtcttgaagtcatgcgccggttaaggctaaactgaa
aggacaagttttggtgactgcgctcctccaagccagttacctcggttcaaagagttggtagctcagagaaccttcgaaaaaccgccctgcaaggcg
gttttttcgttttcagagcaagagattacgcgcagaccaaaacgatctcaagaagatcatcttattaaggggtctgacgctcagtggaacgaaaaatc
aatctaaagtatatatgagtaaacttggtctgacagttaccttagaaaaactcatcgagcatcaaatgaaactgcaatttattcatatcaggattatcaat
accatatttttgaaaaagccgtttctgtaatgaaggagaaaactcaccgaggcagttccataggatggcaagatcctggtatcggtctgcgattccga
ctcgtccaacatcaatacaacctattaatttcccctcgtcaaaaataaggttatcaagtgagaaatcaccatgagtgacgactgaatccggtgagaat
ggcaaaagcttatgcatttctttccagacttgttcaacaggccagccattacgctcgtcatcaaaatcactcgcatcaaccaaaccgttattcattcgtga
ttgcgcctgagcgagacgaaatacgcgatcgctgttaaaaggacaattacaaacaggaatcgaatgcaaccggcgcaggaacactgccagcgc
atcaacaatattttcacctgaatcaggatattcttctaatacctggaatgctgttttcccggggatcgcagtggtgagtaaccatgcatcatcaggagtac
ggataaaatgcttgatggtcggaagaggcataaattccgtcagccagtttagtctgaccatctcatctgtaacatcattggcaacgctacctttgccatg
tttcagaaacaactctggcgcatcgggcttcccatacaatcgatagattgtcgcacctgattgcccgacattatcgcgagcccatttatacccatataaa
tcagcatccatgttggaatttaatcgcggcctcgagcaagacgtttcccgttgaatatggctcataacaccccttgtattactgtttatgtaagcagacagt
tttattgttcatgatgatatatttttatcttgtgcaatgtacatcagagattttgagacacaaccaattattgaaggcctccctaacgggggccttttttgtttc
tggtctcccgcttaacgatcgttggctgagaaaccaattgtccatattgcatcagacattgccgtcactgcgtcttttactggctcttctcgctaaccaaac
cggtaaccccgcttattaaaagcattctgtaacaaagcgggaccaaagccatgacaaaaacgcgtaacaaaagtgtctataatcacggcagaaa
agtccacattgattatttgcacggcgtcacactttgctatgccatagcatttttatccataagattagcggatcctacctgacgctttttatcgcaactctcta
ctgtttctccatacccgagctgtcaccggatgtgctttccggtctgatgagtccgtgaggacgaaacagcctctacaaataattttgtttaatactagaga
aagaggggaaatactagatggcctggcttccccttaatcccattccactcaaagatcgcgtctccatgatctttctgcaatatgggcagatcgatgtaat
agatggcgcgtttgtacttatcgacaagacagggatccgcactcatattcctgttggctcggttgcctgcatcatgctggaacctggtacacgggtttcg
catgcagctgtacgcctggctgcgcaagttggaacattgttggtatgggtgggggaagcgggcgttcgtgtttatgcttctggtcagcctggaggtgcg
cgttcagataagctgctctatcaggcaaaacttgctctggatgaagatttgcgtctgaaggtcgtacgtaaaatgtttgaacttcggtttggagaacctgc
gcctgcccggcgctccgtagagcaactcagaggtatagaaggcagtcgcgtgcgggcaacctacgcacttctggcgaagcaatacggcgtgac
atggaatggacgtcgctacgatccgaaagactgggaaaagggcgatacgatcaaccaatgcattagcgctgcaacttcctgtttatacggcgtaac

tgaagcggcgatacttgcagctggttatgcaccagctattgggtttgtgcatacaggaaagcctctttcctttgtttacgatattgcagacatcattaaattt
gacactgttgtaccgaaagcttttgagatagcgcgtcgtaaccctggtgagccggaccgggaagtccgtttggcgtgcagggatatttttcgcagtagt
aaaacattagccaaattgattccgcttatagaggacgtgcttgccgctggagaaatacaaccgccggccccacctgaagatgcacagcctgttgcc
attccgcttcctgtttcactgggagatgcaggccatcggagtagctgaaatgagtatgttggtcgtggtcactgaaaatgtacctccgcgcttacgagg
cagattagccatctggttgttggaggtacgtgcaggggtatatgtaggtgatgtatccgcaaaaattcgtgaaatgatctgggaacaaatagctggact
ggcggaagaaggcaatgtagtgatggcatgggcaacgaatacggaaacgggatttgagttccagacatttgggttaaacaggcgtaccccggta
gatttggatggtttaaggttggtgtctttttacctgtttgagcgg

pRC2781

ctgtcaccggatgtgctttccggtctgatgagtccgtgaggacgaaacagcctctacaaataattttgtttaatactagagaaagaggggaaatacta
gatgtacacttcaggctatgcacatcgttcttcgtcgttctcatccgcagcaagtaaaattgcgcgtgtctctacggaaaacactacagccgggcttatc
agtgaagttgtctatcgcgaagatcagcccatgatgacgcaacttctactgttgccattgttacagcaactcggtcagcaatcgcgctggcaactctgg
ttaacaccgcaacaaaaactgagtcgggaatgggttcaggcatctgggctacccttaacgaaagtaatgcagattagccagctctccccttgccac
actgtggagtcaatggttcgcgctttacgcacgggcaattacagtgtggtgatcggttggttggcagatgatttgactgaagaagagcatgctgaactt
gttgatgcggcaaatgaaggtaacgctatggggtttattatgcgtccggtaagcgcatcctctcacgccacgagacaactttccgggctaaaaattca
ctctaatttgtatcattaataactcggtaccaaattccagaaaagaggcctcccgaaagggggcctttttcgttttggtccaatggcggcgcgccatc
gaatggtgcaaaacctttcgcggtatggcatgatagcgcccggaagagagtcaattcagggggtgaataatggacatgcctcgtattaaaccgg
gtcagcgtgttatgatggcactgcgtaaaatgattgcaagcggtgaaatcaaaagtggtgaacgtattgcagaaattccgaccgcagcagcactgg
gtgttagccgtatgccggttcgtatcgcactgcgttcactggaacaagaaggtctggttgttcgtctgggtgcacgtggttatgcagcccgtggtgttagc
agcgatcagattcgtgatgcaattgaagttcgtggtgttctggaaggttttgcagcacgtcgtctggcagaacgtggtatgaccgcagaaacccatgc
acgttttgttgtactgattgcagaaggtgaagcactgtttgcagccggtcgcctgaatggtgaagatctggatcgttatgccgcatataatcaggcatttc
atgataccctggttagcgcagcaggtaatggtgcagttgaaagcgcactggcacgtaatggttttgaaccgtttgcagcagccggtgcactggccct
ggatctgatggacctgtctgccgaatatgaacatctgctggcagcacatcgtcagcatcaggcagttctggatgcagttagctgtggtgatgccgaag
gtgcagaacgtattatgcgtgatcatgcactggcagcaattcgtaatgcaaaagtttttgaagcagcagcaagcgcaggcgcaccgctgggtgcag
catggtcaattcgtgcagattgataaggatcctaattggtaacgaatcagacaattgacggctcgagggagtagcatagggtttgcagaatccctgct
tcgtccatttgacaggcacattatgcatcgatgataagctgtcaaacatgagcagatcctctacgccagacgcatcgtggccggcatcaccggcgcc
acaggtgcggttgctggcgcctatatcgccgacatcaccgatggggaagatcgggctcgccacttcgggctcatgagcaaatattttatctgaggtgc
ttcctcgctcactgactcgctgcacgaggcagacctcagcgctagcggagtgtatactggcttactatgttggcactgatgagggtgtcagtgaagtgc
ttcatgtggcaggagaaaaaaggctgcaccggtgcgtcagcagaatatgtgatacaggatatattccgcttcctcgctcactgactcgctacgctcgg
tcgttcgactgcggcgagcggaaatggcttacgaacggggcggagatttcctggaagatgccaggaagatacttaacagggaagtgagagggc
cgcggcaaagccgtttttccataggctccgcccccctgacaagcatcacgaaatctgacgctcaaatcagtggtggcgaaacccgacaggactat
aaagataccaggcgtttccccctggcggctccctcgtgcgctctcctgttcctgcctttcggtttaccggtgtcattccgctgttatggccgcgtttgtctcatt
ccacgcctgacactcagttccgggtaggcagttcgctccaagctggactgtatgcacgaaccccccgttcagtccgaccgctgcgccttatccggta
actatcgtcttgagtccaacccggaaagacatgcaaaagcaccactggcagcagccactggtaattgatttagaggagttagtcttgaagtcatgcg
ccggttaaggctaaactgaaaggacaagttttggtgactgcgctcctccaagccagttacctcggttcaaagagttggtagctcagagaaccttcga
aaaaccgccctgcaaggcggttttttcgttttcagagcaagagattacgcgcagaccaaaacgatctcaagaagatcatcttattaaggggtctgac
gctcagtggaacgaaaaatcaatctaaagtatatatgagtaaacttggtctgacagttaccttagaaaaactcatcgagcatcaaatgaaactgcaa
tttattcatatcaggattatcaataccatatttttgaaaaagccgtttctgtaatgaaggagaaaactcaccgaggcagttccataggatggcaagatcc
tggtatcggtctgcgattccgactcgtccaacatcaatacaacctattaatttcccctcgtcaaaaataaggttatcaagtgagaaatcaccatgagtga
cgactgaatccggtgagaatggcaaaagcttatgcatttctttccagacttgttcaacaggccagccattacgctcgtcatcaaaatcactcgcatcaa
ccaaaccgttattcattcgtgattgcgcctgagcgagacgaaatacgcgatcgctgttaaaaggacaattacaaacaggaatcgaatgcaaccgg
cgcaggaacactgccagcgcatcaacaatattttcacctgaatcaggatattcttctaatacctggaatgctgttttcccggggatcgcagtggtgagta
accatgcatcatcaggagtacggataaaatgcttgatggtcggaagaggcataaattccgtcagccagtttagtctgaccatctcatctgtaacatcat
tggcaacgctacctttgccatgtttcagaaacaactctggcgcatcgggcttcccatacaatcgatagattgtcgcacctgattgcccgacattatcgcg
agcccatttatacccatataaatcagcatccatgttggaatttaatcgcggcctggagcaagacgtttcccgttgaatatggctcataacaccccttgtat
tactgtttatgtaagcagacagttttattgttcatgatgatatatttttatcttgtgcaatgtacatcagagattttgagacacaaccaattattgaaggcctcc

ctaacggggggcctttttttgtttctggtctcccgcttaacgatcgttggctgattggatccaattgacagctagctcagtcctaggtaccattggatccaat
ag

pRC2790

ttttcgccagatatcgacgtctaagaaaccattattatcatgacattaacctataaaaataggcgtatcacgaggccgatcggtattcaattgtgcccaa
cgttgccaggtattcttgtgatttatagatccgcttataggagaggcttttcataaaattccttttaaaatcataacataaaagaatgattcacattaacgga
tccgttaactacgaaaataggcaacttattcttaaggggcaagattaatttatgttttcccgtcaccaacgacaaaatttgcgaggctctttccgaaaata
gggttgatctttgttgtcactggatgtactgtacatccatacagtaactcacaggggctggattgattatgagtaaaggagaagaacttttcactggagtt
gtcccaattcttgttgaattagatggtgatgttaatgggcacaaattttctgtcagtggagagggtgaaggtgatgcaacatacggaaaacttacccttaa
atttatttgcactactggaaaactacctgttccatggccaacacttgtcactactttcgcgtatggtcttcaatgctttgcgagatacccagatcatatgaa
acagcatgactttttcaagagtgccatgcccgaaggttatgtacaggaaagaactatatttttcaaagatgacgggaactacaagacacgtgctgaa
gtcaagtttgaaggtgatacccttgttaatagaatcgagttaaaaggtattgattttaaagaagatggaaacattcttggacacaaattggaatacaact
ataactcacacaatgtatacatcatggcagacaaacaaaagaatggaatcaaagttaacttcaaaattagacacaacattgaagatggaagcgtt
caactagcagaccattatcaacaaaatactccaattggcgatggccctgtccttttaccagacaaccattacctgtccacacaatctgccctttcgaaa
gatcccaacgaaaagagagaccacatgatccttcttgagtttgtaacagctgctgggattacacatggcatggatgaactatacaaataaggatccc
atggtacgcgtgctagaggcatcaaataaaacgaaaggctcagtcgaaagactgggcctttcgttttatctgttgtttgtcggtgaacgctctcctgagt
aggacaaatccgccgccctagacctaggggatatattccgcttcctcgctcactgactcgctacgctcggtcgttcgactgcggcgagcggaaatgg
cttacgaacggggcggagatttcctggaagatgccaggaagatacttaacagggaagtgagagggccgcggcaaagccgtttttccataggctcc
gcccccctgacaagcatcacgaaatctgacgctcaaatcagtggtggcgaaacccgacaggactataaagataccaggcgtttccccctggcgg
ctccctcgtgcgctctcctgttcctgcctttcggtttaccggtgtcattccgctgttatggccgcgtttgtctcattccacgcctgacactcagttccgggtagg
cagttcgctccaagctggactgtatgcacgaaccccccgttcagtccgaccgctgcgccttatccggtaactatcgtcttgagtccaacccggaaaga
catgcaaaagcaccactggcagcagccactggtaattgatttagaggagttagtcttgaagtcatgcgccggttaaggctaaactgaaaggacaag
ttttggtgactgcgctcctccaagccagttacctcggttcaaagagttggtagctcagagaaccttcgaaaaaccgccctgcaaggcggttttttcgtttt
cagagcaagagattacgcgcagaccaaaacgatctcaagaagatcatcttattaatcagataaaatatttctagatttcagtgcaatttatctcttcaaa
tgtagcacctgaagtcagccccatacgatataagttgttactagtgcttggattctcaccaataaaaaacgcccggcggcaaccgagcgttctgaac
aaatccagatggagttctgaggtcattactggatctatcaacaggagtccaagcgagctcgatatcaaattacgccccgccctgccactcatcgcag
tactgttgtaattcattaagcattctgccgacatggaagccatcacagacggcatgatgaacctgaatcgccagcggcatcagcaccttgtcgccttg
cgtataatatttgcccatggtgaaaacgggggcgaagaagttgtccatattggccacgtttaaatcaaaactggtgaaactcacccagggattggctg
agacgaaaaacatattctcaataaaccctttagggaaataggccaggttttcaccgtaacacgccacatcttgcgaatatatgtgtagaaactgccg
gaaatcgtcgtggtattcactccagagcgatgaaaacgtttcagtttgctcatggaaaacggtgtaacaagggtgaacactatcccatatcaccagct
caccgtctttcattgccatacggaattccggatgagcattcatcaggcgggcaagaatgtgaataaaggccggataaaacttgtgcttatttttctttacg
gtctttaaaaaggccgtaatatccagctgaacggtctggttataggtacattgagcaactgactgaaatgcctcaaaatgttctttacgatgccattggg
atatatcaacggtggtatatccagtgatttttttctccattttagcttccttagctcctgaaaatctcgataactcaaaaaatacgcccggtagtgatcttattt
cattatggtgaaagttggaacctcttacgtgccgatcaacgtctca

pRC3017

ctgtcaccggatgtgctttccggtctgatgagtccgtgaggacgaaacagcctctacaaataattttgtttaatactagagaaagaggggaaatacta
gatgacagaatcaacatcccgtcgcccggcatatgctcgcctgttggatcgtgcggtacgcattctggcggtgcgcgatcacagtgagcaagaact
gcgacgtaaactcgcggcaccgattatgggcaaaaatggcccagaagagattgatgctacggcagaagattacgagcgcgttattgcctggtgcc
atgaacatggctatctcgatgacagccgatttgttgcgcgctttatcgccagccgtagccgcaaaggttatggacctgcgcgtattcgccaggaactg
aatcagaaaggtatttcccgcgaagcgacagaaaaagcgatgcgtgaatgtgacatcgactggtgcgcactggcgcgcgatcaggcgacgcga
aaatatggcgaacctttgccaactgtcttttcagaaaaagttaagatccagcgttttctgctctatcgtggctatctgatggaagatatccaggagatttgg
cgaaattttgccgactgataactcggtaccaaattccagaaaagaggcctcccgaaagggggcctttttttcgttttggtccaatggcggcgcgccat
cgaatggtgcaaaacctttcgcggtatggcatgatagcgcccggaagagagtcaattcagggggtgaataatggacatgcctcgtattaaaccgg
gtcagcgtgttatgatggcactgcgtaaaatgattgcaagcggtgaaatcaaaagtggtgaacgtattgcagaaattccgaccgcagcagcactgg
gtgttagccgtatgccggttcgtatcgcactgcgttcactggaacaagaaggtctggttgttcgtctgggtgcacgtggttatgcagcccgtggtgttagc

agcgatcagattcgtgatgcaattgaagttcgtggtgttctggaaggttttgcagcacgtcgtctggcagaacgtggtatgaccgcagaaacccatgc
acgttttgttgtactgattgcagaaggtgaagcactgtttgcagccggtcgcctgaatggtgaagatctggatcgttatgccgcatataatcaggcatttc
atgataccctggttagcgcagcaggtaatggtgcagttgaaagcgcactggcacgtaatggttttgaaccgtttgcagcagccggtgcactggccct
ggatctgatggacctgtctgccgaatatgaacatctgctggcagcacatcgtcagcatcaggcagttctggatgcagttagctgtggtgatgccgaag
gtgcagaacgtattatgcgtgatcatgcactggcagcaattcgtaatgcaaaagtttttgaagcagcagcaagcgcaggcgcaccgctgggtgcag
catggtcaattcgtgcagattgataaggatcctaattggtaacgaatcagacaattgacggctcgagggagtagcatagggtttgcagaatccctgct
tcgtccatttgacaggcacattatgcatcgatgataagctgtcaaacatgagcagatcctctacgccagacgcatcgtggccggcatcaccggcgcc
acaggtgcggttgctggcgcctatatcgccgacatcaccgatggggaagatcgggctcgccacttcgggctcatgagcaaatattttatctgaggtgc
ttcctcgctcactgactcgctgcacgaggcagacctcagcgctagcggagtgtatactggcttactatgttggcactgatgagggtgtcagtgaagtgc
ttcatgtggcaggagaaaaaaggctgcaccggtgcgtcagcagaatatgtgatacaggatatattccgcttcctcgctcactgactcgctacgctcgg
tcgttcgactgcggcgagcggaaatggcttacgaacggggcggagatttcctggaagatgccaggaagatacttaacagggaagtgagagggc
cgcggcaaagccgtttttccataggctccgcccccctgacaagcatcacgaaatctgacgctcaaatcagtggtggcgaaacccgacaggactat
aaagataccaggcgtttccccctggcggctccctcgtgcgctctcctgttcctgcctttcggtttaccggtgtcattccgctgttatggccgcgtttgtctcatt
ccacgcctgacactcagttccgggtaggcagttcgctccaagctggactgtatgcacgaaccccccgttcagtccgaccgctgcgccttatccggta
actatcgtcttgagtccaacccggaaagacatgcaaaagcaccactggcagcagccactggtaattgatttagaggagttagtcttgaagtcatgcg
ccggttaaggctaaactgaaaggacaagttttggtgactgcgctcctccaagccagttacctcggttcaaagagttggtagctcagagaaccttcga
aaaaccgccctgcaaggcggttttttcgttttcagagcaagagattacgcgcagaccaaaacgatctcaagaagatcatcttattaaggggtctgac
gctcagtggaacgaaaaatcaatctaaagtatatatgagtaaacttggtctgacagttaccttagaaaaactcatcgagcatcaaatgaaactgcaa
tttattcatatcaggattatcaataccatatttttgaaaaagccgtttctgtaatgaaggagaaaactcaccgaggcagttccataggatggcaagatcc
tggtatcggtctgcgattccgactcgtccaacatcaatacaacctattaatttcccctcgtcaaaaataaggttatcaagtgagaaatcaccatgagtga
cgactgaatccggtgagaatggcaaaagcttatgcatttctttccagacttgttcaacaggccagccattacgctcgtcatcaaaatcactcgcatcaa
ccaaaccgttattcattcgtgattgcgcctgagcgagacgaaatacgcgatcgctgttaaaaggacaattacaaacaggaatcgaatgcaaccgg
cgcaggaacactgccagcgcatcaacaatattttcacctgaatcaggatattcttctaatacctggaatgctgttttcccggggatcgcagtggtgagta
accatgcatcatcaggagtacggataaaatgcttgatggtcggaagaggcataaattccgtcagccagtttagtctgaccatctcatctgtaacatcat
tggcaacgctacctttgccatgtttcagaaacaactctggcgcatcgggcttcccatacaatcgatagattgtcgcacctgattgcccgacattatcgcg
agcccatttatacccatataaatcagcatccatgttggaatttaatcgcggcctggagcaagacgtttcccgttgaatatggctcataacaccccttgtat
tactgtttatgtaagcagacagttttattgttcatgatgatatatttttatcttgtgcaatgtacatcagagattttgagacacaaccaattattgaaggcctcc
ctaacgggggcctttttttgtttctggtctcccgcttaacgatcgttggctgattggatccaattgacagctagctcagtcctaggtaccattggatccaat
ag

pRC3028

ctcggtaccaaattccagaaaagaggcctcccgaaagggggcctttttcgttttggtccaatggcggcgcgccatcgaatggtgcaaaacctttcg
cggtatggcatgatagcgcccggaagagagtcaattcagggggtgaataatggacatgcctcgtattaaaccgggtcagcgtgttatgatggcac
tgcgtaaaatgattgcaagcggtgaaatcaaaagtggtgaacgtattgcagaaattccgaccgcagcagcactgggtgttagccgtatgccggttcg
tatcgcactgcgttcactggaacaagaaggtctggttgttcgtctgggtgcacgtggttatgcagcccgtggtgttagcagcgatcagattcgtgatgca
attgaagttcgtggtgttctggaaggttttgcagcacgtcgtctggcagaacgtggtatgaccgcagaaacccatgcacgttttgttgtactgattgcaga
aggtgaagcactgtttgcagccggtcgcctgaatggtgaagatctggatcgttatgccgcatataatcaggcatttcatgataccctggttagcgcagc
aggtaatggtgcagttgaaagcgcactggcacgtaatggttttgaaccgtttgcagcagccggtgcactggccctggatctgatggacctgtctgccg
aatatgaacatctgctggcagcacatcgtcagcatcaggcagttctggatgcagttagctgtggtgatgccgaaggtgcagaacgtattatgcgtgat
catgcactggcagcaattcgtaatgcaaaagtttttgaagcagcagcaagcgcaggcgcaccgctgggtgcagcatggtcaattcgtgcagattga
taaggatcctaattggtaacgaatcagacaattgacggctcgagggagtagcatagggtttgcagaatccctgcttcgtccatttgacaggcacattat
gcatcgatgataagctgtcaaacatgagcagatcctctacgccagacgcatcgtggccggcatcaccggcgccacaggtgcggttgctggcgcct
atatcgccgacatcaccgatggggaagatcgggctcgccacttcgggctcatgagcaaatattttatctgaggtgcttcctcgctcactgactcgctgc
acgaggcagacctcagcgctagcggagtgtatactggcttactatgttggcactgatgagggtgtcagtgaagtgcttcatgtggcaggagaaaaa
aggctgcaccggtgcgtcagcagaatatgtgatacaggatatattccgcttcctcgctcactgactcgctacgctcggtcgttcgactgcggcgagcg
gaaatggcttacgaacggggcggagatttcctggaagatgccaggaagatacttaacagggaagtgagagggccgcggcaaagccgtttttcca

taggctccgcccccctgacaagcatcacgaaatctgacgctcaaatcagtggtggcgaaacccgacaggactataaagataccaggcgtttcccc
ctggcggctccctcgtgcgctctcctgttcctgcctttcggtttaccggtgtcattccgctgttatggccgcgtttgtctcattccacgcctgacactcagttcc
gggtaggcagttcgctccaagctggactgtatgcacgaaccccccgttcagtccgaccgctgcgccttatccggtaactatcgtcttgagtccaaccc
ggaaagacatgcaaaagcaccactggcagcagccactggtaattgatttagaggagttagtcttgaagtcatgcgccggttaaggctaaactgaa
aggacaagttttggtgactgcgctcctccaagccagttacctcggttcaaagagttggtagctcagagaaccttcgaaaaaccgccctgcaaggcg
gttttttcgttttcagagcaagagattacgcgcagaccaaaacgatctcaagaagatcatcttattaaggggtctgacgctcagtggaacgaaaaatc
aatctaaagtatatatgagtaaacttggtctgacagttaccttagaaaaactcatcgagcatcaaatgaaactgcaatttattcatatcaggattatcaat
accatatttttgaaaaagccgtttctgtaatgaaggagaaaactcaccgaggcagttccataggatggcaagatcctggtatcggtctgcgattccga
ctcgtccaacatcaatacaacctattaatttcccctcgtcaaaaataaggttatcaagtgagaaatcaccatgagtgacgactgaatccggtgagaat
ggcaaaagcttatgcatttctttccagacttgttcaacaggccagccattacgctcgtcatcaaaatcactcgcatcaaccaaaccgttattcattcgtga
ttgcgcctgagcgagacgaaatacgcgatcgctgttaaaaggacaattacaaacaggaatcgaatgcaaccggcgcaggaacactgccagcgc
atcaacaatattttcacctgaatcaggatattcttctaatacctggaatgctgttttcccggggatcgcagtggtgagtaaccatgcatcatcaggagtac
ggataaaatgcttgatggtcggaagaggcataaattccgtcagccagtttagtctgaccatctcatctgtaacatcattggcaacgctacctttgccatg
tttcagaaacaactctggcgcatcgggcttcccatacaatcgatagattgtcgcacctgattgcccgacattatcgcgagcccatttatacccatataaa
tcagcatccatgttggaatttaatcgcggcctggagcaagacgtttcccgttgaatatggctcataacaccccttgtattactgtttatgtaagcagacag
ttttattgttcatgatgatatatttttatcttgtgcaatgtacatcagagattttgagacacaaccaattattgaaggcctccctaacggggggcctttttttgttt
ctggtctcccgcttaacgatcgttggctgattggatccaattgacagctagctcagtcctaggtaccattggatccaatagctgtcaccggatgtgctttc
cggtctgatgagtccgtgaggacgaaacagcctctacaaataattttgtttaatactagagaaagaggggaaatactagatgaaaactgaactgac
cctgaatgtattacagaccatgaacgcacaggaatatgaagatatccgggctgccggaagtgatgaacgccgtgagctgactcacgccgtgatgc
gggagctggatgcgccggataactggacgatgaacggcgagtacggcagcgagtttggcggctttttccccgtccaggtccgtttcacgcctgccc
acgaacgttttcacctggcgttatgttcaccgggcgatgtctctcaggtctgggtgctggttctggtgaatgccggtggtgagccgtttgcggtggttcag
gtacagcggcggtttgcatcggaagccgtcagccattcactggcactggcggcgtcactggatacgcaggggtacagcgttaacgacatcatccat
atcctgatggcggaaggaggtcaggtatgataa

pRC3058

gtttctggtctcccgcttaacgatcgttggctgagaaaccaattgtccatattgcatcagacattgccgtcactgcgtcttttactggctcttctcgctaacca
aaccggtaaccccgcttattaaaagcattctgtaacaaagcgggaccaaagccatgacaaaaacgcgtaacaaaagtgtctataatcacggcag
aaaagtccacattgattatttgcacggcgtcacactttgctatgccatagcatttttatccataagattagcggatcctacctgacgctttttatcgcaactct
ctactgtttctccatacccgagctgtcaccggatgtgctttccggtctgatgagtccgtgaggacgaaacagcctctacaaataattttgtttaatactaga
gaaagaggggaaatactagatggcctggcttccccttaatcccattccactcaaagatcgcgtctccatgatctttctgcaatatgggcagatcgatgt
aatagatggcgcgtttgtacttatcgacaagacagggatccgcactcatattcctgttggctcggttgcctgcatcatgctggaacctggtacacgggttt
cgcatgcagctgtacgcctggctgcgcaagttggaacattgttggtatgggtgggggaagcgggcgttcgtgtttatgcttctggtcagcctggaggtg
cgcgttcagataagctgctctatcaggcaaaacttgctctggatgaagatttgcgtctgaaggtcgtacgtaaaatgtttgaacttcggtttggagaacct
gcgcctgcccggcgctccgtagagcaactcagaggtatagaaggcagtcgcgtgcgggcaacctacgcacttctggcgaagcaatacggcgtg
acatggaatggacgtcgctacgatccgaaagactgggaaaagggcgatacgatcaaccaatgcattagcgctgcaacttcctgtttatacggcgta
actgaagcggcgatacttgcagctggttatgcaccagctattgggtttgtgcatacaggaaagcctctttcctttgtttacgatattgcagacatcattaaa
tttgacactgttgtaccgaaagcttttgagatagcgcgtcgtaaccctggtgagccggaccgggaagtccgtttggcgtgcagggatatttttcgcagta
gtaaaacattagccaaattgattccgcttatagaggacgtgcttgccgctggagaaatacaaccgccggccccacctgaagatgcacagcctgttg
ccattccgcttcctgtttcactgggagatgcaggccatcggagtagctgaaatgagtatgttggtcgtggtcactgaaaatgtacctccgcgcttacgag
gcagattagccatctggttgttggaggtacgtgcaggggtatatgtaggtgatgtatccgcaaaaattcgtgaaatgatctgggaacaaatagctgga
ctggcggaagaaggcaatgtagtgatggcatgggcaacgaatacggaaacgggatttgagttccagacatttgggttaaacaggcgtaccccggt
agatttggatggtttaaggttggtgtctttttttacctgtttgagcggctcggtaccaaattccagaaaagaggcctcccgaaaggggggcctttttttcgtttt
ggtccaatggcggcgcgccatcgaatggtgcaaaacctttcgcggtatggcatgatagcgcccggaagagagtcaattcagggtggtgaatatgg
ctgaagcgcaaaatgatcccctgctgccgggatactcgtttaatgcccatctggtggcgggtttaacgccgattgaggccaacggttatctcgattttttt
atcgaccgaccgctgggaatgaaaggttatattctcaatctcaccattcgcggtcaggggtggtgaaaaatcagggacgagaatttgtttgccgac
cgggtgatattttgctgttcccgccaggagagattcatcactacggtcgtcatccggaggctcgcgaatggtatcaccagtgggtttactttcgtccgcg

cgcctactggcatgaatggcttaactggccgtcaatatttgccaatacggggttctttcgcccggatgaagcgcaccagccgcatttcagcgactttttt
gggcaaatcattaacgccgggcaaggggaagggcgctattcggagctgctggcgataaatctgcttgagcaattgttactgcggcgcatgctagcg
attaacggatcgctccatccaccgatggataatcgggtacgcgaggcttgtcagtacatcagcgatcacctggcagacagcaattttgatatcgcca
gcgtcgcacagcatgtttgcttgtcgccgtcgcgtctgtcacatcttttccgccagcagttagggattagcgtcttaagctggcgcgaggaccaacgtat
cagccaggcgaagctgcttttgagcaccacccggatgcctatcgccaccgtcggtcgcaatgttggttttgacgatcaactctatttctcgcgggtattta
aaaaatgcaccggggccagcccgagcgagttccgtgccggtttggaagaaaaagtgaatgatgtagccgtcaagttgtcatgataagatcctattc
cagcgggattaaagaggagcgattaagcatggttactatcaatacggaatctgctttaacgccacgttctttgcgggatacgcggcgtatgaatatgtt
tgtttcggtagctgctgcggtcgcaggattgttatttggtcttgatatcggcgtaatcgccggagcgttgccgttcattaccgatcactttgtgctgaccagtc
gtttgcaggaatgggtggttagtagcatgatgctcggtgcagcaattggtgcgctgtttaatggttggctgtcgttccgcctggggcgtaaatacagcct
gatggcgggggccatcctgtttgtactcggttctatagggtccgcttttgcgaccagcgtagagatgttaatcgccgctcgtgtggtgctgggcattgctg
tcgggatcgcgtcttacaccgctcctctgtatctttctgaaatggcaagtgaaaacgttcgcggtaagatgatcagtatgtaccagttgatggtcacactc
ggcatcgtgctggcgtttttatccgatacagcgttcagttatagcggtaactggcgcgcaatgttgggggttcttgctttaccagcagttctgctgattattct
ggtagtatttctgccaaatagcccgcgctggctggcggaaaaggggcgtcatattgaggcggaagaagtattgcgtatgctgcgcgatacgtcgga
aaaagcgcgagaagaactcaacgaaattcgtgaaagcctgaagttaaaacagggcggttgggcactgtttaagatcaaccgtaacgtccgtcgtg
ctgtgtttctcggtatgttgttgcaggcgatgcagcagtttaccggtatgaacatcatcatgtactacgcgccgcgtatcttcaaaatggcgggctttacg
accacagaacaacagatgattgcgactctggtcgtagggctgacctttatgttcgccacctttattgcggtgtttacggtagataaagcagggcgtaaa
ccggctctgaaaattggtttcagcgtgatggcgttaggcactctggtgctgggctattgcctgatgcagtttgataacggtacggcttccagtggcttgtc
ctggctctctgttggcatgacgatgatgtgtattgccggttatgcgatgagcgccgcgccagtggtgtggatcctgtgctctgaaattcagccgctgaaa
tgccgcgatttcggtattacctgttcgaccaccacgaactgggtgtcgaatatgattatcggcgcgaccttcctgacactgcttgatagcattggcgctg
ccggtacgttctggctctacactgcgctgaacattgcgtttgtgggcattactttctggctcattccggaaaccaaaaatgtcacgctggaacatatcga
acgcaaactgatggcaggcgagaagttgagaaatatcggcgtctgataaggatcctaattggtaacgaatcagacaattgacggctcgagggagt
agcatagggtttgcagaatccctgcttcgtccatttgacaggcacattatgcatcgatgataagctgtcaaacatgagcagatcctctacgccggacg
catcgtggccggcatcaccggcgccacaggtgcggttgctggcgcctatatcgccgacatcaccgatggggaagatcgggctcgccacttcgggc
tcatgagcaaatattttatctgaggtgcttcctcgctcactgactcgctgcacgaggcagacctcagcgctagcggagtgtatactggcttactatgttgg
cactgatgagggtgtcagtgaagtgcttcatgtggcaggagaaaaaaggctgcaccggtgcgtcagcagaatatgtgatacaggatatattccgctt
cctcgctcactgactcgctacgctcggtcgttcgactgcggcgagcggaaatggcttacgaacggggcggagatttcctggaagatgccaggaag
atacttaacagggaagtgagagggccgcggcaaagccgtttttccataggctccgcccccctgacaagcatcacgaaatctgacgctcaaatcag
tggtggcgaaacccgacaggactataaagataccaggcgtttccccctggcggctccctcgtgcgctctcctgttcctgcctttcggtttaccggtgtca
ttccgctgttatggccgcgtttgtctcattccacgcctgacactcagttccgggtaggcagttcgctccaagctggactgtatgcacgaaccccccgttca
gtccgaccgctgcgccttatccggtaactatcgtcttgagtccaacccggaaagacatgcaaaagcaccactggcagcagccactggtaattgattt
agaggagttagtcttgaagtcatgcgccggttaaggctaaactgaaaggacaagttttggtgactgcgctcctccaagccagttacctcggttcaaag
agttggtagctcagagaaccttcgaaaaaccgccctgcaaggcggttttttcgttttcagagcaagagattacgcgcagaccaaaacgatctcaag
aagatcatcttattaaggggtctgacgctcagtggaacgaaaaatcgagtaaacttggtctgacagtgatcggcacgtaagaggttccaactttcacc
ataatgaaataagatcactaccgggcgtattttttgagttgtcgagattttcaggagctaaggaagctaaaatggagaaaaaaatcactggatatacc
accgttgatatatcccaatggcatcgtaaagaacattttgaggcatttcagtcagttgctcaatgtacctataaccagaccgttcagctggatattacgg
cctttttaaagaccgtaaagaaaaataagcacaagttttatccggcctttattcacattcttgcccgcctgatgaatgctcatccggaattacgtatggca
atgaaagacggtgagctggtgatatgggatagtgttcacccttgttacaccgttttccatgagcaaactgaaacgttttcatcgctctggagtgaatacc
acgacgatttccggcagtttctacacatatattcgcaagatgtggcgtgttacggtgaaaacctggcctatttccctaaagggtttattgagaatatgtttttt
cgtctcagccaatccctgggtgagtttcaccagttttgatttaaacgtggccaatatggacaacttcttcgcccccgttttcaccatgggcaaatattatac
gcaaggcgacaaggtgctgatgccgctggcgattcaggttcatcatgccgtttgtgatggcttccatgtcggcagatgcttaatgaatacaacagtact
gcgatgagtggcagggcggggcgtaaagagtttgtagaaacgcaaaaaggccatccgtcaggatggccttctgcttaatttgatgcctggcagttta
tggcgggcgtcctgcccgccaccctccgggccgttgcttcgcaacgttcaaatccgctcccggcggatttgtcctactcaggagagcgttcaccgac
aaacaacagataaaacgaaaggcccagtctttcgactgagcctttcgttttatttgatgcctggcagttccctactctcgcatggggagaccccacact
accatcggcgctac

### S2 Table. Python scripts

**Script 1**

**Searches through DNA sequences to identify spacer sequences and records matching results along with relevant metadata:**

```
import os

import csv


def reverse_complement(sequence):

    complement = str.maketrans("acgt", "tgca")

    return sequence.translate(complement)[::-1]


def hamming_distance(seq1, seq2):

    """Calculate the Hamming distance between two sequences."""

    return sum(el1 != el2 for el1, el2 in zip(seq1, seq2))


def allowed_mismatches(sequence, query, max_mismatches):

    """Find all positions where the query matches the sequence with allowed mismatches."""

    matches = []

    query_len = len(query)


    for i in range(len(sequence) - query_len + 1):

        subseq = sequence[i:i + query_len]

        mismatches = hamming_distance(subseq, query)

        if mismatches <= max_mismatches:

            matches.append((i, subseq, mismatches))  # Include mismatches for concatenation

    return matches


def process_fastq(input_fastq, output_csv, max_mismatches):

    search_sequence = "tatgtttagagtgttccccgcgccagcggggataaacc".lower()


    results_found = False

    all_results = []  # List to store all results before writing to the CSV
```

```
try:
    with open(input_fastq, 'r') as fastq_file:
        while True:
            header = fastq_file.readline().strip()  # Read and store the sequence header
            sequence = fastq_file.readline().strip().lower()
            fastq_file.readline()  # Skip the plus line
            fastq_file.readline()  # Skip the quality score line

            if not sequence:
                break

            # Search in forward strand
            forward_matches = allowed_mismatches(sequence, search_sequence, max_mismatches)
            for position, mismatched_seq, mismatches in forward_matches:
                following_33bp = sequence[position + len(search_sequence):position + len(search_sequence) + 33]
                concatenated = mismatched_seq + following_33bp
                all_results.append([concatenated, mismatches, "Forward", following_33bp, header])
                results_found = True

            # Search in reverse complement
            rev_comp_sequence = reverse_complement(sequence)
            reverse_matches = allowed_mismatches(rev_comp_sequence, search_sequence, max_mismatches)
            for position, mismatched_seq, mismatches in reverse_matches:
                following_33bp = rev_comp_sequence[position + len(search_sequence):position + len(search_sequence) + 33]
                concatenated = mismatched_seq + following_33bp
                all_results.append([concatenated, mismatches, "Reverse", following_33bp, header])
                results_found = True

    if results_found:
        # Sort the results: forward strand at the top, reverse strand at the bottom
```

```
            all_results.sort(key=lambda x: x[2])  # Sort by the 'Strand' column ("Forward" before "Reverse")

            # Reassign the count after sorting
            for count, result in enumerate(all_results, start=1):
                result.insert(0, count)  # Insert the count at the beginning of each row

            # Write the sorted and updated results to the CSV file
            try:
                with open(output_csv, 'w', newline='') as csv_file:
                    writer = csv.writer(csv_file)
                    # Updated headers based on the new requirements
                    writer.writerow(["Count", "Leader-Repeat-Spacer Unit", "Number of Mismatches", "Fastq
Strand", "Spacer", "Metadata (Fastq Header)"])
                    writer.writerows(all_results)

                print(f"Results successfully written to '{output_csv}'.")

            except IOError as e:
                print(f"Error: Unable to write to the file '{output_csv}'. {e}")
        else:
            print(f"No matches found for '{search_sequence}' in '{input_fastq}'.")

    except FileNotFoundError:
        print(f"Error: The file '{input_fastq}' was not found.")
    except IOError as e:
        print(f"Error: There was an issue with reading the file '{input_fastq}'. {e}")
    except Exception as e:
        print(f"An unexpected error occurred: {e}")

def process_directory(directory_path, max_mismatches):
    if not os.path.isdir(directory_path):
        print(f"Error: The directory '{directory_path}' does not exist.")
        return
```

```
    # Loop through all files in the directory
    for filename in os.listdir(directory_path):
        if filename.endswith(".fastq"):
            input_fastq = os.path.join(directory_path, filename)
            output_csv = os.path.join(directory_path, filename.replace(".fastq", ".csv"))
            print(f"Processing file: {input_fastq}")
            process_fastq(input_fastq, output_csv, max_mismatches)

if __name__ == "__main__":
    try:
        directory_path = input("Enter the directory path containing FASTQ files: ").strip()
        max_mismatches = int(input("Enter the number of allowed mismatches: ").strip())

        if not directory_path or max_mismatches < 0:
            print("Error: A valid directory path and a non-negative number of mismatches must be provided.")
            exit()

        process_directory(directory_path, max_mismatches)

    except ValueError:
        print("Error: Invalid number of mismatches provided. It must be an integer.")
    except Exception as e:
        print(f"An unexpected error occurred: {e}")
```

**Script 2**

**Converts csv files generated from script 1 into FASTA format:**

```
import os
import pandas as pd

def csv_to_fasta():
    # Prompt for the directory path containing the CSV files
    directory = input("Enter the directory path containing the CSV files: ")

    # Iterate through each file in the directory
    for filename in os.listdir(directory):
        # Process only .csv files
        if filename.endswith('.csv'):
            # Define the full path to the input file
            csv_file = os.path.join(directory, filename)

            # Create the corresponding output FASTA file name
            fasta_file = os.path.join(directory, filename.rsplit('.', 1)[0] + '.fasta')

            # Read the CSV file
            df = pd.read_csv(csv_file)

            # Clean column names (strip whitespace and convert to lowercase)
            df.columns = df.columns.str.strip().str.lower()

            # Verify columns
            print(f"Processing {filename} with columns: {df.columns}")

            # Extract base name of CSV file for FASTA headers
            base_name = filename.rsplit('.', 1)[0]

            # Open the FASTA file for writing
            with open(fasta_file, 'w') as fasta_out:
```

```
        # Iterate through each row in the DataFrame
        for index, row in df.iterrows():
            # Get the sequence and count
            sequence = row['spacer']  # Updated to match the correct column name
            count = row['count']  # Assuming the count is in the 'count' column

            # Create the FASTA header without .csv in the name
            header = f">{base_name}_{count}"

            # Write the header and sequence to the FASTA file
            fasta_out.write(f"{header}\n{sequence}\n")

    print(f"Finished writing to {fasta_file}")

# Run the function
csv_to_fasta()
```

**Script 3**

**Concatenates multiple FASTA files:**

```
import os
def concatenate_fasta_files(directory, output_file):
    fasta_files = [f for f in os.listdir(directory) if f.endswith('.fasta')]
    if not fasta_files:
        print("No FASTA files found in the directory.")
        return

    with open(output_file, 'w') as outfile:
        for fasta_file in fasta_files:
            with open(os.path.join(directory, fasta_file), 'r') as infile:
                outfile.write(infile.read())

    print(f"Concatenation complete! Output saved as: {output_file}")

def main():
    directory = input("Please enter the directory path containing the fasta files: ").strip()
    if not os.path.isdir(directory):
        print("Invalid directory path.")
        return

    output_file = input("Please enter the name for the output fasta file (e.g., output.fasta): ").strip()

    if not output_file.endswith('.fasta'):
        output_file += '.fasta'
    concatenate_fasta_files(directory, output_file)

if __name__ == "__main__":
    main()
```

**Script 4**

**Removes duplicate spacer sequences and orders the remaining spacers by their frequency:**

```
from Bio import SeqIO

def remove_duplicates_with_count_sorted(input_fasta, output_fasta):
    sequence_counts = {}
    sequence_records = {}

    for record in SeqIO.parse(input_fasta, "fasta"):
        seq_str = str(record.seq)
        if seq_str in sequence_counts:
            sequence_counts[seq_str] += 1
        else:
            sequence_counts[seq_str] = 1
            sequence_records[seq_str] = record

    sorted_sequences = sorted(sequence_counts.items(), key=lambda x: x[1], reverse=True)

    records_to_write = []
    for seq_str, count in sorted_sequences:
        record = sequence_records[seq_str]
        record.id += f"_count{count}"
        record.description = ""
        records_to_write.append(record)

    SeqIO.write(records_to_write, output_fasta, "fasta")
    print(f"Processed sequences. Unique sequences sorted by count written to {output_fasta}")

input_fasta = input("Enter the input FASTA file name: ")
output_fasta = input("Enter the output FASTA file name: ")

remove_duplicates_with_count_sorted(input_fasta, output_fasta)
```

### Script 5

**Removes any spacer sequence that has an 80% or greater sequence identity to the parental spacer in the RC5311 LacZ reporter CRISPR array:**

```
from Bio import SeqIO

from Bio import pairwise2


input_fasta = input("Enter the input FASTA file name (including path if necessary): ").strip()

output_fasta = input("Enter the output FASTA file name (including path if necessary): ").strip()


reference_seq = "gctttcgcagacgcgcggcgatacgctcacgca"

threshold_identity = 80  # Percent identity threshold


def calculate_percent_identity(seq1, seq2):

    """Calculate the percent identity between two sequences."""

    alignment = pairwise2.align.globalxx(seq1, seq2, one_alignment_only=True, score_only=True)

    percent_identity = (alignment / max(len(seq1), len(seq2))) * 100

    return percent_identity


filtered_sequences = []

try:

    for record in SeqIO.parse(input_fasta, "fasta"):

        identity = calculate_percent_identity(str(record.seq), reference_seq)

        if identity < threshold_identity:

            filtered_sequences.append(record)


    SeqIO.write(filtered_sequences, output_fasta, "fasta")

    print(f"Filtered sequences have been saved to {output_fasta}")

except FileNotFoundError:

    print(f"Error: File {input_fasta} not found. Please check the file name and path.")

except Exception as e:

    print(f"An error occurred: {e}")
```

**Script 6**

**Automates a BLASTN workflow that compares the spacer sequences to a reference database, extracts PAM sequences and annotates the data with match counts and frequency adjustment:**

```
import subprocess

import os

import csv

from collections import Counter


def reverse_complement(seq):

    complement = str.maketrans('ACGTacgt', 'TGCAtgca')

    return seq.translate(complement)[::-1]


def extract_pam(sequence, sstart, send, sstrand):

    if sstrand == 'plus':

        pam_position = sstart - 4

        if pam_position >= 0:

            return sequence[pam_position:sstart-1]

    elif sstrand == 'minus':

        pam_position = sstart

        if pam_position + 2 < len(sequence):

            pam_sequence = sequence[pam_position:pam_position + 3]

            return reverse_complement(pam_sequence)

    return "N/A"


def read_sequences(file_path):

    sequences = {}

    print(f"Reading sequences from {file_path}...")

    try:

        with open(file_path, 'r') as f:

            seq_id = None

            for line in f:

                line = line.strip()

                if line.startswith('>'):

                    seq_id = line[1:]
```

```
                sequences[seq_id] = ''
            elif seq_id:
                sequences[seq_id] += line
        print(f"Loaded {len(sequences)} sequences.")
    except FileNotFoundError:
        print(f"Error: File {file_path} not found.")
    except PermissionError:
        print(f"Error: Permission denied for file {file_path}.")
    return sequences

def perform_blast(query_fasta, db_name):
    temp_output_file = f"{os.path.splitext(query_fasta)[0]}_temp_results.txt"
    blastn_command = [
        "blastn",
        "-query", query_fasta,
        "-db", db_name,
        "-out", temp_output_file,
        "-outfmt", "6 qseqid sseqid pident length mismatch gapopen qstart qend sstart send evalue bitscore
sstrand"
        ]
    print(f"Running BLAST command: {' '.join(blastn_command)}")
    try:
        subprocess.run(blastn_command, check=True)
        print("BLAST search completed.")
    except subprocess.CalledProcessError as e:
        print(f"An error occurred during BLAST search: {e}")
        return None
    return temp_output_file

def process_blast_results(temp_file, query_sequences, db_sequences):
    jb028_results = []
    p1656_results = []
    aligned_to_p1656 = set()
    with open(temp_file, 'r') as temp_file:
```

```
        lines = temp_file.readlines()
    for line in lines:
        fields = line.strip().split('\t')
        if len(fields) >= 13:
            try:
                pident = float(fields[2])
                length = int(fields[3])
                if pident == 100.0 and length == 33:
                    qseqid = fields[0]
                    sseqid = fields[1]
                    sstart = int(fields[8])
                    send = int(fields[9])
                    sstrand = fields[12]
                    query_sequence = query_sequences.get(qseqid, 'Unknown')
                    db_sequence = db_sequences.get(sseqid, '')
                    pam_sequence = extract_pam(db_sequence, sstart, send, sstrand)
                    if query_sequence:
                        pam_sequence += query_sequence[0]
                    pam_sequence = pam_sequence.upper()
                    if "JB028" in sseqid:
                        jb028_results.append([qseqid, query_sequence] + fields[1:3] + fields[3:8] + [sstart, send] +
fields[10:] + [pam_sequence])
                    elif "p1656" in sseqid:
                        p1656_results.append([qseqid, query_sequence] + fields[1:3] + fields[3:8] + [sstart, send]
+ fields[10:] + [pam_sequence])
                        aligned_to_p1656.add(qseqid)
            except ValueError:
                pass
    return jb028_results, p1656_results, aligned_to_p1656


def calculate_match_counts_and_frequency(results):
    sequences = [row[1] for row in results]
    sequence_counts = Counter(sequences)
    for row in results:
```

```
        sequence = row[1]
        match_count = sequence_counts[sequence]
        frequency_adjustment = 1 / match_count
        row.append(match_count)
        row.append(frequency_adjustment)

def write_output(jb028_results, p1656_results, aligned_to_p1656, output_file):
    with open(output_file, 'w', newline='') as csvfile:
        csvwriter = csv.writer(csvfile)
        column_headings = [
            "Count", "Spacer ID", "Sequence", "Database", "pident", "length", "mismatch", "gapopen",
            "qstart", "qend", "sstart", "send", "evalue", "bitscore", "sstrand", "PAM",
            "Spacer Match Counts", "Frequency Adjustment"
        ]
        csvwriter.writerow(column_headings)
        count = 1
        calculate_match_counts_and_frequency(p1656_results)
        calculate_match_counts_and_frequency(jb028_results)
        for row in p1656_results:
            csvwriter.writerow([count] + row)
            count += 1
        for row in jb028_results:
            if row[0] not in aligned_to_p1656:
                csvwriter.writerow([count] + row)
                count += 1
    print(f"Results saved to {output_file}.")

def main():
    print("Script started.")
    db_name = input("Enter the base name of the BLAST database (e.g., JB028-p1656): ")
    db_fasta_file = f"{db_name}.fasta"
    if not os.path.isfile(db_fasta_file):
        print(f"Error: The file {db_fasta_file} does not exist.")
```

```
        return
    db_sequences = read_sequences(db_fasta_file)
    directory = input("Enter the directory containing .fasta files: ")
    if not os.path.isdir(directory):
        print(f"Error: The directory {directory} does not exist.")
        return
    fasta_files = [f for f in os.listdir(directory) if f.lower().endswith('.fasta')]
    if not fasta_files:
        print("No .fasta files found in the directory.")
        return
    for fasta_file in fasta_files:
        fasta_path = os.path.join(directory, fasta_file)
        if not os.path.isfile(fasta_path):
            print(f"Skipping invalid file: {fasta_path}")
            continue
        query_sequences = read_sequences(fasta_path)
        temp_results = perform_blast(fasta_path, db_name)
        if temp_results is None:
            continue
        jb028_results, p1656_results, aligned_to_p1656 = process_blast_results(temp_results,
query_sequences, db_sequences)
        output_file = f"{os.path.splitext(fasta_file)[0]}-blast-{db_name}.csv"
        write_output(jb028_results, p1656_results, aligned_to_p1656, output_file)
        if os.path.exists(temp_results):
            os.remove(temp_results)
    print("Script completed successfully.")


if __name__ == "__main__":
    main()
```

**S3 Table. Distribution of cell-length phenotypes in representative micrographs**

Representative .czi micrographs were scored using a custom Python-based image-analysis workflow. Cell lengths were calibrated from embedded image metadata and the 50 $\mu$m scale bar, and segmented objects were assigned to predefined length-based categories: normal rods (<4 $\mu$m), mildly elongated (4 to <8 $\mu$m), filaments (8 to <15 $\mu$m), or extreme filaments (≥15 $\mu$m). Septating, overlapping, clustered, or poorly resolved cells were assigned to an ambiguous category. Values represent approximate percentages within the fields shown.

| Phenotype category | Uninduced | Cas1–Cas2 | Cas1 | Cas2 | dCas1–Cas2 | Cas1–Cas2 IHFα | Cas1–Cas2 lexA3 |
|---|---|---|---|---|---|---|---|
| Normal rods (<4 $\mu$m) | ~90–95% | ~55–60% | ~85–90% | ~90–95% | ~85–90% | ~60–65% | ~85–90% |
| Mildly elongated (4 to <8 $\mu$m) | ~5–8% | ~20–25% | ~8–12% | ~5–8% | ~8–10% | ~20–25% | ~8–10% |
| Filaments (8 to <15 $\mu$m) | ~0–1% | ~10–15% | ~0–2% | ~0–1% | ~0–2% | ~8–12% | ~0–2% |
| Extreme filaments (≥15 $\mu$m) | 0% | ~1–3% | 0% | 0% | 0% | ~1–2% | 0% |
| Ambiguous / dividing / clustered | <5% | <5% | <5% | <5% | <5% | <5% | <5% |
